# Genome size evolution in the diverse insect order Trichoptera

**DOI:** 10.1101/2021.05.10.443368

**Authors:** Jacqueline Heckenhauer, Paul B. Frandsen, John S. Sproul, Zheng Li, Juraj Paule, Amanda M. Larracuente, Peter J. Maughan, Michael S. Barker, Julio V. Schneider, Russell J. Stewart, Steffen U. Pauls

## Abstract

**Background:** Genome size is implicated in form, function, and ecological success of a species. Two principally different mechanisms are proposed as major drivers of eukaryotic genome evolution and diversity: Polyploidy (i.e., whole genome duplication: WGD) or smaller duplication events and bursts in the activity of repetitive elements (RE). Here, we generated *de novo* genome assemblies of 17 caddisflies covering all major lineages of Trichoptera. Using these and previously sequenced genomes, we use caddisflies as a model for understanding genome size evolution in diverse insect lineages.

**Results:** We detect a ~14-fold variation in genome size across the order Trichoptera. We find strong evidence that repetitive element (RE) expansions, particularly those of transposable elements (TEs), are important drivers of large caddisfly genome sizes. Using an innovative method to examine TEs associated with universal single copy orthologs (i.e., BUSCO genes), we find that TE expansions have a major impact on protein-coding gene regions, with TE-gene associations showing a linear relationship with increasing genome size. Intriguingly, we find that expanded genomes preferentially evolved in caddisfly clades with a higher ecological diversity (i.e., various feeding modes, diversification in variable, less stable environments).

**Conclusion:** Our findings provide a platform to test hypotheses about the potential evolutionary roles of TE activity and TE-gene associations, particularly in groups with high species, ecological, and functional diversities.

## Background

Genome size is a fundamental biological character. Studying its evolution may potentially lead to a better understanding of the origin and underlying processes of the myriad forms and functions of plants and animals. This diversification process remain at the core of much biological research. Given their high species, ecological and functional diversities, insects are excellent models for such research. To date 1,345 insect genome size estimates have been published (Gregory, 2005: Animal Genome Size Database: http://www.genomesize.com, last accessed 2021-04-30) ranging 240-fold from 69 Mbp in chironomid midges [1] to 16.5p Gbp in the mountain grasshopper *Podisma pedestris* [2]. Genome size variation relates poorly to the number of coding genes or the complexity of the organism (C-value enigma, [3],[4],[5],[6]) and evolutionary drivers of genome size variation remain a topic of ongoing debate (e.g. [7], [8], [9], [10]). Two principally different mechanisms are proposed as primary drivers of eukaryotic genome size evolution: Whole genome duplication (WGD, i.e., polyploidy) or smaller duplication events and expansion of repetitive elements (REs, [5]). While WGD is ubiquitous in plant evolution, it has been regarded as the exception in animals [11], [12]. However, ancient WGD has been hypothesized to be an important driver of evolution of mollusks (e.g. [13]) amphibians (e.g. [14], [15], fish (e.g. [16], [17], [18]) and arthropods (e.g. [19], [20], [21]), including multiple putative ancient large-scale gene duplications within Trichoptera [22].

RE expansion is an important driver of genome size variation in many eukaryotic genomes [23], [24]. The two major categories of REs are tandem repeats (e.g., satellite DNA) and mobile transposable elements (TEs). TEs are classified into class I [retrotransposons: endogenous retroviruses (ERVs), related long terminal repeat (LTR) and non-LTR retrotransposons: SINEs (Short Interspersed Nuclear Elements), LINEs (Long Interspersed Nuclear Elements)] and class II elements (DNA transposons, [25]). In insects, the known genomic proportion of TEs ranges from 1% in the antarctic midge *Belgica antarctica* [26] to 65% in the migratory locust *Locusta migratoria* [27]. Broad-scale analysis of TE abundance in insects suggests that some order-specific signatures are present, however, major shifts in TE abundance are also common at shallow taxonomic levels [28], [29], including in Trichoptera [30]. The movement and proliferation of REs can have deleterious consequences on gene function and genome stability [31], [32], [33], [34], [35]. Moreover, repeat content and abundance can turn over rapidly even over short evolutionary time scales (reviewed in [36]). This rapid evolution has consequences for genome evolution and speciation, e.g., repeat divergence causes genetic incompatibilities between even closely related species [37]. However, TEs can also be sources of genomic innovation with selective advantages for the host [38], [39], [40], [41], [42], [43] and they can contribute to global changes in gene regulatory networks [44], [45], [46]. Investigating RE dynamics in diverse clades provides a powerful lens for understanding their roles in genome function and evolution. Broadly studying of RE dynamics in species-rich groups with wide variation in RE activity is an important step towards efficiently identifying study systems at finer taxonomical scales (natural populations, species complexes, or recently diverged species) that are ideally suited to advance our understanding of molecular and evolutionary mechanisms underlying genome evolution. In addition, by taking this biodiversity genomics approach, we can develop new model systems and eventually better understand links between environmental factors, genome size evolution, adaptation, and speciation (see [47]).

With more than 16,500 species, caddisflies (Trichoptera) are among the most diverse of all aquatic insects [48]. Their species richness is reflective of their ecological diversity, including, e.g. microhabitat specialization, a full array of feeding modes, and diverse use of underwater silk secretions [49], [50]. An initial comparison of six caddisfly species found wide genome size variation in Trichoptera (ranging from 230 Mbp to 1.4 Gbp). In that study, we hypothesized that the observed variation was correlated with caddisfly phylogeny and that TEs contributed to a suborder-specific increase of genome size [30].

Here, we present a multi-faceted analysis to investigate genome size evolution in the order Trichoptera, as an example for highly diversified non-model organisms. Specifically, we (i) estimated genome size for species across the order to explore phylogenetic patterns in the distribution of genome size variation in Trichoptera and (ii) generated 17 new Trichoptera genomes to analyze, in conjunction with 9 existing genomes, the causes (WGD, TE expansions) of genome size variation in the evolution of caddisflies. Studying the genomic diversity of this highly diversified insect order adds new insights into drivers of genome size evolution with potential to shed light on how genome size is linked to form, function, and ecology.

### Data Description

#### Genomic resources

Here, we combined long- and short-read sequencing technologies to generate 17 new *de novo* genome assemblies across a wide taxonomic range, covering all major lineages of Trichoptera. Details on sequencing coverage and assembly strategies are given in DataS1_Sup.2, DataS1_Sup.3, and supplementary note 3. To assess quality, we calculated assembly statistics with QUAST v5.0.2 [51], examined gene completeness with BUSCO v3.0.2 [52], [53] and screened for potential contamination with taxon-annotated GC-coverage (TAGC) plots using BlobTools v1.0 ([94], supplementary Figs. S31-S47). The new genomes are of comparable or better quality than other Trichoptera genomes previously reported in terms of BUSCO completeness and contiguity (Table 1). This study increases the number of assemblies in this order from nine to 26, nearly tripling the number of available caddisfly genomes and thus providing a valuable resource for studying genomic diversity across this ecologically diverse insect order. The annotation of these genomes predicted 6,413 to 12,927 proteins (Datas1_Sup.2). Most of the annotated proteins (94.4% - 98.8%) showed significant sequence similarity to entries in the NCBI nr database. GO Distributions were similar to previously annotated caddisfly genomes, i.e. the major biological processes were cellular and metabolic processes. Catalytic activity was the largest subcategory in molecular function and the cell membrane subcategories were the largest cellular component (supplementary Figs. S1-S30). This project has been deposited at NCBI under BioProject ID: PRJNA558902. For accession numbers of individual assemblies see Table 1.

**Table 1.** Comparison of assembly and annotation statistics of all available Trichoptera Genomes. *Assemblies produced in this study. **N_Arthropoda_=2442

| Species | Abbreviation | Accession number | Length (bp) | N50 (kbp) | No. of scaffolds/contigs | BUSCOS** |
| --- | --- | --- | --- | --- | --- | --- |
| *Agapetus fuscipes* * | AF | JAGTXP000000000 | 552,637,417 | 2.8 | 291,536 | C:39.7% [S:39.2%,D:0.5%],F:35.8%,M:24.5%,n:2442 |
| *Agraylea sexmaculata* * | AS | JAGTTH000000000 | 196,044,125 | 86 | 7,050 | C:94.3% [S:89.8%,D:4.5%],F:2.3%,M:3.4%,n:2442 |
| *Agrypnia vestita* [30] | AV | JADDOH000000000 | 1,352,945,503 | 111.8 | 25,541 | C:87.3% [S:79.0%,D:8.3%],F:5.5%,M:7.2%,n:2442 |
| *Drusus annulatus* * | DA | JAGWCC000000000 | 727,941,535 | 1,043.7 | 2,401 | C:93.5% [S:93.0%,D:0.5%],F:3.3%,M:3.2%,n:2442 |
| *Glossosoma conforme* * | GC1 | JAGTXR000000000 | 568,249,599 | 2,212.1 | 653 | C:88.9% [S:88.0%,D:0.9%],F:2.9%,M:8.2%,n:2442 |
| *Glossosoma conforme* [124] | GC2 | GCA_003347265.1 | 604,293,666 | 17.1 | 119,821 | C:74.2% [S:73.5%,D:0.7%],F:17.9%,M:7.9%,n:2442 |
| *Glyptotaelius pellucidula* [125] | GP | Glyptotaelius_pellucidus_k51_scaffolds | 623,431,006 | 1.6 | 461,749 | C:15.7% [S:15.7%,D:0.0%],F:31.4%,M:52.9%,n:2442 |
| *Halesus radiatus* * | HR | JAHDVE000000000 | 973,356,502 | 125.2 | 12,484 | C:85.7% [S:83.3%,D:2.4%],F:4.9%,M:9.4%,n:2442 |
| *Himalopsyche phryganea* * | HP | JAGVSL000000000 | 633,785,554 | 4,634 | 710 | C:95.5% [S:94.8%,D:0.7%],F:2.3%,M:2.2%,n:2442 |
| *Hesperophylax magnus* [30] | HM | JADDOG000000000 | 1,275,967,528 | 768.2 | 6,877 | C:92.5% [S:85.9%,D:6.6%],F:2.7%,M:4.8%,n:2442 |
| *Hydropsyche tenuis* [57] | HT | GCA_009617725.1 | 229,663,394 | 2,190.1 | 403 | C:94.4% [S:93.5%,D:0.9%],F:3.2%,M:2.4%,n:2442 |
| *Lepidostoma basale* * | LB | JAGTTH000000000 | 769,208,668 | 1,052 | 1,621 | C:93.9%[S:92.8%,D:1.1%],F:3.1%,M:3.0%,n:2442 |
| *Limnephilus lunatus* [69] | LL | GCA_000648945.2 | 1,369,180,260 | 69.1 | 58,718 | C:79.3%[S:74.6%,D:4.7%],F:11.7%,M:9.0%,n:2442 |
| *Micrasema longulum* * | ML2 | JAGXCS000000000 | 668,600,304 | 2.5 | 368,330 | C:78.6%[S:77.7%,D:0.9%],F:5.9%,M:15.5%,n:2442 |
| *Micrasema longulum* * | ML1 | JAGVSM000000000 | 585,245,295 | 170.5 | 5,451 | C:38.2%[S:38.0%,D:0.2%],F:31.6%,M:30.2%,n:2442 |
| *Micrasema minimum* * | MM | JAGVSQ000000000 | 329,257,313 | 69.5 | 7,561 | C:55.4%[S:55.2%,D:0.2%],F:11.7%,M:32.9%,n:2442 |
| *Micropterna sequax* * | MS | JAGUCF000000000 | 778,692,278 | 7.9 | 144,286 | C:43.4%[S:42.0%,D:1.4%],F:25.5%,M:31.1%,n:2442 |
| *Odontocerum albicorne* * | OA | JAGTXQ000000000 | 1,305,984,461 | 266.4 | 9,303 | C:91.1%[S:90.1%,D:1.0%],F:4.8%,M:4.1%,n:2442 |
| *Parapysche elsis* * | PE | JAGVSN000000000 | 282,185,525 | 5,591.7 | 159 | C:95.0%[S:94.5%,D:0.5%],F:2.4%,M:2.6%,n:2442 |
| *Philopotamus ludificatus* * | PL | JAGXCT000000000 | 360,300,449 | 67.5 | 37,274 | C:91.0%[S:89.4%,D:1.6%],F:5.9%,M:3.1%,n:2442 |
| *Plectrocnemia conspersa* [57] | PC | GCA_009617715.1 | 396,695,105 | 869 | 1,614 | C:93.5%[S:92.6%,D:0.9%],F:4.3%,M:2.2%,n:2442 |
| *Rhyacophila brunnea* * | RB | JAGYXB000000000 | 1,086,872,538 | 1,030.6 | 2,125 | C:94.5%[S:91.6%,D:2.9%],F:2.8%,M:2.7%,n:2442 |
| *Rhyacophila evoluta* * | RE2 | JAGVSQ000000000 | 565,830,460 | 9.9 | 114,057 | C:71.7%[S:71.3%,D:0.4%],F:20.5%,M:7.8%,n:2442 |
| *Rhyacophila evoluta* * | RE1 | JAGVSO000000000 | 562,550,625 | 9.7 | 111,706 | C:71.7%[S:71.4%,D:0.3%],F:20.6%,M:7.7%,n:2442 |
| *Sericostoma sp.*[126] | SS | GCA_003003475.1 | 1,015,727,762 | 3.2 | 561,698 | C:26.4%[S:26.4%,D:0.0%],F:34.4%,M:39.2%,n:2442 |
| *Stenopsyche tienhuanesis* [100] | ST | GCA_008973525.1 | 451,494,475 | 1,296.7 | 552 | C:94.2%[S:90.8%,D:3.4%],F:3.4%,M:2.4%,n:2442 |

We downloaded existing Trichoptera genomes from GenBank (https://www.ncbi.nlm.nih.gov/genome/) or Lepbase (http://download.lepbase.org/v4/) and used these in conjunction with our newly generated genomes to analyze genome size evolution as explained in the following sections of this manuscript.

#### Flow cytometry

In addition to genomic sequence data, we used flow cytometry to detect genome size variation across the order. Our study increased the number of species with available flow cytometry-based genome size estimates from 4 [55] to 31. Estimates were submitted to the Animal Genome Size Database (http://www.genomesize.com).

### Analysis

#### Genome size evolution in Trichoptera

Based on the genomes of six trichopteran species, Olsen et al. [30] found a 3-fold suborder-specific increase of genome size and hypothesized that genome size variation is correlated with their phylogeny. To test this hypothesis, we first reconstructed phylogenetic relationships by analyzing ~2,000 single-copy BUSCO genes from the 26 study species (Figs. 1 & 2, Fig. S48). We obtained a molecular phylogeny that was in agreement with recent phylogenetic hypotheses ([56], see supplementary note 6) and which showed that Trichoptera is divided into two suborders: Annulipalpia (Figs. 1 & 2: Clade A, blue) and Integripalpia [consisting of basal Integripalpia (Fig. 1: Clade B1-3, light green) and infraorder Phryganides (Fig. 1: clade B4, dark green)]. Trichopterans use silk to build diverse underwater structures (see illustrations Fig. 1; supplementary note 6, supplementary Fig. S48). Thus, we refer to Annulipalpia as ‘fixed retreat- and net-spinners’, to Phryganides (Integripalpia) as ‘tube case-builders’, and to basal Integripalpia as ‘cocoon-builders’.

**Fig. 1:**
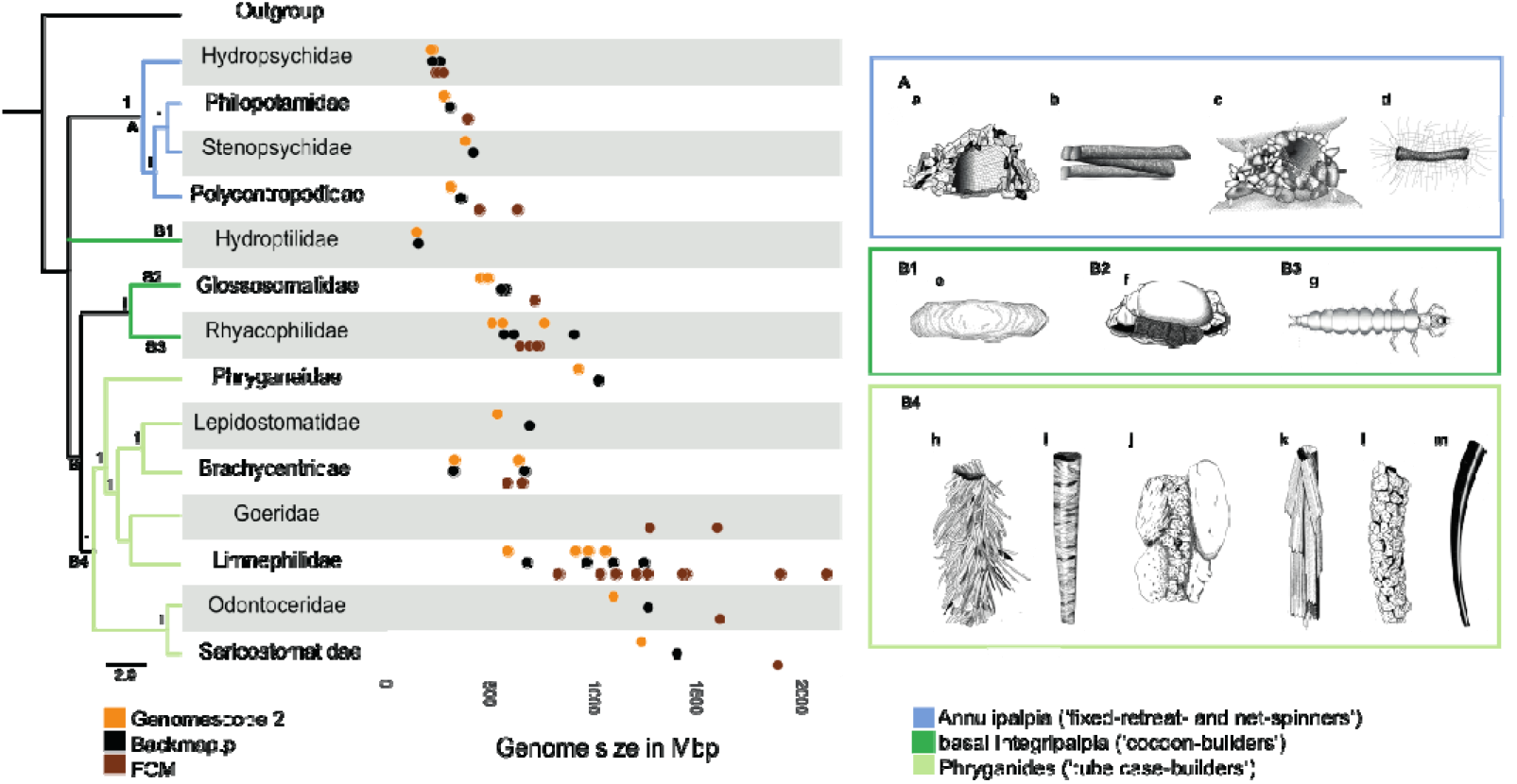
Ecological diversity (right) and genome size (left) in caddisflies. Phylogenetic relationships derived from ASTRAL-III analyses using single BUSCO genes. Goeridae, which was not included in the BUSCO gene set, was placed according to [56]. ASTRAL support values (local posterior probabilities) higher than 0.9 are given for each node. The placement of Hydroptilidae (clade B1) was ambiguous. Since its placement was poorly supported in our analyses, we placed it according to Thomas et al. [56]. Taxa were collapsed to family level. Trichoptera are divided into two suborders: Annulipalpia (‘fixed retreat- and net-spinners’, clade A: blue) and Intergripalpia (clade B: green) which includes basal Integripalpia (‘cocoon-builders’, clades B1-B3, dark green) and Phryganides or ‘tube case-builders’ (clade B4: light green). ‘Cocoon-builders’ are divided into ‘purse case’-(clade B1), ‘tortoise case-building’ (clade B2) and ‘free-living’ (clade B3) families. Genome size estimates based on different methods (Genomescope2: orange, Backmap.pl: black, Flow Cytometry (FCM): brown) are given for various caddisfly families. Each dot corresponds to a mean estimate of a species. For detailed information on the species and number of individuals used in each method see Data S1_Sup.7 -Genome size - Summary. Colors and clade numbers in the phylogenetic tree refer to colored boxes with illustrations. The following species are illustrated by Ralph Holzenthal: a: *Hydropsyche sp.* (Hydropsychidae); b: *Chimarra sp.* (Philopotamidae); c: *Stenopsyche sp.* (Stenopsychidae); d: *Polycentropus* sp. (Polycentropodidae); e: *Agraylea sp.* (Hydroptilidae); f: *Glossosoma sp.* (Glossosomatidae); g: *Rhyacophila sp.* (Rhyacophilidae); h: *Fabria inornata* (Phryganeidae); i: *Micrasema* sp. (Brachycentridae); j:*Goera fuscula* (Goeridae); k: *Sphagnophylax meiops* (Limnephilidae); l: *Psilotreta sp.* (Odontoceridae), m: *Grumicha grumicha* (Sericostomatidae).

**Fig. 2:**
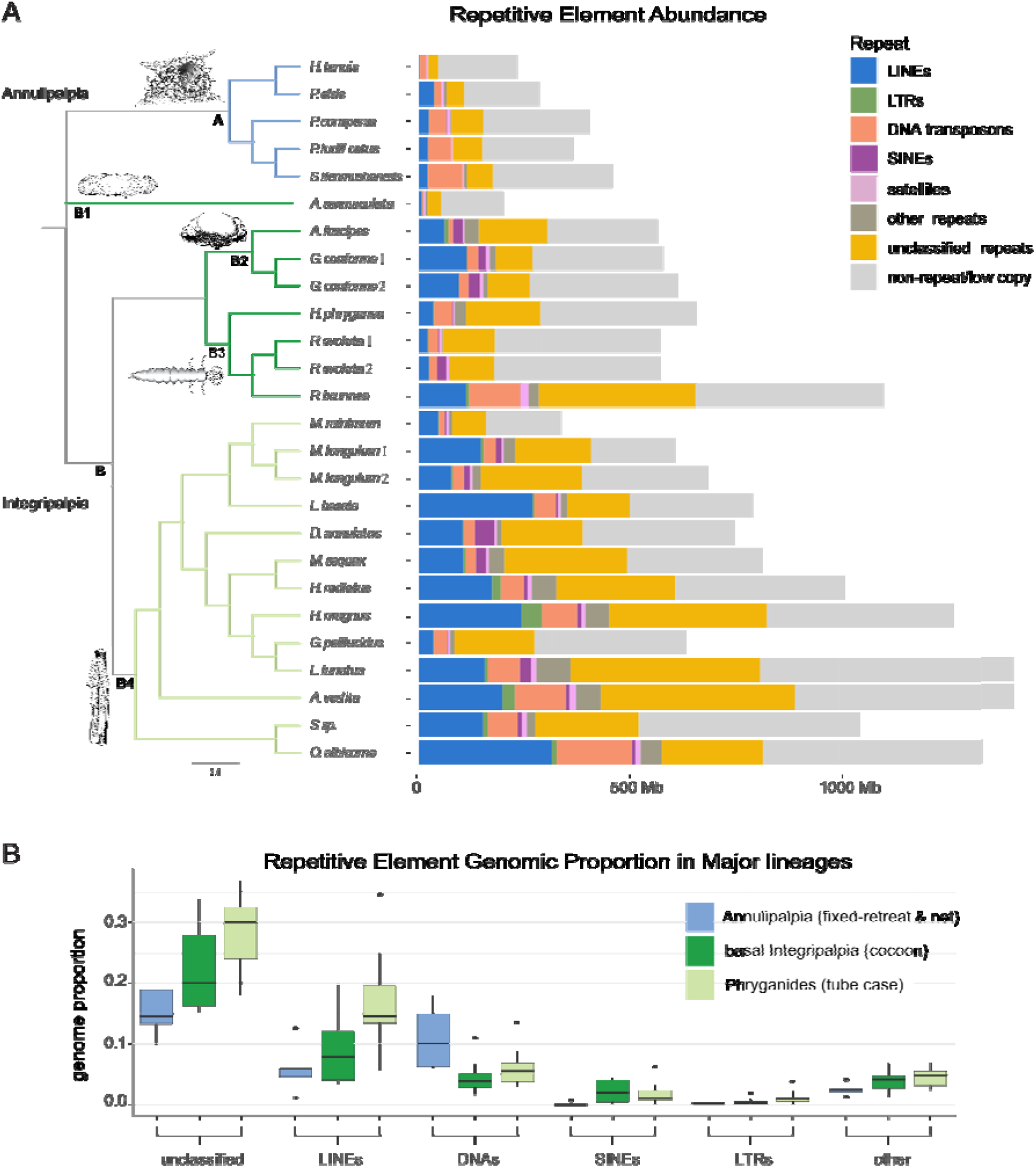
Repeat abundance and classification in 26 caddisfly genomes. Number of bp for each repeat type is given for each caddisfly genome. A: Repeat abundance and classification. Phylogenetic tree was reconstructed with ASTRAL-III using single BUSCO genes from the genome assemblies. The placement of Hydroptilidae (clade B1) was ambiguous. Since its placement was poorly supported in our analyses, we placed the single hydroptilid taxon (*Agraylea sexmaculata*) according to Thomas et al. [56]. Species names corresponding to the abbreviations in the tree can be found in Table 1. Trichoptera are divided into two suborders: Annulipalpia (‘fixed retreat- and net-spinners’, clade A: blue) and Intergripalpia (clade B: green) which includes basal Integripalpia (‘cocoon-builders’, clades B1-B3, dark green) and Phryganides or ‘tube case-builders’ (clade B4: light green). ‘Cocoon-builders’ are divided into ‘purse case’-(clade B1), ‘tortoise case-building’ (clade B2) and ‘free-living’ (clade B3) families. An illustration of a representative of each clade is given. The “other_repeats” category includes: rolling-circles, *Penelope*, low-complexity, simple repeats, and small RNAs. B: Boxplots summarizing shifts in the genomic proportion of RE categories in major Trichoptera lineages.

We used three approaches for estimating genome size across Trichoptera: *k-mer* distribution-estimates, backmapping of sequence data to available draft genomes (as described in [57]), and flow cytometry (FCM, supplementary note 7, supplementary figures S49-S72, DataS1_Sup.7). FCM estimates can be affected by chromatin condensation, the proportion of cells in G0 to G1 phases [58], [59] and endoreplication in insect cells and tissues [60]. Sequence-based estimates can be affected by repetitive elements in the genome resulting in smaller genome size estimates (e.g. [61], [55], [62]), as well as by GC-content because sequence library preparation including PCR amplification steps are associated with underrepresentation of GC and AT rich regions [63]. Bland-Altman plots (supplementary note 8, Fig. S73) revealed general agreement of all three methods in our study. However, the FCM estimates were generally higher compared to the sequence-based estimates (Fig. 1, DataS1_Sup.7) and, among all three approaches, this measure is expected to be the most accurate [8]. We observe that variation among the methods increased with genome size, indicating issues potentially caused by repeat content (see Results *Repeat dynamics*).

We observed large variation in genome size across the order. Genome size tends to be lower in ‘fixed retreat- and net-spinners’ and ‘cocoon-builders’ compared to ‘tube case-builders’ (Fig. 1). Specifically, we observe that genome size varies ~14-fold ranging from 1C = 154 Mbp in ‘cocoon-builders’ (Fig. 1, B1: Hydroptilidae) to 1C = 2129 Mbp in ‘tube case-builders’ (Fig. 1, clade B4: Limnephilidae). Of the 29 species analyzed by FCM, *Halesus digitatus* (Fig. 1, clade B4: Limnephilidae, Intergripalpia) possessed the largest genome (1C = 2129 Mbp), while the genome of *Hydropsyche saxonica* (Fig. 1, clade A: Hydropsychidae, ‘fixed retreat- and net-spinners’) was the smallest (1C = 242 Mbp). Genome size estimates based on sequence-based methods (*kmer*-based and back-mapping) range from 1C = 154 - 160 Mb in *Agraylea sexmaculata* (Fig. 1, clade B1: Hydroptilidae, ‘cocoon-builders’) to 1C = 1238 - 1400 Mbp in *Sericostoma* sp. (Fig. 1, clade B4: Sericostomatidae, ‘tube case-builders’).

### Repeat Dynamics

#### Repetitive element abundance and classification

To understand the structural basis of genome size variation across the order Trichoptera we explored repetitive element (RE) content. We found that major expansions of transposable elements (TEs) contribute to larger genomes in ‘tube case-’ and some ‘cocoon-builders’, but particularly in ‘tube case-builders’ with an average of ~600 Mbp of REs compared to ~138 Mbp in ‘fixed retreat- and net-spinners’ (Fig. 2 A, B). LINEs are the most abundant classified TEs in ‘cocoon-’ and ‘tube case-builders’ and comprise >154 Mb on average in ‘tube case-builders’, or an average genome proportion of 16.9% (range = 5.6–34.7%). This represents a 1.8- and 2.8-fold increase in genome proportion relative to ‘cocoon-builders’ and ‘fixed retreat- and net-spinners’, respectively. The LINE abundance of >312 Mbp in *Odontocerum albicorne* exceeds the entire assembly lengths (152–282 Mbp) of the three smallest genome assemblies (*Hydropsyche tenuis*, *Parapsyche elsis*, and *Agraylea sexmaculata*) (Fig. 2 A, B). DNA transposons also comprise large genomic fractions in both ‘cocoon-’ and ‘tube case-builders’ (averages of 54.4 Mbp and 32.8 Mbp, respectively). However, despite containing a large number of bps, they make up a smaller fraction of total bps in the genomes of ‘cocoon-’ and ‘tube case-builders’ than in ‘fixed retreat- and net-spinners’ (average genome proportion = 5.9%, 4.5%, and 11.1% in ‘tube case-builders’, ‘cocoon-builders’, and ‘fixed retreat- and net-spinners’, respectively) (Fig. 2 B), and thus cannot, by themselves, explain the larger genome sizes. SINEs, LTRs, *Penelope* (grouped with “other” repeats in Fig. 2), and satDNAs show a disproportionate increase in ‘cocoon-’ and ‘tube case-builders’, however, all categories combined make up a relatively small proportion of their genomes (all less than 3% on average in Integripalpia) (Fig. 2, B). Unclassified repeats are the most abundant repeat category across all Trichoptera, and they also show disproportionate expansions in both ‘cocoon-’ and ‘case-builders’ relative to ‘fixed retreat- and net-spinners’ (Fig. 2 A, B). The general trends noted in our assembly-based analysis of REs were corroborated by our reference-free analysis of repeat abundance (Figs. S122, S123 supplementary note 10).

#### TE age distribution analysis

To test whether the observed abundance patterns of specific TEs are driven by shared ancient proliferation events or more recent/ongoing activity of the respective TEs, we analyzed TE age distribution plots. These plots allow us to visualize specific RE classes/superfamilies that account for shifts in RE composition and abundance and infer the relative timing of those shifts based on the distribution of sequence divergence within each RE category. TE age distributions showed a high abundance of recently diverged TE sequences in ‘cocoon-’ and ‘tube case-builders’, particularly in LINEs, DNA transposons, and LTRs in which the majority of TEs for a given class show 0–10% sequence divergence within copies of a given repeat (Fig. 3). This trend was particularly pronounced among ‘tube case-builders’ with several species showing high abundance of LINEs and DNA transposons with 0–5% sequence divergence (Fig. 3). This pattern suggests that the observed TE expansion is due primarily to ongoing TE activity within lineages rather than a few shared bursts of activity in ancestral lineages. This is further supported by our analysis of repeat sub-classes with age distribution plots (Fig. S124). For example, in our study, LINE abundance is often due to the expansion of different LINE subclasses even between species in the same sub-clade (e.g., compare *Lepidostoma* with *Micrasema*, *Himalopsyche* with *Glossosoma*; Fig. S124). We also find evidence of shared ancient bursts of SINE activity in ‘cocoon-’ and ‘tube case-builders’, although SINEs are not an abundant repeat class in any species (avg. genomic proportion=1.9% stdev=1.7%) (Fig. S124).

**Fig. 3:**
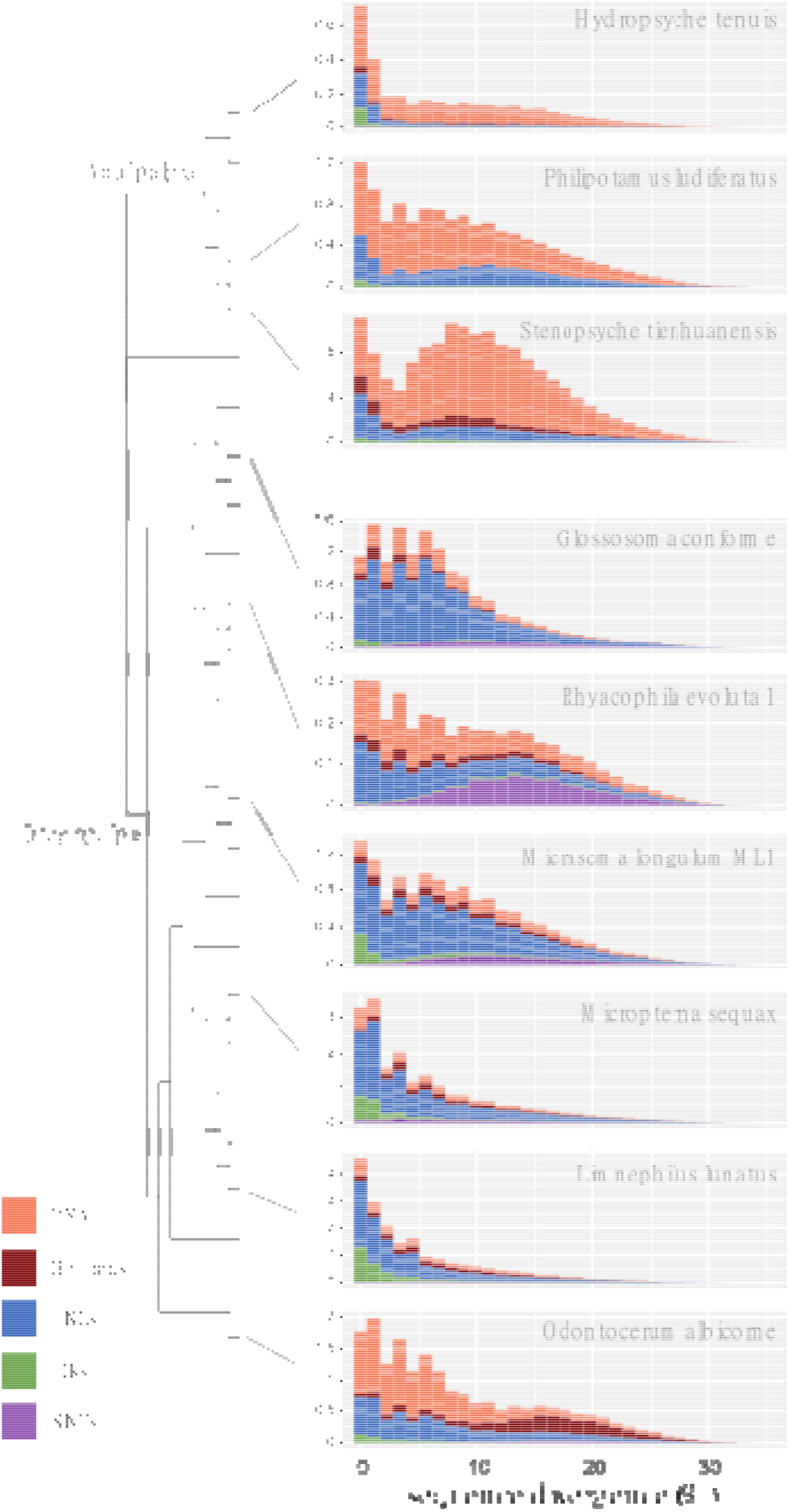
Transposable element age distribution landscapes. Representative examples are chosen from major Trichoptera lineages. The y-axis shows TE abundance as a proportion of the genome (e.g., 1.0 = 1% of the genome). The x-axis shows sequence divergence relative to TE consensus sequences for major TE classes. TE classes with abundance skewed toward the left (i.e., low sequence divergence) are inferred to have a recent history of diversification relative to TE classes with right-skewed abundance. Plots were generated in dnaPipeTE. Plots for all species are shown in supplementary Fig. S123. For tip labels of the phylogenetic tree see Fig. 2.

#### Associations between TE sequences and protein-coding genes

During early exploration of our sequence data, we made an unexpected discovery that in some lineages, universal single copy orthologs or “BUSCO genes”, showed higher than expected coverage depth of mapped reads in one or more of their sequence fragments. Further analysis showed that these high coverage BUSCO sequence regions are typically RE sequences (primarily TEs) that are either embedded within or located immediately adjacent to BUSCO genes, such that the BUSCO algorithm includes them in its annotation of a given gene. We refer to BUSCO genes containing these putative RE fragments as ‘TE-associated BUSCOs’ (supplementary Fig. S125, supplementary note 11). By estimating how many times they occur, we can quantitatively measure how TE-gene interactions change with changing genome size. In fact, we detected a positive linear relationship between TE-gene interactions and increasing genome size when measured with this accidently discovered metric. We found major expansions of TE-associated BUSCOs in ‘cocoon-’ and ‘tube case-builders’ (Fig. 4A) that are significantly correlated with total repeat abundance, as well as the genomic proportion of LINEs and DNA transposons (supplementary Fig. S126). TE-associated BUSCOs comprise a relatively large fraction of total BUSCO genes in these lineages (averages of 11.2% and 21.4% of total BUSCOs in ‘cocoon-‘ and ‘tube case-builders’, respectively), compared to annulipalpian lineages (avg = 6.2%). This finding highlights the major impact of REs on the composition of protein-coding genes in species with repeat-rich genomes. The BUSCO-associated sequences may represent TEs recently inserted into BUSCO genes, the remnants left behind following historical TE transposition events, or TE sequences that are immediately adjacent to and inadvertently classified as BUSCO sequences.

**Fig. 4:**
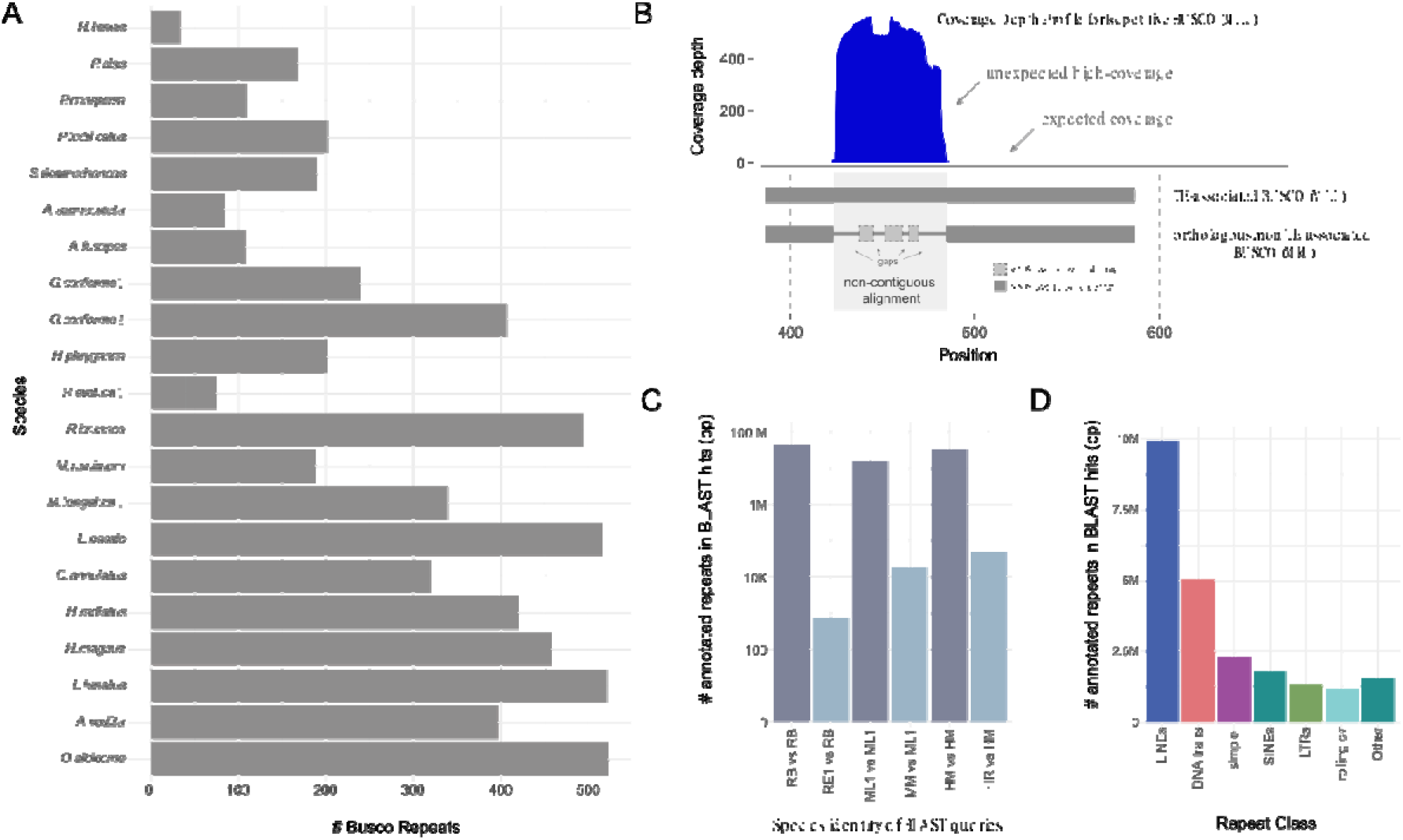
TE-BUSCO-gene associations in Trichoptera species. (A) Raw abundance of TE-associated BUSCO sequences present in the assembly of 2442BUSCOs in the OrthoDB 9 Endopterygota dataset. (B) Upper plot: An example of a coverage depth profile of a TE-associated BUSCO gene [BUSCO EOG090R02Q9 from ML1 (‘inflated species’)] which shows unexpected high coverage in the second exon putatively due to the presence of a RE-derived sequence fragment. Lower plot: A typifying alignment between a TE-associated BUSCO and its orthologous BUSCO from a closely related species (‘reference species’) that lack TE-association. The non-TE-associated orthologous BUSCO shows non-contiguous alignment in regions of inflated coverage in the TE-associated BUSCO, consistent with the presence of a RE-derived sequence fragment in the TE-associated BUSCO that is absent in the reference species. (C) Summary of total bases annotated as REs obtained from each of two BLAST searches. First, when we used BLAST to compare an TE-associated BUSCOs against an assembly for the same species BLAST hits included megabases of annotated repeats (dark plots). Second, when non-TE-associated orthologs of the TE-associated BUSCOs in the first search are taken from a close relative and compared against the inflated species using BLAST, there is a dramatic drop in BLAST hits annotated as REs. Note log scale on the y-axis. (D) Summary of annotations for BLAST hits for classified REs when TE-associated BUSCOs are compared against an assembly of the same species using BLAST.

To confirm that unexpectedly high-coverage sequence regions in TE-associated BUSCOs were in fact TE-derived sequences, we compared patterns of BUSCO gene structure (though pairwise alignment) across species pairs in which high-coverage regions (i.e., putative TE sequences) were present in the BUSCO gene of one species (i.e., the “inflated” species), but absent in the homologous BUSCO of the other (i.e., the “reference” species). This analysis showed that in 73 of 75 randomly sampled alignments, reference species showed gaps or highly non-contiguous alignments in high-coverage regions of the inflated species (Fig. 4B), suggesting that sequence insertions are typically present in high-coverage sequence regions of TE-associated BUSCOs. Our subsequent BLAST analysis showed that comparing a TE-associated BUSCO against its own assembly produced thousands-millions of BLAST hits from many contigs (Fig. 4C). This confirmed that the indel sequence present in high-coverage regions of “inflated” species show high sequence similarity to repetitive elements elsewhere in the genome. We then used an intersect analysis on the BLAST results to confirm that the large majority of the excessive BLAST hits overlap with RE annotations throughout the genome, most of which are TEs with LINEs and DNA transposons being most abundant (Fig. 4D, DataS2_Sup.5). Finally, we found that if we replaced the TE-associated BLAST query sequence with the homologous, but non-TE associated BUSCO from its counterpart reference species, the number of BLAST hits was fewer (Fig. 4C, DataS2_Sup.6), offering further evidence that the TE sequence insertions driving the pattern of high-coverage in read mapping excessive BLAST hits are absent in reference species and thus carriable across relatively short time scales within Trichoptera. Taken together, these findings provide strong evidence that TE sequences (especially LINEs and DNA transposons) inadvertently annotated by BUSCO can account for the high-coverage regions we observe in BUSCO genes (Fig. 4D).

Our accidental discovery that quantifying the frequency of TE-associated BUSCOs can serve as an estimate of TE-gene associations may prove useful in other systems given the wide use of BUSCO analysis in genomic studies. Finer details supporting the TE-gene association analysis are reported in supplementary note 11.

#### Gene and genome duplications

Recently, a transcriptome-based study found evidence for putative ancient gene and genome duplications in hexapods, including potential WGD events in caddisflies [22], suggesting that duplication events could be responsible for some genome size variation in Trichoptera. We investigated whether this pattern persists with whole genome data and found that the age distribution of duplications in 18 genomes were significantly different compared to the background rate of gene duplication (Figs. S137 & S138). To identify if any significant peak is consistent with a potential WGD, we used mixture modeling to identify peaks in these gene age distributions, which recovered no obvious peak consistent with an ancient WGD. To further investigate potential WGD, we used Smudgeplot [64] to visualize the haplotype structure and to estimate ploidy of the genomes.

While Smudgeplot predicted most of the genomes to be diploid, four genomes with rather small genome sizes (230 Mb – 650 Mbp) were predicted to be tetraploid (*Hydropsyche tenuis*, *Rhyacophila evoluta* RSS1 and HR1, *Parapsyche elsis*). However, the Genomescope 2 results indicate that these are highly homozygous samples. Low heterozygosity is a known confounder of smudgeplot analyses (see https://github.com/KamilSJaron/smudgeplot/wiki/tutorial-strawberry) because it inflates the signal of duplication when compared to the low level of heterozygosity. We therefore interpret these four putative polyploids as artifacts of low heterozygosity in the analysis.

## Discussion

The drivers and evolutionary consequences of genome size evolution are a topic of ongoing debate. Several models have been proposed [8]. Some hypothesize genome size to be a (mal)adaptive trait by impacting phenotypic traits such as developmental/life history, body size and other cell-size related effects [65], [66], [67], [68] reviewed in [8]. On the other hand, neutral theories suggest that DNA accumulation occurs only by genetic drift without selective pressures playing a major role in the accumulation or loss of DNA [the mutational hazard hypothesis (MHH, [23]) and the mutational equilibrium hypothesis (MEH, [24])]. The MHH only allows for small deleterious effects for the accumulation of extra DNA which is accompanied by higher mutation rates in larger genomes [23], while the MEH focuses on the balance between insertions and deletions. It suggests that genome expansions arise by ‘bursts’ of duplication events or TE activity and genome shrinkage may be caused by a more constant rate of small deletions [24].

In this study, we observe that genome size varies ~14-fold across the order Trichoptera, with lower genome size estimates in ‘fixed retreat- and net-spinners’ and ‘cocoon-builders’ compared to ‘tube case-builders’ and explore potential drivers of genome size evolution. Although, recent genomic studies have shown evidence of bursts of gene duplication and gene family expansion during the evolution of hexapods [22], [69] the presence of ancient genome duplication events are still a subject of debate [70], [71], [72]. We found neither evidence for whole duplication events when computing haplotype structure and ploidy with Smudgeplot, nor evidence of ancient WGD in the gene age distribution in our Trichoptera genomes although we recognize that some of our current genome assemblies might be too fragmented to infer synteny. This does not mean that we can rule out that duplication events played a role in genome size evolution in Trichoptera in the past. The emergence of PacBio HiFi genomes of caddisflies (e.g., Darwin Tree of Life is currently planning to sequence 28 caddisfly genomes; https://www.darwintreeoflife.org/) will allow a deeper exploration of putative ancient duplication events in Trichoptera.

We found evidence that TE expansions (especially LINEs) were important drivers of genome size evolution in Trichoptera (Fig. 2, Figs. S122 & S123), which is consistent with the mutational equilibrium hypothesis (MEH). The TE age distribution analyses suggested that the high abundance of LINEs was due to ongoing/recent activity occurring independently across ‘cocoon-’ and particularly ‘tube case-builders’ (Fig. 3, Fig. S124). Thus, the shift to large genomes in these lineages does not appear to be due to a single (or few) shared ancient events, rather they maintained dynamic turnover in composition of their large genomes. Mutational bias affecting pathways tied to TE-regulation may affect insertion/deletion ratios and subsequently lead to lineage-specific shifts in genome size equilibrium [73]. Such changes may be stochastic (e.g., due to drift), or linked to traits that evolve on independent trajectories as lineages diverge and are thereby constrained by phylogeny. Ecological factors, demographic history, and effective population size can further impact mutation rates. For example, environmental stress can trigger bursts of TE activity and elevated mutation rates [74], [75], [76] driving lineages that occupy niche space with frequent exposure to environmental stress toward increased TE loads and larger genomes. Similarly, lineages with small effective population sizes or which are prone to population bottlenecks may have higher mutation rates and/or reduced efficacy of natural selection which would otherwise purge mildly deleterious TE load.

Although our study is not designed to pinpoint specific forces maintaining large genomes in some lineages, the pattern we observe in the distribution of genome size (i.e. lower genome size estimates in ‘fixed retreat- and net-spinners’ and ‘cocoon-builders’ compared to ‘tube case-builders’) leads us to hypothesize that ecological factors may play a role in genome size evolution in the order. The three focal groups discussed here exhibit markedly different ecological strategies. Larvae of ‘fixed retreat- and net-spinners’ generally occupy relatively narrow niche space in oxygen-rich flowing-water (mostly stream/river) environments where they rely on water currents to bring food materials to their filter nets. The evolutionary innovation of tube-case making is thought to have enabled ‘tube case-builders’ to occupy a much greater diversity of ecological niche space by allowing them to obtain oxygen in lentic (e.g., pond, lake, marsh) environments which are much more variable in temperature and oxygen availability than lotic environments [77], [78]. This environmental instability is greater over short (daily, seasonal) and long-time scales (centuries, millennia) [79]. It is thus plausible these tube case-building lineages experience greater environmental stress and less stable population demographics that could lead to both more frequent TE bursts and reduced efficacy of natural selection in purging deleterious effects of TE expansions as described above [23], [24].

We show that TE expansions (especially LINEs and DNA transposons) in ‘cocoon-’ and ‘tube case-builders’ have a major impact on protein-coding gene regions (Fig. 4). These TE-gene associations show a linear relationship with increasing genome size. This trend is particularly pronounced among ‘tube case-builders’ in which TE-associated BUSCOs comprise an average of 21.4% of total BUSCO genes (compared to 6.2% in annulipalpians). This finding corroborates other studies highlighting the role of TEs as drivers of rapid genome evolution [80], [81], [82], [83] and highlights their impact on genomic regions that have potential effects on phenotypes. Questions remain as to what evolutionary roles such changes in genic regions may play. In general, TE insertions are considered to have deleterious effects on their host’s fitness activity [84], [85]. They are known to “interrupt” genes [33], pose a risk of ectopic recombination that can lead to genome rearrangements [34], [31], [86], and have epigenetic effects on neighboring sequences [87], [88]. Therefore, purifying selection keeps TEs at low frequencies [33]. However, there is growing evidence that TE activity can also be a critical source of new genetic variation driving diversification via chromosomal rearrangements and transposition events which can result in mutations [89], including examples, of co-option [90], e.g. recent research in mammals has shown that DNA transposon fragments can be co-opted to form regulatory networks with genome-wide effects on gene expression [44].

Ecological correlates with genome size are widely discussed in other taxa [91], [92], [93], [94], [95]. Caddisflies and other diverse insect lineages that feature various microhabitat specializations, feeding modes, and/or the use of silk represent evolutionary replicates with contrasting traits and dynamic genome size evolution. They thus have high potential as models for understanding links between ecology and the evolution of REs, genomes, and phenotypes. Our study lays a foundation for future work in caddisflies that investigates the potential impact of TE expansions on phenotypes and tests for evidence of co-option/adaptive impacts of TE-rich genomes against a null of neutral or slightly deleterious effects.

### Potential implications

Many open questions remain as to the causes and consequences of genome size evolution. As we move forward in an era where genome assemblies are attainable for historically intractable organisms (e.g. due to constraints given large genome sizes, tissue limitations, no close reference available) we can leverage new model systems spanning a greater diversity of life to understand how genomes evolve. Here, we provide genomic resources and new genome size estimates across lineages of an underrepresented insect order that spans major variation in genome size. These data allowed us to study genome size evolution in a phylogenetic framework to reveal lineage-specific patterns in which genome size correlates strongly with phylogeny and ecological characteristics within lineages. We find that large genomes dominate lineages with a wider range of ecological variation, and that ongoing recent TE activity appears to maintain large genomes in these lineages. This leads us to hypothesize that ecological factors may be linked to genome size evolution in this group. The future directions spawned by our findings highlight the potential for using Trichoptera and other diverse insect groups to understand the link between ecological and genomic diversity, a link that has been challenging to study with past models [8].

We also show that TE expansions are associated with increasing genome size and have an impact on protein-coding regions. These impacts have been greatest in the most species-rich and ecologically diverse caddisfly clades. While TEs are generally considered to have deleterious effects on their host’s fitness activity, their roles can also be neutral or even adaptive. TE activity can be a critical source of new genetic variation and thus an important driver for diversification. Caddisflies and potentially other non-model insect groups are excellent models to test these contrasting hypotheses, as well as the potential impact of TEs on phenotypes. Using these models, especially with respect to the increasing emergence of high-quality insect genomes [96], will allow researchers to identify recurring patterns in TE dynamics and investigate their evolutionary implications across diverse clades.

## Methods

### DNA extraction, library preparation, sequencing, and sequence read processing

We extracted high molecular weight genomic DNA (gDNA) from 17 individuals (15 species) of caddisfly larvae (for sampling information, see DataS1_Sup.1) after removing the intestinal tracts using a salting-out protocol adapted from [97] as described in supplementary note 1. We generated gDNA libraries for a low-cost high-contiguity sequencing strategy, i.e. employing a combination of short (Illumina) and long read (Nanopore or PacBio) technologies as described in supplementary notes 2. For details on sequencing coverage for each specimen see DataS1_Sup.3.

### *De novo* genome assembly, annotation and quality assessment

We applied different assembly strategies for different datasets. First, we applied a long-read assembly method using wtdbg2 v2.4 [98] with subsequent short-read polishing with Pilon v1.22 [99] as this method revealed good results in previous *de novo* assemblies in caddisflies [55]. In cases where this pipeline did not meet the expected quality regarding contiguity and BUSCO completeness, we applied *de novo* hybrid assembly approaches of MaSuRCA v.3.1.1 [100] (supplementary note 3). Illumina-only data was assembled with SPAdes [101] explained in supplementary note 3. Prior to annotating the individual genomes with MAKER2 v2.31.10 [102], [103] we used RepeatModeler v2.0 and RepeatMasker v4.1.0, to identify species-specific repetitive elements in each of the assemblies, relative to RepBase libraries v20181026; www.girinst.org). Transcriptome evidence for the annotation of the individual genomes included their species-specific or closely related *de novo* transcriptome provided by 1KITE (http://www.1kite.org/; last accessed November 11, 2019, DataS1_Sup.9) or downloaded from Genbank as well as the cDNA and protein models from *Stenopsyche tienmushanensis* [104] and *Bombyx mori* (AR102, GenBank accession ID# GCF_000151625.1). Additional protein evidence included the uniprot-sprot database (downloaded September 25, 2018). We masked repeats based on species-specific files produced by RepeatModeler. For *ab initio* gene prediction, species specific AUGUSTUS gene prediction models as well as *Bombyx mori* SNAP gene models were provided to MAKER. The EvidenceModeler [105] and tRNAscan [106] options in MAKER were employed to produce a weighted consensus gene structure and to identify tRNAs genes. MAKER default options were utilized for BLASTN, BLASTX, TBLASTX searches. Two assemblies (*Agapetus fuscipens* GL3 and *Micrasema longulum* ML1) were not annotated because of their low contiguity. All protein sequences were assigned putative names by BlastP Protein–Protein BLAST 2.2.30+ searches [107] and were functionally annotated using command line Blast2Go v1.3.3 [108], see supplementary note 4, Figs. S1-S30).

We calculated assembly statistics with QUAST v5.0.2 [51] and examined completeness with BUSCO v3.0.2 [52], [53] using the Endopterygota odb9 dataset with the options --*long,* –*m = genome* and *–sp= fly*. A summary of the assembly statistics and BUSCO completeness is given in Table 1. The final genome assemblies and annotations were screened and filtered for potential contaminations with taxon-annotated GC-coverage (TAGC) plots using BlobTools v1.0 [54]. Details and blobplots are given in supplementary note 5 & supplementary Figs. S31-S47.

### Species tree reconstruction

We used the single-copy orthologs resulting from the BUSCO analyses to generate a species tree. We first combined single-copy ortholog amino acid files from each species into a single FASTA for each ortholog. We then aligned them with the MAFFT L-INS-i algorithm [109]. We selected amino acid substitution models for each ortholog using ModelFinder (option -m mfp,[110] in IQtree v.2.0.6 [111] and estimated a maximum likelihood tree with 1000 ultrafast bootstrap replicates [112] with the BNNI correction (option -bb 1000 -bnni). We combined the best maximum likelihood tree from each gene for species tree analysis in ASTRAL-III [113]. A locus tree was inferred using the alignment file (-s) and the partition file (-S) with the settings –prefix loci and -T AUTO in IQtree. Gene and site concordance factors were calculated with IQTree using the species tree (-t), the locus tree (--gcf) and the alignment file (-s) with 100 quartets for computing the site concordance factors (--scf 100) and --prefix concord for computing the gene concordance factors. We visualized the trees using FigTree v.1.4.4 (http://tree.bio.ed.ac.uk/software/figtree/).

### Genome size estimations and genome profiling

Genome size estimates of 27 species were conducted using flow cytometry (FCM) according to Otto [114] using *Lycopersicon esculentum* cv. Stupické polnítyčkové rané (2C◻=L◻.96Lpg;[115]) as internal standard and propidium iodine as stain. Additionally, we used trimmed, contamination filtered short-read data (see supplementary note 2) to conduct genome profiling (estimation of major genome characteristics such as size, heterozygosity, and repetitiveness) using a *k-mer* distribution-based method (GenomeScope 2.0, [64]. Genome scope profiles are available online (see links to Genomescope 2 in DataS1_Sup.4). In addition, we applied a second sequencing-based method for genome size estimates, which uses the backmapping rate of sequenced reads to the assembly and coverage distribution (backmap.pl v0.1, [57]. Details of all three methods are described in supplementary note 7. Coverage distribution per position and genome size estimate from backmap.pl are shown in Figs. S49-72). We assessed the congruence among the three quantitative methods of measurement (Genomescope2, Backmap.pl and FCM) with Bland-Altman-Plots using the function BlandAltmanLeh::bland.altman.plot in ggplot2 [116] in RStudio (RStudio Team (2020). RStudio: Integrated Development for R. RStudio, PBC, Boston, MA URL http://www.rstudio.com/; supplementary note 8, Fig. S73).

### Repeat dynamics

#### Repeat abundance and classification

We identified and classified repetitive elements in the genome assemblies of each species using RepeatModeler2.0 [117]. We annotated repeats in the contamination filtered assemblies with RepeatMasker 4.1.0 (http://www.repeatmasker.org) using the custom repeat libraries generated from RepeatModeler2 for each respective assembly with the search engine set to “ncbi” and using the -xsmall option. We converted the softmasked assembly resulting from the first RepeatMasker round into a hardmasked assembly using the lc2n.py script (https://github.com/PdomGenomeProject/repeat-masking). Finally, we re-ran RepeatMasker on the hard-masked genome with RepeatMasker’s internal arthropod repeat library using - species “Arthropoda”. We then merged RepeatMasker output tables from both runs by parsing them with a script (RM_table_parser_families_.py, available at https://github.com/jhcaddisfly/TE-gene_intersect_analysis) and then combined the resulting data columns for the two runs in Excel.

We also estimated repetitive element abundance and composition using RepeatExplorer2[118], [119] and dnaPipeTE v.1.3.1 [120]. These reference-free approaches quantifies repeats directly from unassembled short-read data. These analyses allowed us to test for general consistency of patterns with our assembly-based approach described above, and to test for the presence of abundant repeat categories such as satellite DNAs which can comprise large fractions of genomes yet can be prone to poor representation in the genome assembly. Prior to analysis, we normalized contamination filtered (see supplementary note 2) input data sets to 0.5x coverage using RepeatProfiler [121] and seqtk (https://github.com/lh3/seqtk), and then ran RepeatExplorer2 clustering with the Metazoa 3.0 database specified for annotation (supplementary Fig. S122) and dnaPipeTE with the -RM_lib flag set to the Repbase v20170127 repeat library (supplementary Fig. S123).

#### TE age distribution analysis

We further characterized repetitive element dynamics in Trichoptera by analyzing TE landscapes, which show relative age differences among TE sequences and their genomic abundance. We used these analyses to test whether abundance patterns of specific TEs are driven by shared ancient proliferation events or more recent/ongoing activity of the respective TEs. For example, if shared ancient proliferation is driving abundance patterns of a given TE, the majority of its copies would show moderate to high sequence divergence (e.g., >10% pairwise divergence). In contrast, if abundance patterns are driven by recent/ongoing activity of a given TE, we would expect the majority of its sequences to show low sequence divergence (e.g., 0–10%). We generated TE age distribution plots using dnaPipeTE v1.3.1 [120] with genomic coverage for each species sampled to 0.5X prior to analysis and the - RM_lib flag set to the Repbase v20170127 repeat library (supplementary Fig. S124).

#### TE sequence associations with protein-coding genes

We analyzed BUSCO genes for all species to quantify the abundance of TE-associated BUSCOs across samples and investigated associations between TEs and genic sequences in Trichoptera lineages by quantifying the abundance of TE-associated BUSCO genes (for presence and absence of TE-associated BUSCOs see Fig. S125, DataS2_Sup.3). This analysis also allowed us to quantify shifts in associations between TEs and genic regions across Trichoptera lineages with varying repeat abundance. We identified BUSCO genes with high-coverage sequence regions based on coverage profiles and quantified their genomic abundance by using each TE-associated BUSCO as a query in a BLAST search against their respective genome assembly. We then conducted intersect analysis for all unique BUSCO hits from high coverage sequences to determine if these were annotated as TEs. We calculated the total number of bases in filtered BLAST after subtracting the number of bases at the locus belonging to all ‘complete’ BUSCO genes and categorized high coverage sequence regions in BUSCO genes based on their annotation status and repeat classification using custom scripts (available at https:// https://github.com/jhcaddisfly/TE-gene_intersect_analysis). We plotted the number of the high coverage BUSCO sequence regions belonging to repetitive element categories (i.e., classes and subclasses) alongside plots of the relative genomic abundance of each respective category. In addition, we investigated BUSCO genes with regions of high coverage by pairwise alignments. Specifically, we visualized alignments of BUSCOs with high coverage sequence regions (i.e., the “inflated species”) alongside orthologous BUSCOS that lack such regions taken from closely related species (i.e., the “reference” species). We further tested this prediction by taking the set of BUSCOs that only exhibited high coverage regions in the inflated species and contrasted results of two BLAST searches followed by an intersect analysis. A detailed description of this method is provided in supplementary note 11.

### Gene and Genome duplications

#### Inference of WGDs from gene age distributions

To recover signal from potential WGDs, for each genome, we used the DupPipe pipeline to construct gene families and estimate the age distribution of gene duplications [122], https://bitbucket.org/barkerlab/evopipes/src/master/). We translated DNA sequences and identified ORFs by comparing the Genewise [123] alignment to the best□hit protein from a collection of proteins from 24 metazoan genomes from Metazome v3.0. For all DupPipe runs, we used protein◻guided DNA alignments to align our nucleic acid sequences while maintaining the ORFs. We estimated synonymous divergence (Ks) using PAML with the F3X4 model [124] for each node in the gene family phylogenies. We first identified taxa with potential WGDs by comparing their paralog ages to a simulated null distribution without ancient WGDs using a K-S goodness-of-fit test [125]. We then used mixture modeling to identify if any significant peaks consistent with a potential WGD and to estimate their median paralog Ks values. Significant peaks were identified using a likelihood ratio test in the boot.comp function of the package mixtools in R [126].

#### Visualization of genome structure to estimate ploidy using smudgeplots

We visualized the genome structure and estimated ploidy levels with smudgeplot. For this purpose, we extracted genomic kmers from kmer counts produced with jellyfish (as described above in “Genome size estimation and genome profiling”) using “jellyfish dump” with coverage thresholds previously estimated from kmer histograms using the smudgeplot.py script. We computed the set of kmer pairs with the Smudgeplot tool hetkmers. After generating the list of kmer pair coverages, we generated smudgeplots using the coverage of the kmer pairs and the “plot” tool within Smudgeplot. Ploidy as well as the haploid kmer coverage was estimated directly from the data and compared to the estimates reported by Genomescope2 (see DataS1-Sup.4). Details of the method and smudgeplots are given in supplementary figures S74-121.

## Supporting information

DataS1_Sup.6

DataS1_Sup.1

DataS1_Sup.2

DataS1_Sup.3

DataS1_Sup.4

DataS1_Sup.5

DataS2_Sup.6

DataS1_Sup.7

DataS1_Sup.8

DataS1_Sup.9

DataS1_Sup.10

DataS2_Sup.1

DataS2_Sup.2

DataS2_Sup.3

DataS2_Sup.4

DataS2_Sup.5

## Data and materials availability

This project has been deposited at NCBI under BioProject ID: PRJNA558902. The data sets supporting the results of this article are available in the supplementary, data files S1 and S2 and at https://byu.box.com/v/trich-genomes. The data available at the link will be uploaded to GigaDB when the paper is accepted.

## Declarations

### Consent for publication

Not applicable

## Author’s contributions

Conceptualization –JH, JSS, PBF, SUP

Data curation – JH

Formal Analysis - JH, JM, JP, JSS, PBF, ZL

Funding acquisition – AML, PBF, SUP, RJS, RJS

Investigation – JH, JP, JSS, PBF, ZL

Methodology – AML, JSS, JP, JVS, PBF

Project administration – SUP

Resources – JP, MB, PBF, SUP

Visualization - JH, JSS

Writing – original draft – JH, JSS, PBF, ZL

Writing – review & editing – AML, JH, JM, JP, JSS, JVS, MB, PBF, RJS, RJS, SUP, ZL

## Acknowledgments

The authors thank Ralph Holzenthal for providing illustrations of larval Trichoptera and the structures they build. We thank Bob Wisseman for collecting *Himalopsyche phryganeae.* We thank both reviewers for their insightful critiques and interest in our manuscript.

