## Supplementary Material for "Genome size evolution in the diverse insect order Trichoptera"

**Supplementary note 1: DNA extraction**

For extraction of high molecular weight genomic DNA (gDNA) from 17 individuals (15 species) of caddisfly larvae, we used a salting-out protocol adapted from [1]. After removing the intestinal tracts, tissue lysis was done in 600 µl lysis buffer (10 mM Tris-Hcl, 400 mM NaCl, and 100 mM  EDTA, pH 8.0), 40 µl 10% SDS and 5 µl Proteinase K (20 mg/ml) on a MKR23 thermoshaker (Hettich Benelux, Geldermalsen, Netherlands) overnight at 37° C. After adding 20 µl RNAse A (10 mg/ml), we incubated the samples for another 30 min at 37°C. Then, we added 240 µl 5M NaCl and inverted the tube 5-10 times. To pellet the precipitate, we centrifuged the tube at 10000 rpm at 4°C for 10 min in a Z 300 K centrifuge (Hermle, Gosheim, Germany) and transferred the supernatant to a new tube. After adding 1 volume of chloroform-isoamylalcohol (24:1), we carefully inverted the tube 20-30 times and centrifuged again as above. To pellet the precipitated DNA, we mixed the supernatant with 1.2 ml 100% ethanol in a new tube and centrifuged again for 10 min. After washing the DNA two times with freshly prepared 70% ethanol and subsequent air-drying for 20 min, we resuspended the DNA pellet in 1X TE buffer. We quantified the DNA using a Qubit 4.0 fluorometer with the dsDNA Broad Range Kit (ThermoFisher Scientific, Waltham, USA), checked its purity with a DS11 spectrophotometer (DeNovix, Wilmington, DE, USA) and verified DNA integrity on a 2200 TapeStation (Agilent Technologies, Santa Clara, USA) with a Genomic Tape.

**Supplementary note 2: Sequencing strategies**

***Short read (Illumina) sequencing***

All 17 individuals (15 species) were subject to Illumina paired-end sequencing. We prepared genomic libraries for Illumina sequencing from 400-500 ng gDNA using the NEBNext Ultra II FS DNA Library Preparation Kit (New England Biolabs, Ipswich, MA, USA) following the manufacturer’s manual. To achieve a mean insert size of 400-500 bp, we conducted enzymatic fragmentation with subsequent size selection. We chose a combinatorial dual indexing approach using the NEBNext Multiplex Oligos for Illumina (Dual Index Set 1) and amplified libraries with five PCR cycles. We quantified each library on a Qubit 4.0 fluorometer with the 1x dsDNA HS Assay Kit and checked the fragment size distribution on a 2200 TapeStation using a High Sensitivity D1000 Tape. Illumina paired-end (150 bp) sequencing was done on a HiSeq 2000 sequencer at Novogene. After checking the quality of Illumina reads using FastQC v0.11.8 (<http://www.bioinformatics.babraham.ac.uk/projects/fastqc>), we trimmed of overrepresented k-mers using autotrim.pl v0.6.1 [2] with Trimmomatic v0.38 [3] and a custom adapter file (ILLUMINACLIP: <adapter_combined.fa>:2:30:10), SLIDINGWINDOW:4:20 and MINLEN:50 and further processed reads with Cutadapt v2.23 [4] using the following parameters:-- pair-filter=any -l=140, --max-n=0 for sample which showed a warning flag at per base sequence content in FastQC. To filter out potentially contaminated reads, we used Kraken 2 v2.0.8-beta [5], [6] with the standard Kraken 2 database and only kept unclassified reads for further analyses.

***Long-read sequencing***

In addition to Illumina sequencing, we sequenced 14 individuals (14 species) with Oxford Nanopore or PacBio long-read technologies.

*Oxford Nanopore sequencing*

To generate Oxford Nanopore long-reads we sheared high-molecular-weight gDNA (2.0-3.2 µg; > 60 kb) to a mean fragment size of about 10 kb using g-TUBES (Covaris, Woburn, MA, USA), centrifuging the samples at 6000 rpm for 1 min in an Eppendorf 5424 Centrifuge. Libraries were prepared with the SQK-LSK109 kit (Oxford Nanopore, Oxford, UK) according to the manufacturer’s manual (version: GDE_9063_v109_revB_23May2018) with the following modifications: (1.) During DNA repair and end-prep, we used nuclease-free water instead of DNA CS; (2.) the incubation time of the end-prep reaction was 15 min (instead of 5 min) at 65°C; (3.) mixing was always done by pipetting; (4.) we used 80% ethanol (instead of 70%) for magnetic bead-based cleanup; (5.) we increased the time for DNA-binding to the magnetic beads to 15-20 min; (6.) the magnetic beads were air-dried for 1-2 min; (7.) after the adapter ligation, we carried out the clean-up with 45 µl Ampure XP beads (instead of 40 µl; Beckman Coulter, Brea, CA, USA); (8.) when preparing the library for loading onto the flow cell, we used 14 µl (instead of 12 µl) of the DNA library. To enrich for long fragments after adapter ligation, we used the L Fragment Buffer (LFB). After both the end-prep and adapter ligation steps, we quantified DNA using a Qubit 4.0 fluorometer.

*Oxford Nanopore whole genome amplification*

For the small species *Agapetus fuscipes* and *Agraylea sexmaculata*, gDNA obtained from a single individual was insufficient for direct library preparation for Nanopore sequencing. Therefore, we performed whole genome amplification (WGA) following the “Premium whole genome amplification protocol (SQK-LSK109), version 23 May 2018”. First, we diluted gDNA samples with 1x TE buffer to obtain a DNA input of ca. 2 ng. We mixed 5 µl normalized gDNA with 5 µl reconstituted DLB buffer (from the Repli-g Mini Kit, Qiagen, Hilden, Germany) and incubated for 3 min at room temperature. Then, 10 µl of reconstituted Stop Solution were added and mixed by pipetting. We added this DNA reaction mixture to a 0.2 ml PCR tube that contained 29 µl Repli-g Reaction buffer and 1 µl Repli-g DNA Polymerase. After mixing by pipetting, the reaction was incubated for 16 h at 30°C, followed by 3 min at 65°C in a Master Cycler (Eppendorf, Hamburg, Germany). We purified the reactions using 90 µl Ampure XP beads (Beckman and Coulter, Brea, U.S.A.). The beads were washed only once with 80% freshly prepared ethanol. We resuspended the DNA in 50 µl nuclease-free water and quantified it with the Qubit BR kit. Afterwards, we mixed 1.5 µg of the amplified DNA with 3 µl NEBuffer 2, 1.5 µl T7 endonuclease I (New England Biolabs, Ipswich, U.S.A.) and water to achieve a reaction volume of 30 µl. This reaction mixture was incubated for 15 min at 37°C. We cleaned up reactions using 35 µl of the pre-made custom bead suspension (made from Ampure XP beads following the protocol). The beads were washed twice with 80% ethanol and the DNA resuspended in 50 µl nuclease-free water. The DNA was quantified with the Qubit BR kit. Subsequently, the library preparation continued with the end-prep and FFPE repair following the protocol for the SQK-LSK109 library preparation with the addition of a barcoding step for multiplexed sequencing using the EXP-NBD104 Native Barcoding Expansion kit (Oxford Nanopore).

We sequenced each Oxford Nanopore library on a single flow cell using the MinION portable DNA sequencer. For basecalling Nanopore reads from their raw .fast5 files, we used Poretools v0.6.0 [7] or Guppy Basecalling Software v2.3.1. Reads resulting from the whole genome amplification protocol were demultiplexed using guppy_barcoder (Guppy Basecalling Software v2.3.1: https://nanoporetech.com/nanopore-sequencing-data-analysis) with --barcode_kits "EXP-NBD104". After trimming adapters with Porechop v.2.0.4 (https://github.com/rrwick/Porechop) using default parameters, we used FASTQ Screen like tools (<https://github.com/schellt/fqs-tools>) to filter out potential contaminants. For this purpose, we mapped the long reads to a custom-made database with minimap2 –x map-ont. The database contains viral, bacteria and human sequences, as well as potential parasites of Trichoptera (Ascogregaria, Gregarina, Mermithidae) and Trichoptera genomes (as positive control). We created an ID list with awk and extracted the screening info with the paf2fqs.pl script of FASTQ Screen like tools (https://github.com/schellt/fqs-tools). Screening info was extracted with the paf2fqs.pl script from FastQ Screen like tools using the paf. file obtained from minimap2 and the previously created ID list. We kept reads without hits and reads that mapped to Trichoptera genomes with subseq command of seqtk 1.3-r106 (https://github.com/lh3/seqtk).

*PacBio sequencing*

Two species, *Micrasema minimum* and *Agraylea sexmaculata,* were sequenced with PacBio. We constructed a SMRTbell library following the instructions of the SMRTbell Express Prep kit v2.0 with low DNA Input Protocol (Pacific Biosciences, Menlo Park, CA). Following ligation with T-overhang SMRTbell adapters at 20°C overnight, we purified the SMRTbell library with an AMPure PB bead clean up step with 0.45X volume of AMPure PB beads. To remove short SMRTbell templates < 3kb, we performed a size-selection step with AMPure PB Beads. For this purpose, we diluted the AMPure PB beads with elution buffer (40% volume/volume) and added a 2.2X volume to the DNA sample. We performed one SMRT cell sequencing run each on the Sequel System II with Sequel II Sequencing Kit 2.0. CLR mode was run for 30 hour movie time with NO pre-extension and Software SMRTLINK 8.0. After obtaining a subreads.bam file containing automatically trimmed reads, we used bam2fastq (https://github.com/jts/bam2fastq) to extract and save reads in fastq format. To filter out contaminations, we mapped reads to our custom database with minimap2 -x map-pb and followed the steps described above.

**Supplementary note 3: Assembly strategies**

*Long-read assembly with subsequent short-read polishing*

In detail, we first conducted a long-read assembly of the Oxford Nanopore Technology sequencing reads using a fuzzy Bruijn graph approach with wtdbg2 v2.4 [8], following a correction with long-reads by re-mapping these to the assembly with Minimap2 v14 [9] and polishing with Racon v1.3.1 [10]. We then used nanopolish 0.11.1 [11] to further improve the consensus accuracy of the assembly. For this purpose, we used nanopolish index to create an index readdb file that links read ids with their signal-level data in the FAST5 files. To compute a new consensus sequence of our draft assembly, we first aligned the long-reads to the draft assembly with minimap and sorted alignments with samtools. Subsequently, we ran a consensus algorithm with nanopolish variants (--consensus --min-candidate-frequency 0.1) on our draft assembly. We used nanopolish_makerange.py to split our draft genome assembly into 50kb segments, to run this step on several parts of the assembly in parallel. We generated the polished genome in fasta format with nanopolish vcf2fasta. To obtain an improved representation of the genome based on the short-read data, we used Pilon v1.22 [12]to further improve the assembly. Therefore, we mapped the preprocessed Illumina reads to the „nanopolished“ assembly with bwa mem and sorted the read alignments by leftmost coordinates using sort options of SAMtools v1.9 [13]. We used Pilon v1.22 [14], option --fix indels) to identify inconsistencies between the input genome and the evidence in the reads. We used purge_dups 1.2.3 [15] to purge haplotigs and overlaps in an assembly based on read depth.

### Hybrid assemblies

#### *DeBruijn graph and overlap-layout-consensus assembly using MaSuRCA*

We conducted a *de novo* hybrid assembly with the raw Illumina data together with the long-reads using MaSuRCA v.3.1.1 [137-138]. In the config file for each run, we specified the insert size and a standard deviation (10% of insert size) for the Illumina reads, as well as jellyfish hash size (estimated_genome_size*~long-read coverage). All other parameters were left as defaults. We used purge_dups 1.2.3 to purge haplotigs and overlaps in an assembly based on read depth.

*SPAdes hybrid assembly*

For datasets with low long-read coverage, we conducted a hybrid assembly using SPAdes v3.12.0 [16] with the preprocessed Illumina and Oxford Nanopore reads. The option --cov-cutoff auto was applied. SPAdes contigs ≥ 500 bp were further scaffolded with the long-reads using SLR (https://github.com/luojunwei/SLR) and gaps were closed with TGS-Gapcloser (https://github.com/BGI-Qingdao/TGS-GapCloser).

*Short-read assembly*

For Illumina only datasets, we conducted short-read assembly with the preprocessed Illumina reads using SPAdes v3.13.1 [17] with the option *--cov-cutoff auto.*

**Supplementary note 4: Functional annotation of protein coding genes**

The annotation of the genomes resulted in the prediction of 6,413 to 12,927 proteins. Most of the annotated proteins (94.4% - 98.8%) showed significant sequence similarity to entries in the NCBI nr database. GO Distributions were similar to previously annotated caddisfly genomes. Specifically, the major biological processes were cellular and metabolic processes. Catalytic activity was the largest subcategory in molecular function. Regarding the cellular component category, most genes were assigned to the cell subcategory or to the membrane subcategory.

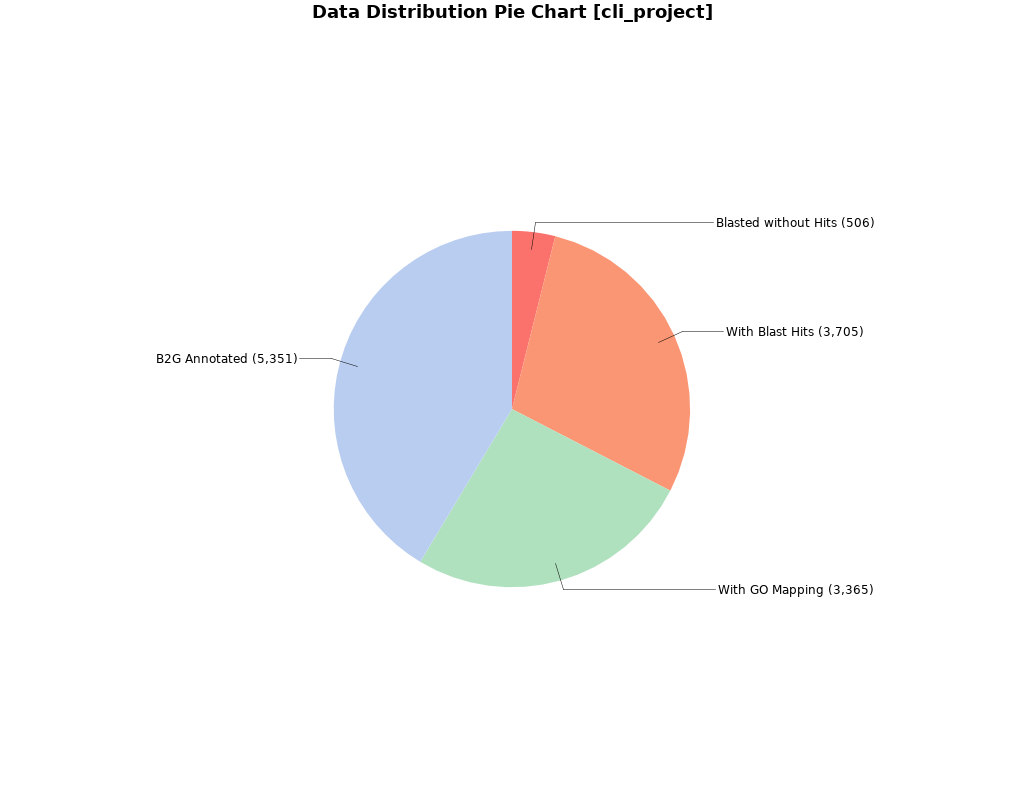

**Fig. S1. Blast2GO Annotation Results of *Drusus annulatus*.** Pie charts showing the percentage of proteins with functional Blast2GO annotations, verified by BLAST and mapped to GO terms.

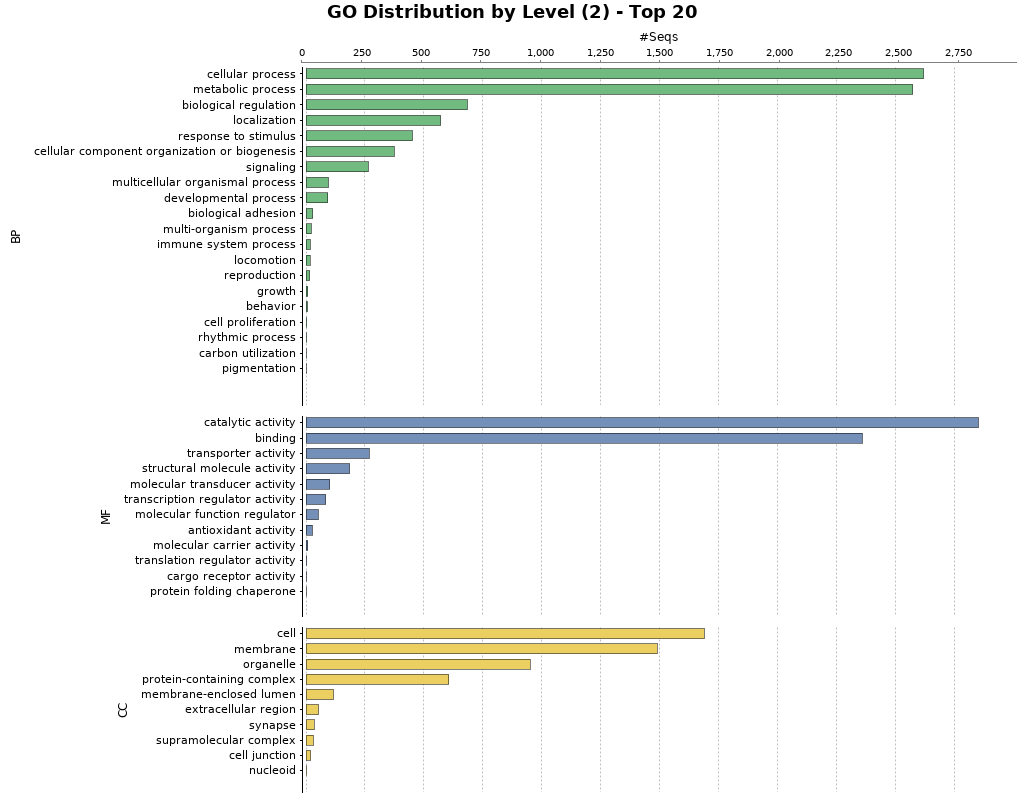

**Fig. S2. Blast2GO Functional Annotation for *Drusus annulatus.*** Barplot showing GO terms characterized by biological process, molecular function, and cellular component. Barplots are grouped by biological process (BP), molecular function (MF), and cellular component (CC).

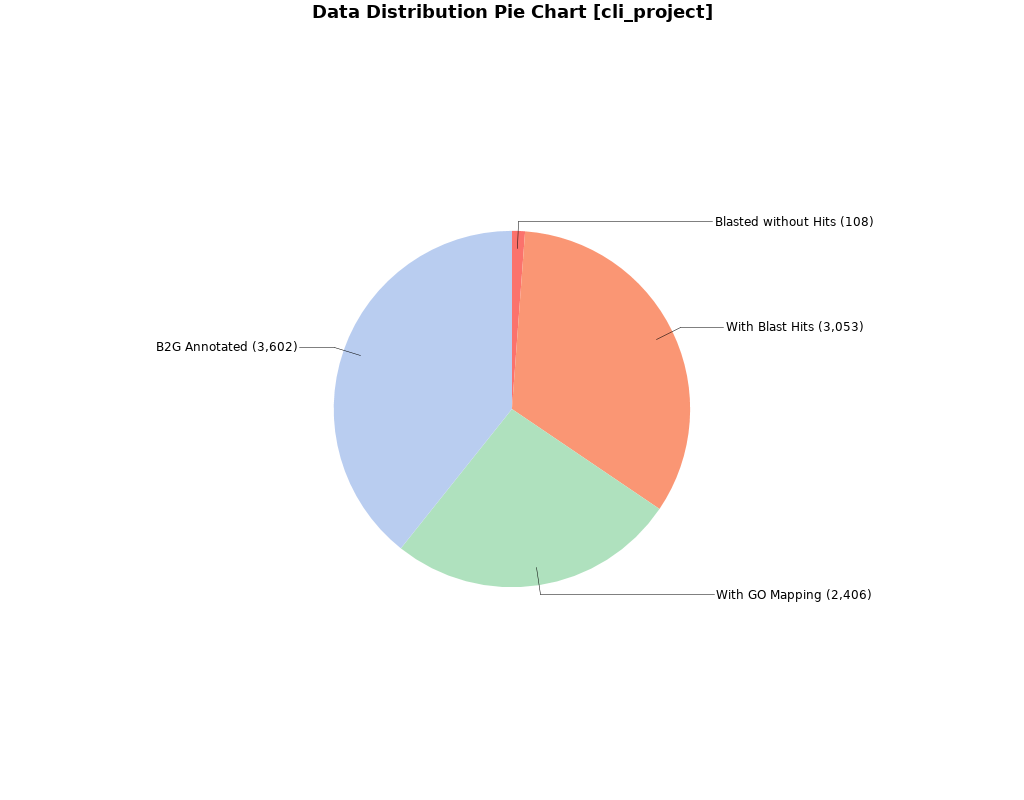

**Fig. S3. Blast2GO Annotation Results of *Agraylea sexmaculata*.** Pie charts showing the percentage of proteins with functional Blast2GO annotations, verified by BLAST and mapped to GO terms.

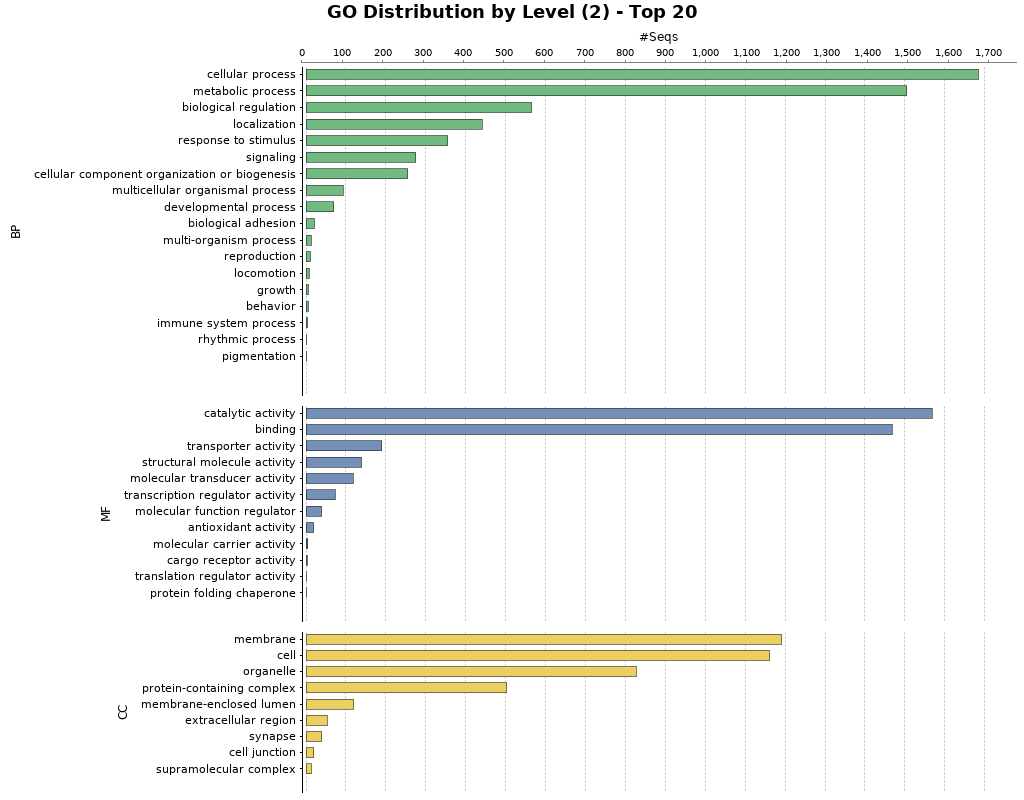

**Fig. S4. Blast2GO Functional Annotation for *Agraylea sexmaculata.*** Barplot showing GO terms characterized by biological process, molecular function, and cellular component. Barplots are grouped by biological process (BP), molecular function (MF), and cellular component (CC).

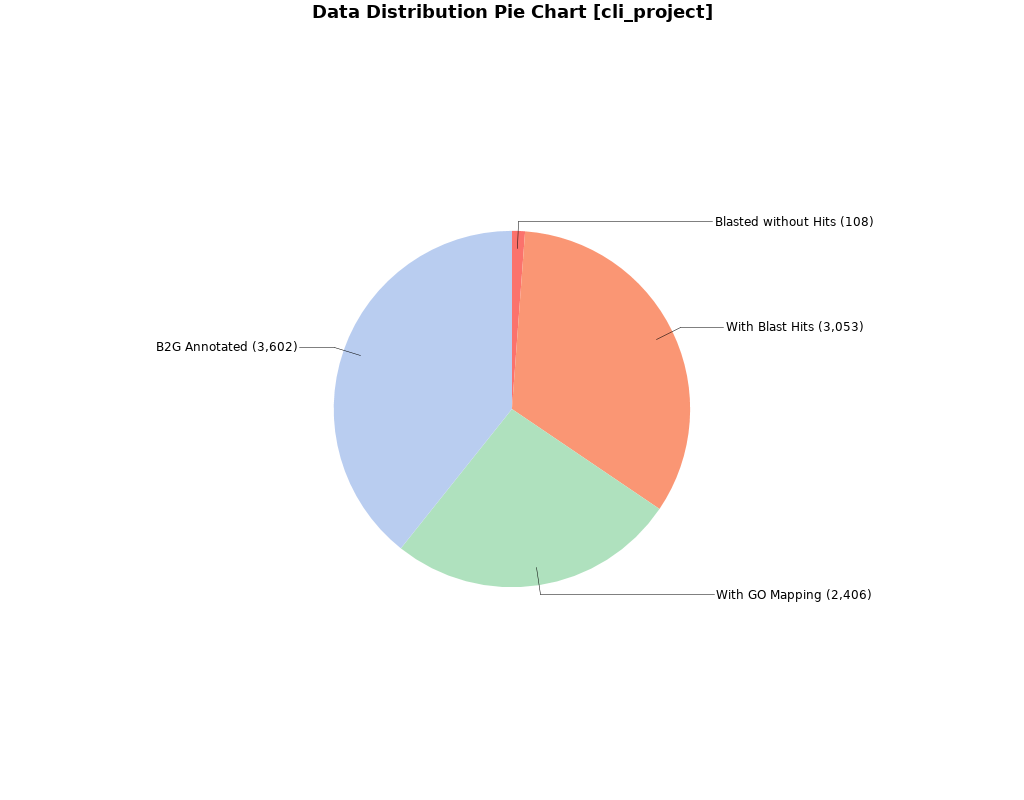

**Fig. S5. Blast2GO Annotation Results of *Glossosoma conforme*.** Pie charts showing the percentage of proteins with functional Blast2GO annotations, verified by BLAST and mapped to GO terms.

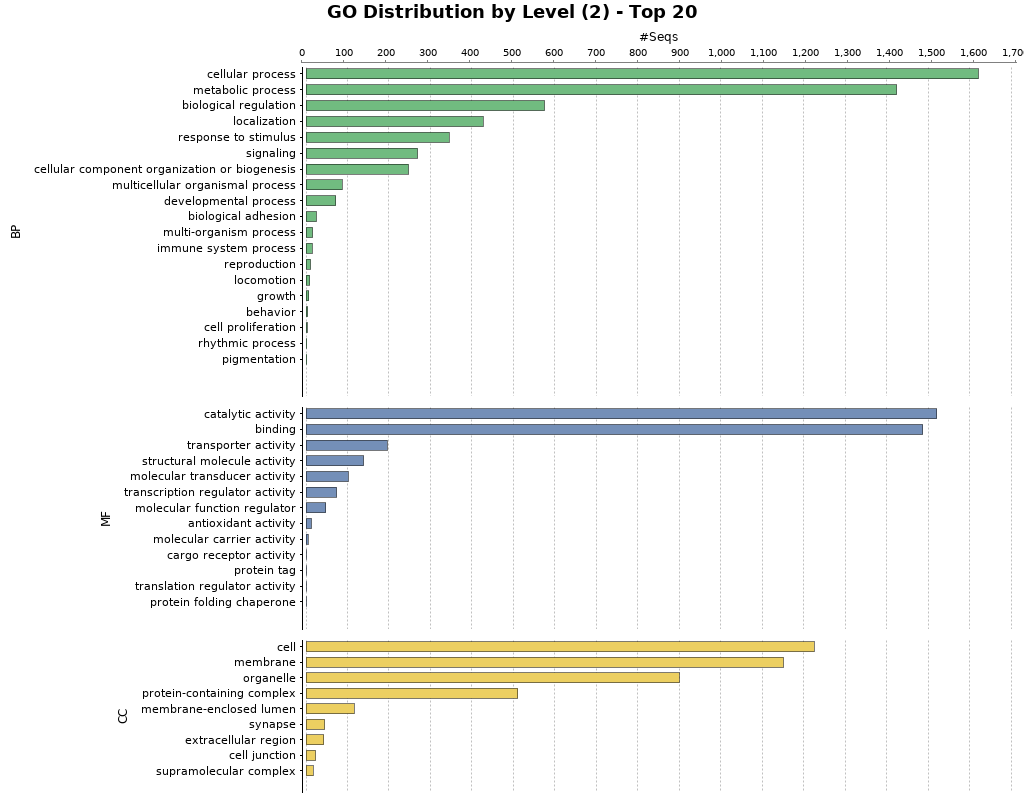

**Fig. S6. Blast2GO Functional Annotation for *Glossosoma conforme.*** Barplot showing GO terms characterized by biological process, molecular function, and cellular component. Barplots are grouped by biological process (BP), molecular function (MF), and cellular component (CC).

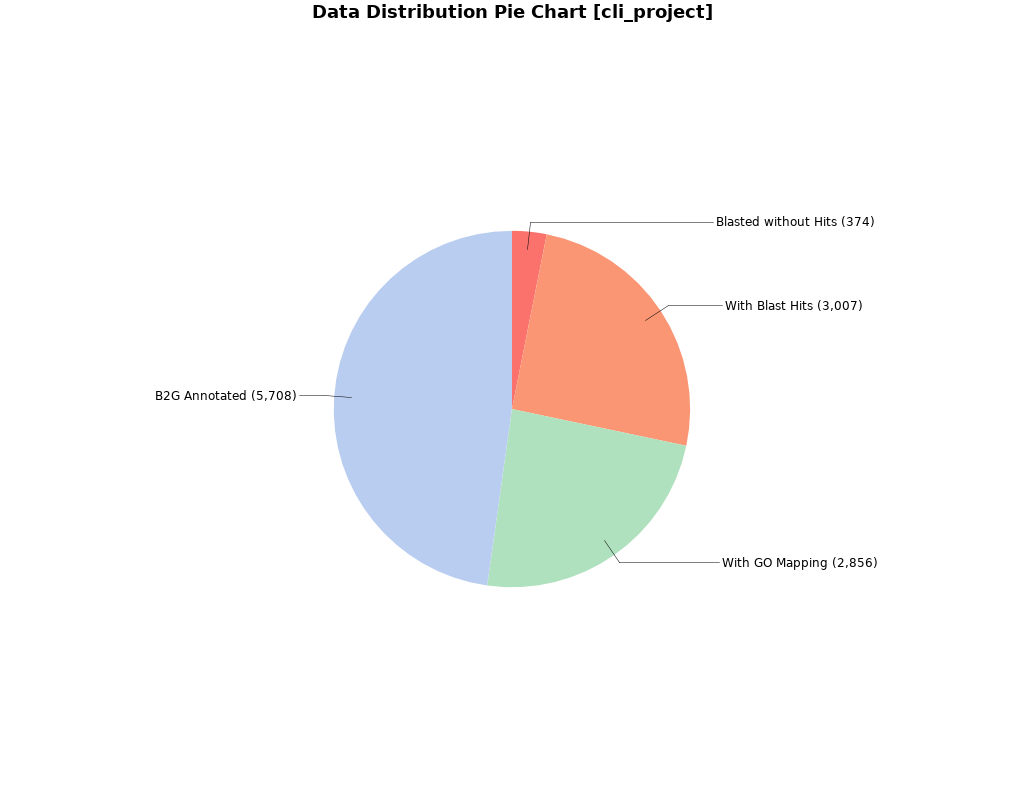

**Fig. S7. Blast2GO Annotation Results of *Halesus radiatus*.** Pie charts showing the percentage of proteins with functional Blast2GO annotations, verified by BLAST and mapped to GO terms.

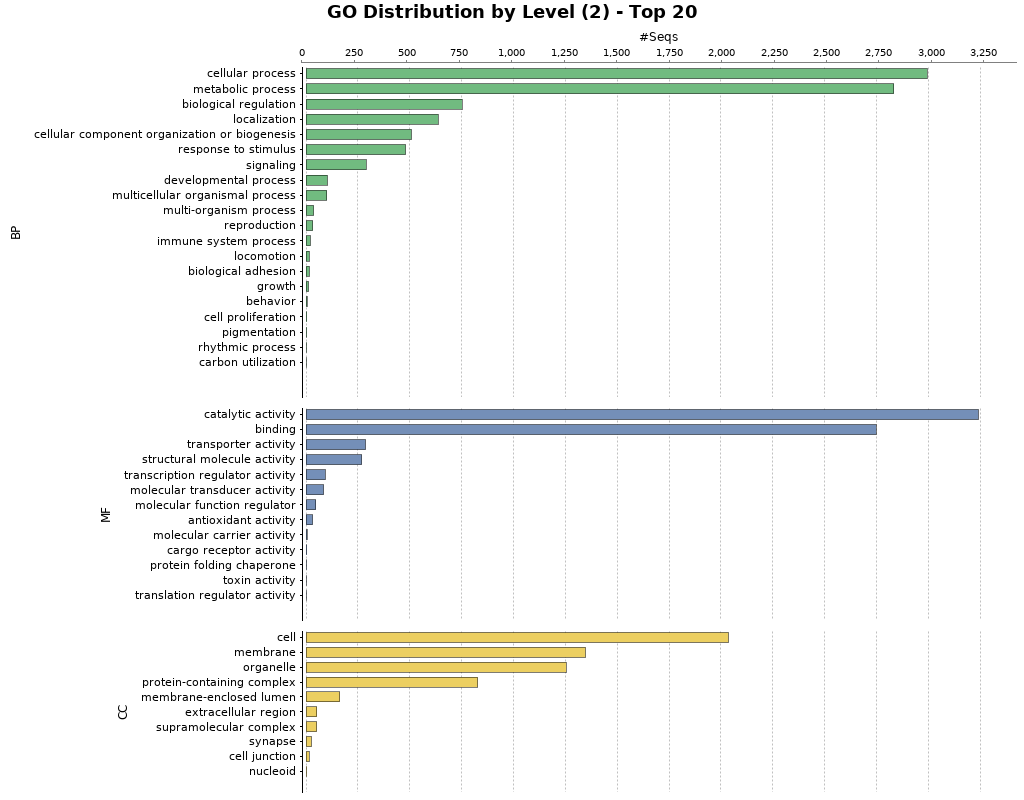

**Fig. S8. Blast2GO Functional Annotation for *Halesus radiatus.*** Barplot showing GO terms characterized by biological process, molecular function, and cellular component. Barplots are grouped by biological process (BP), molecular function (MF), and cellular component (CC).

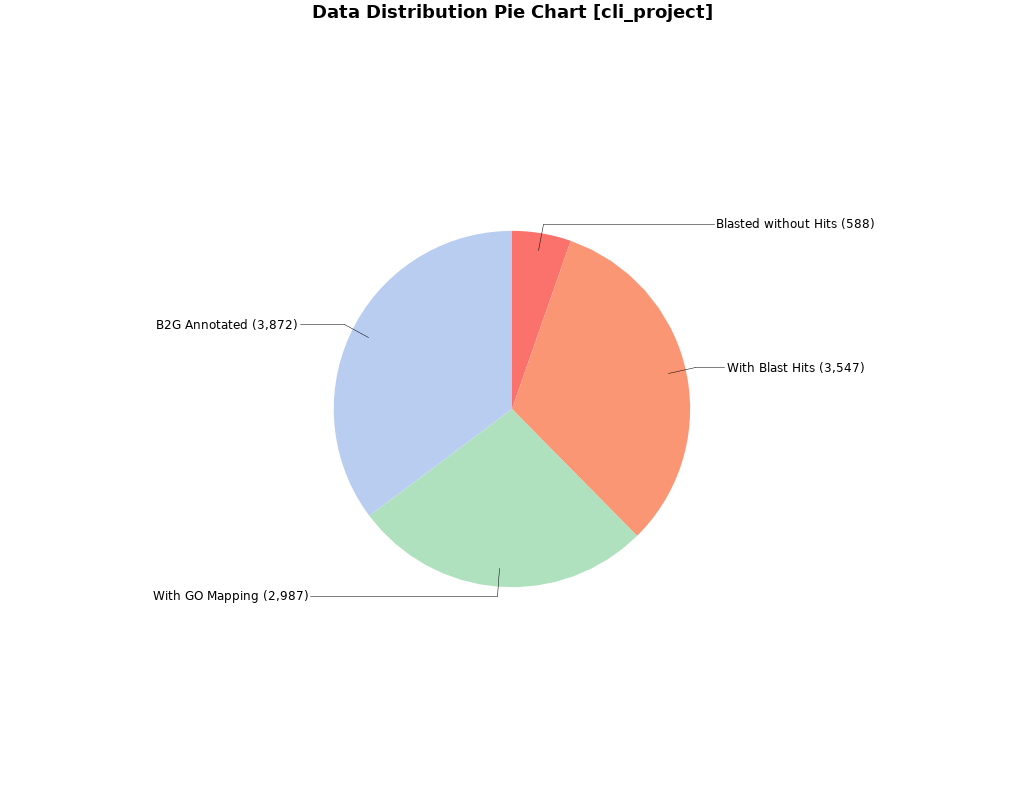

**Fig. S9. Blast2GO Annotation Results of *Himalopsyche phryganeae*.** Pie charts showing the percentage of proteins with functional Blast2GO annotations, verified by BLAST and mapped to GO terms.

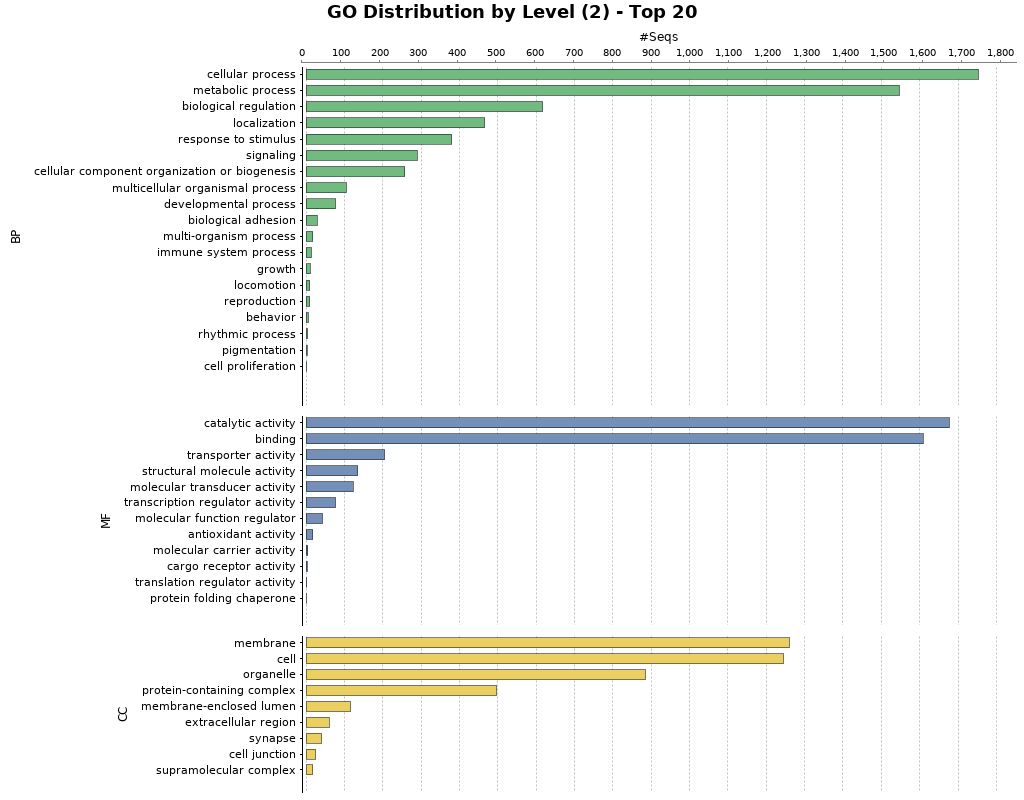

**Fig. S10. Blast2GO Functional Annotation for *Himalopsyche phryganeae.*** Barplot showing GO terms characterized by biological process, molecular function, and cellular component. Barplots are grouped by biological process (BP), molecular function (MF), and cellular component (CC).

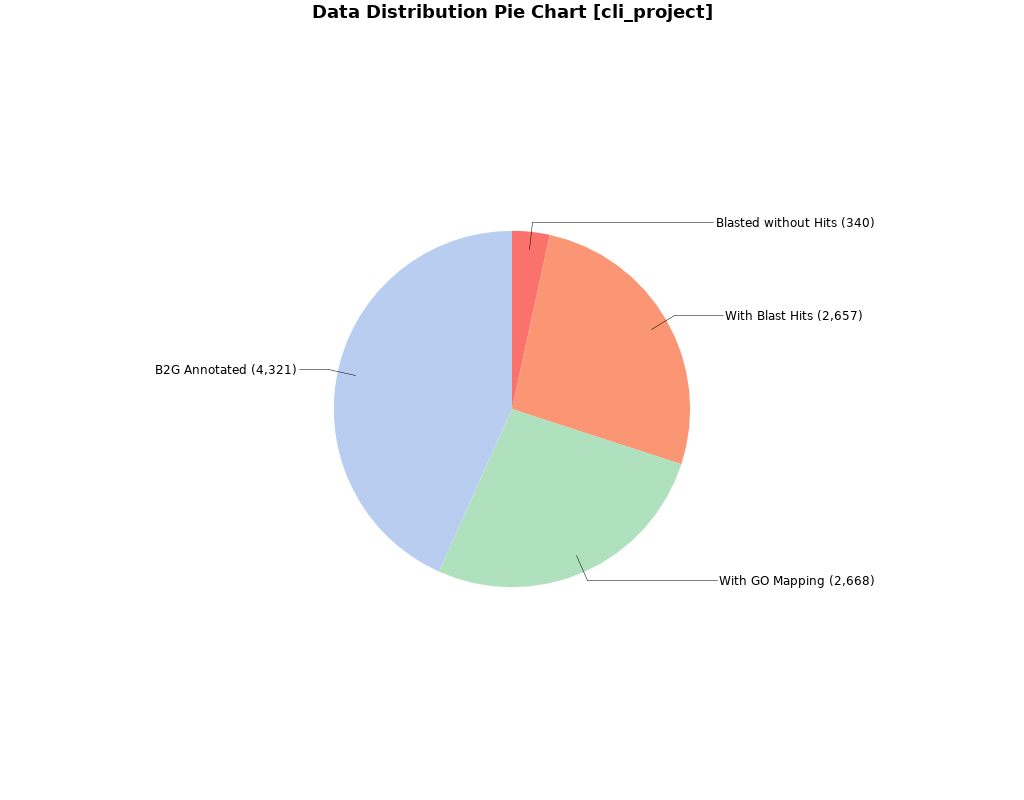

**Fig. S11. Blast2GO Annotation Results of *Lepidostoma basale*.** Pie charts showing the percentage of proteins functional Blast2GO annotations, verified by BLAST and mapped to GO terms.

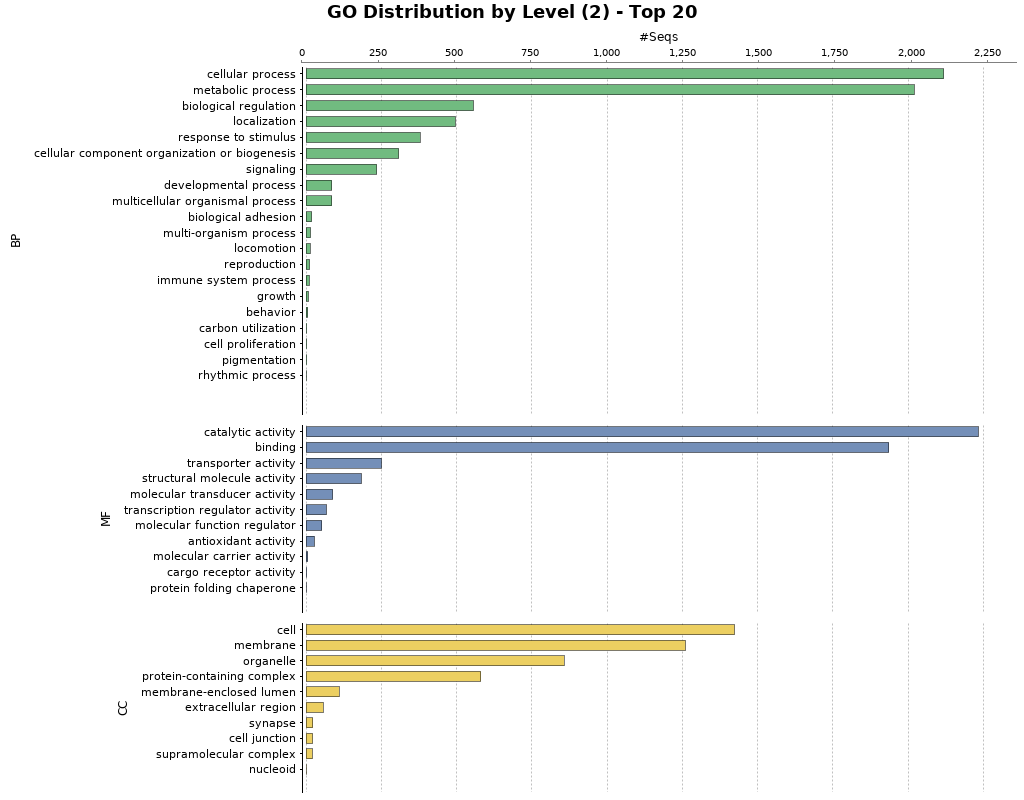

**Fig. S12. Blast2GO Functional Annotation for *Lepidostoma basale.*** Barplot showing GO terms characterized by biological process, molecular function, and cellular component. Barplots are grouped by biological process (BP), molecular function (MF), and cellular component (CC).

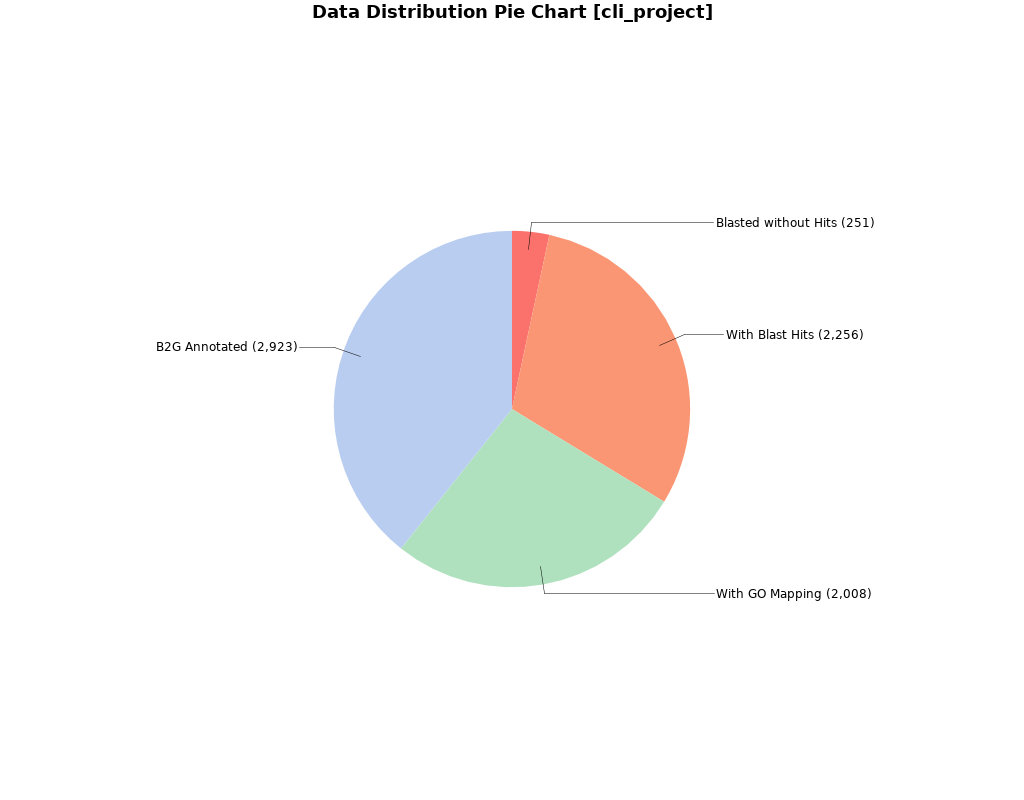

**Fig. S13. Blast2GO Annotation Results of *Micrasema longulum ML3.*** Pie charts showing the percentage of proteins with functional Blast2GO annotations, verified by BLAST and mapped to GO terms.

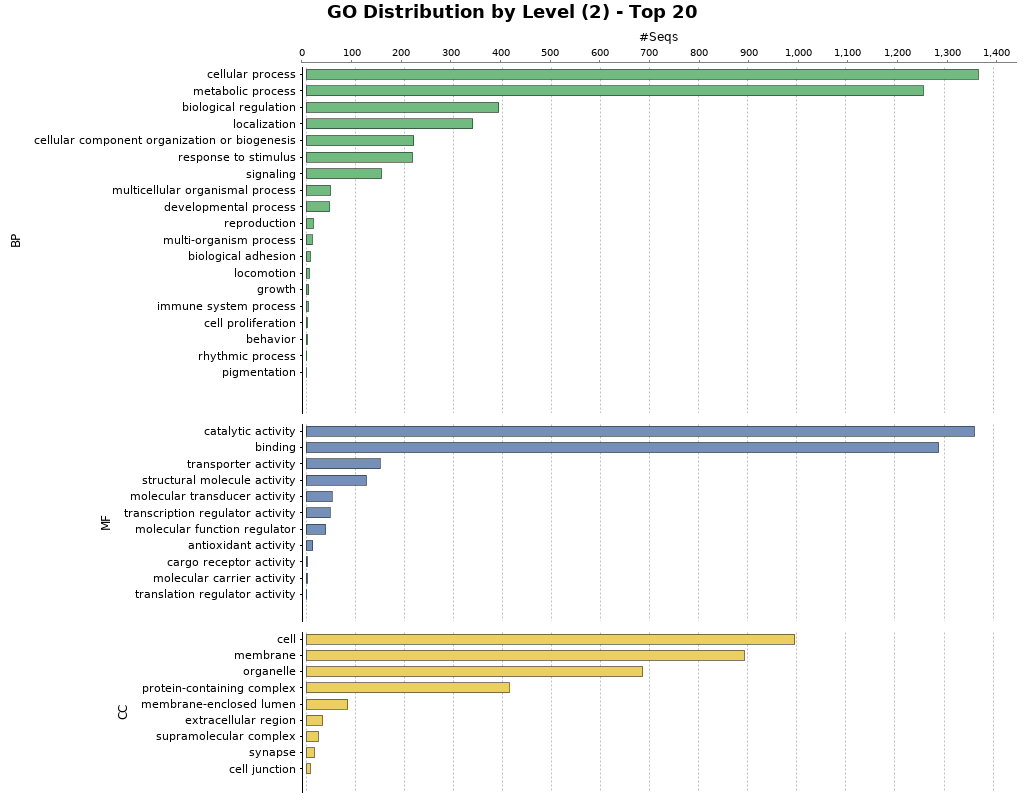

**Fig. S14. Blast2GO Functional Annotation for *Micrasema longulum ML3.*** Barplot showing GO terms characterized by biological process, molecular function, and cellular component. Barplots are grouped by biological process (BP), molecular function (MF), and cellular component (CC).

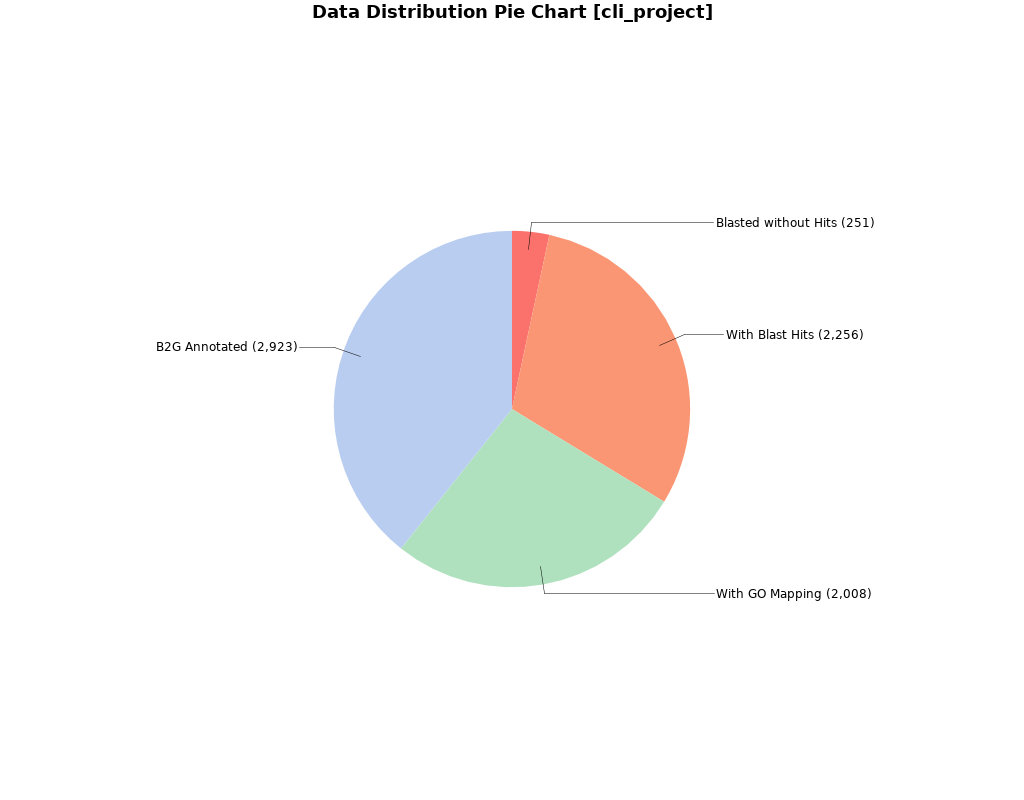

**Fig. S15. Blast2GO Annotation Results of *Micrasema minimum*.** Pie charts showing the percentage of proteins with functional Blast2GO annotations, verified by BLAST and mapped to GO terms.

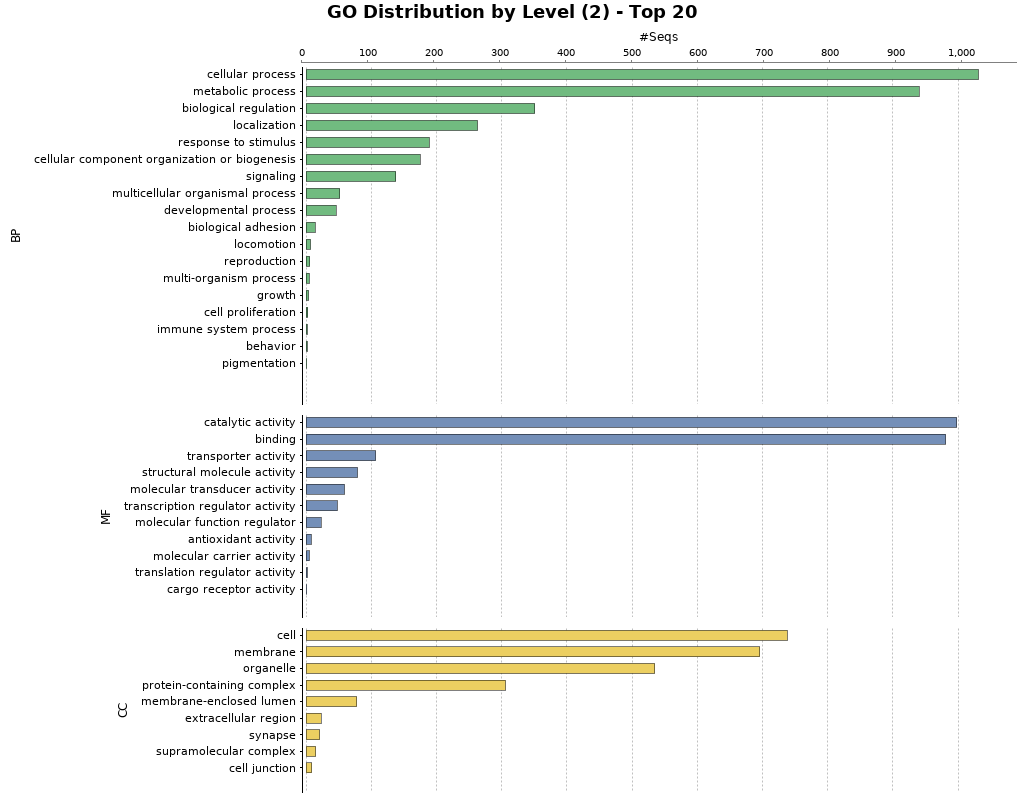

**Fig. S16. Blast2GO Functional Annotation for *Micrasema minimum.*** Barplot showing GO terms characterized by biological process, molecular function, and cellular component. Barplots are grouped by biological process (BP), molecular function (MF), and cellular component (CC).

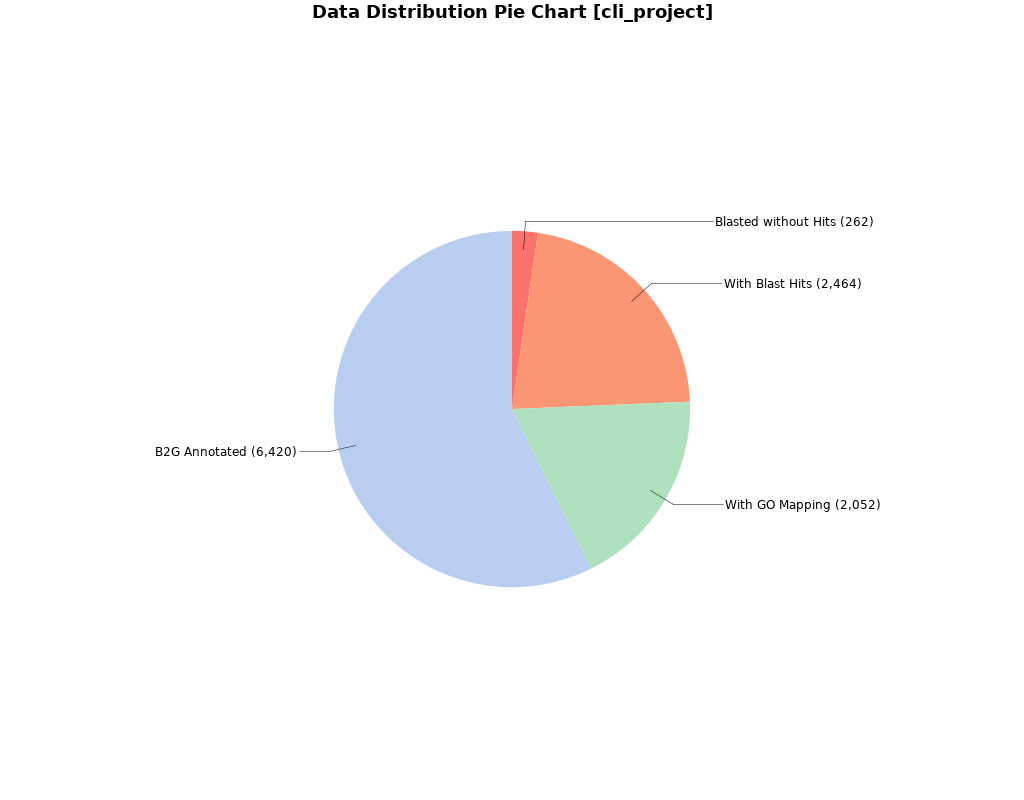

**Fig. S17. Blast2GO Annotation Results of *Micropterna sequax*.** Pie charts showing the percentage of proteins in Micropterna sequax with functional Blast2GO annotations, verified by BLAST and mapped to GO terms.

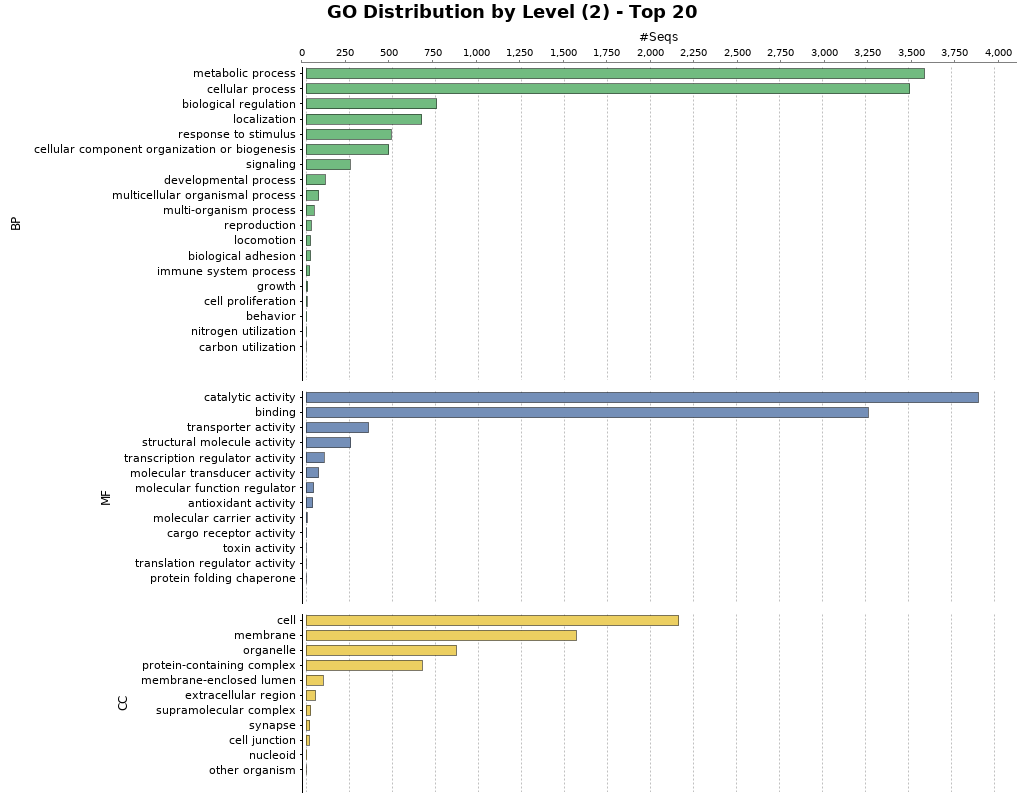

**Fig. S18. Blast2GO Functional Annotation for *Micropterna sequax.*** Barplot showing GO terms characterized by biological process, molecular function, and cellular component. Barplots are grouped by biological process (BP), molecular function (MF), and cellular component (CC).

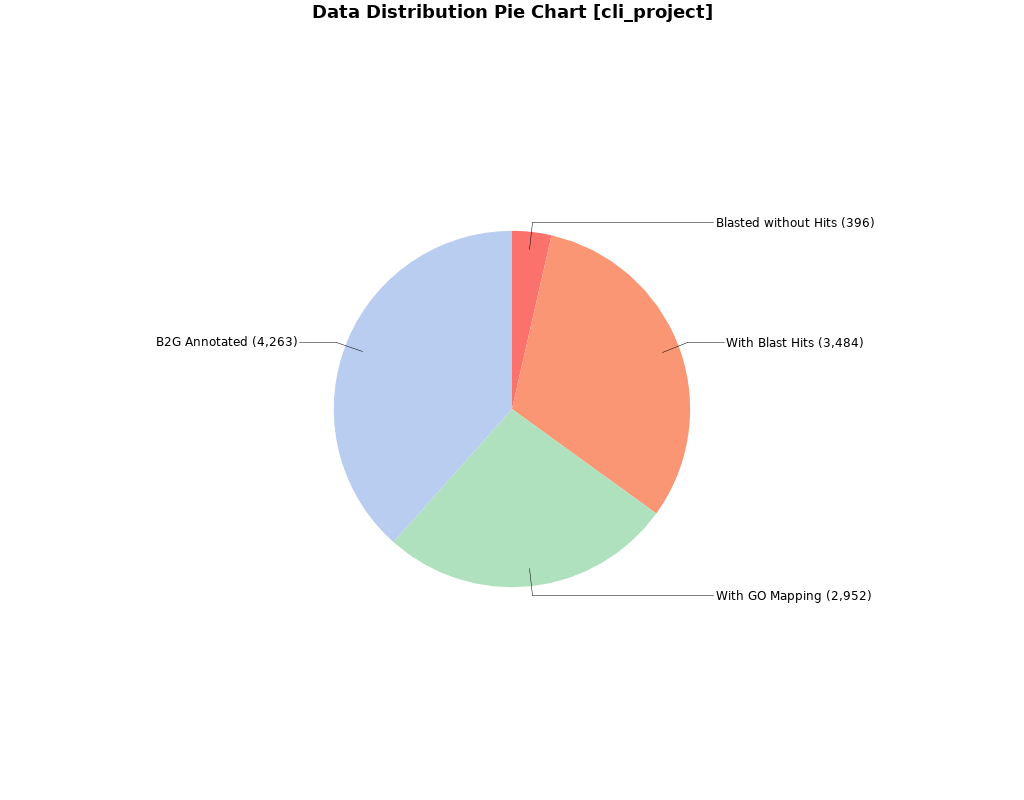

**Fig. S19. Blast2GO Annotation Results of *Odontocerum albicorne*.** Pie charts showing the percentage of proteins with functional Blast2GO annotations, verified by BLAST and mapped to GO terms.

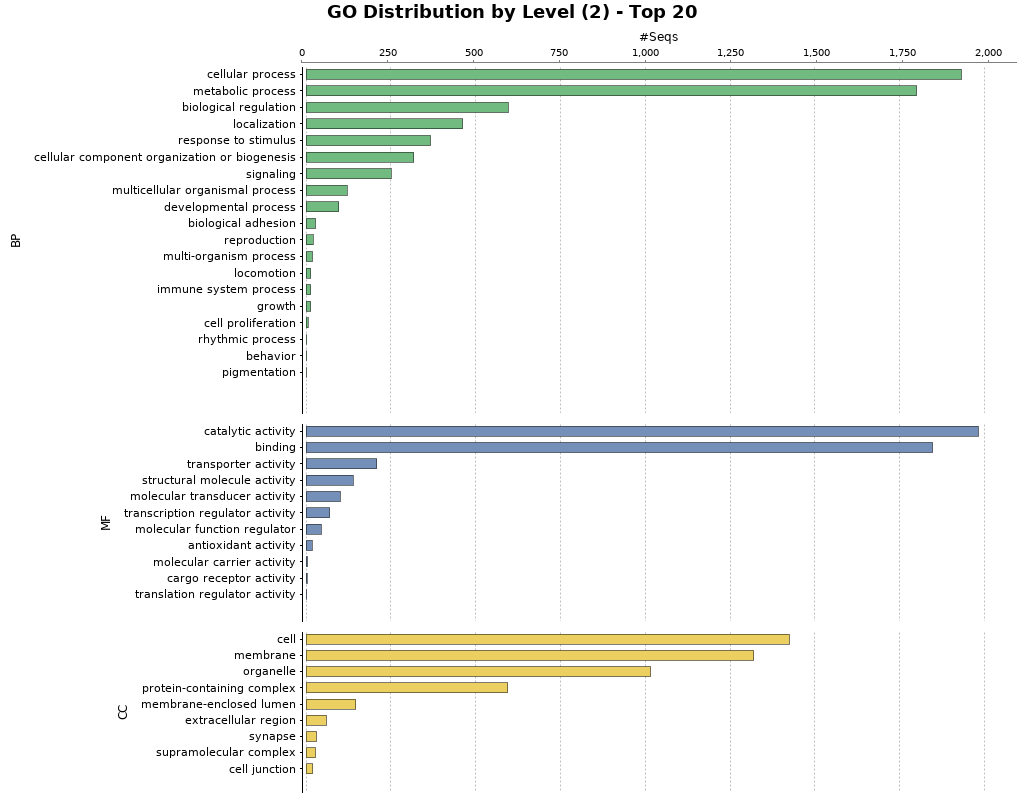

**Fig. S20. Blast2GO Functional Annotation for *Odontocerum albicorne.*** Barplot showing GO terms characterized by biological process, molecular function, and cellular component. Barplots are grouped by biological process (BP), molecular function (MF), and cellular component (CC).

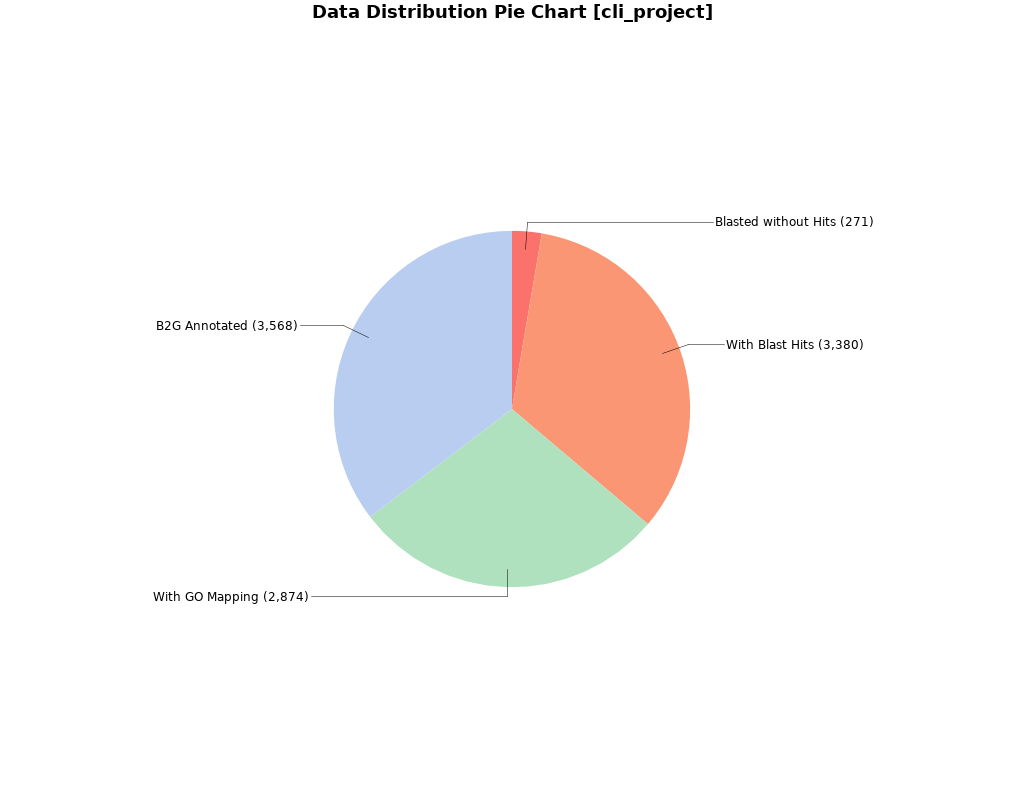

**Fig. S21. Blast2GO Annotation Results of *Parapsyche elsis*.** Pie charts showing the percentage of proteins with functional Blast2GO annotations, verified by BLAST and mapped to GO terms.

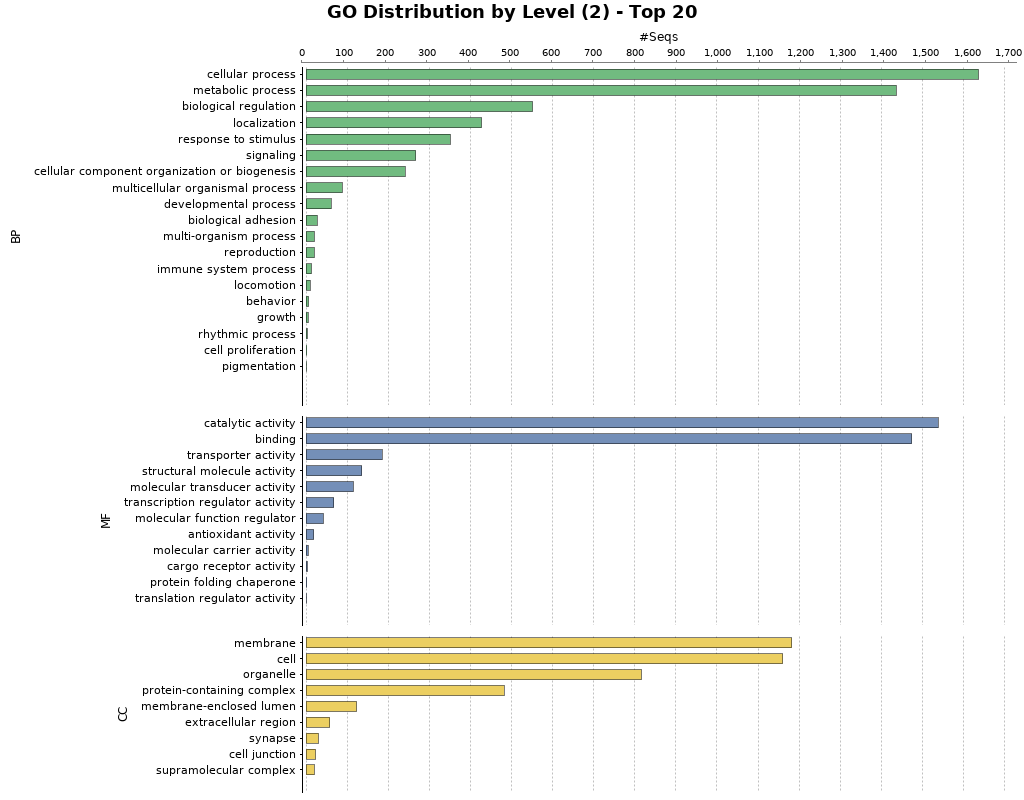

**Fig. S22. Blast2GO Functional Annotation for *Parapsyche elsis.*** Barplot showing GO terms characterized by biological process, molecular function, and cellular component. Barplots are grouped by biological process (BP), molecular function (MF), and cellular component (CC).

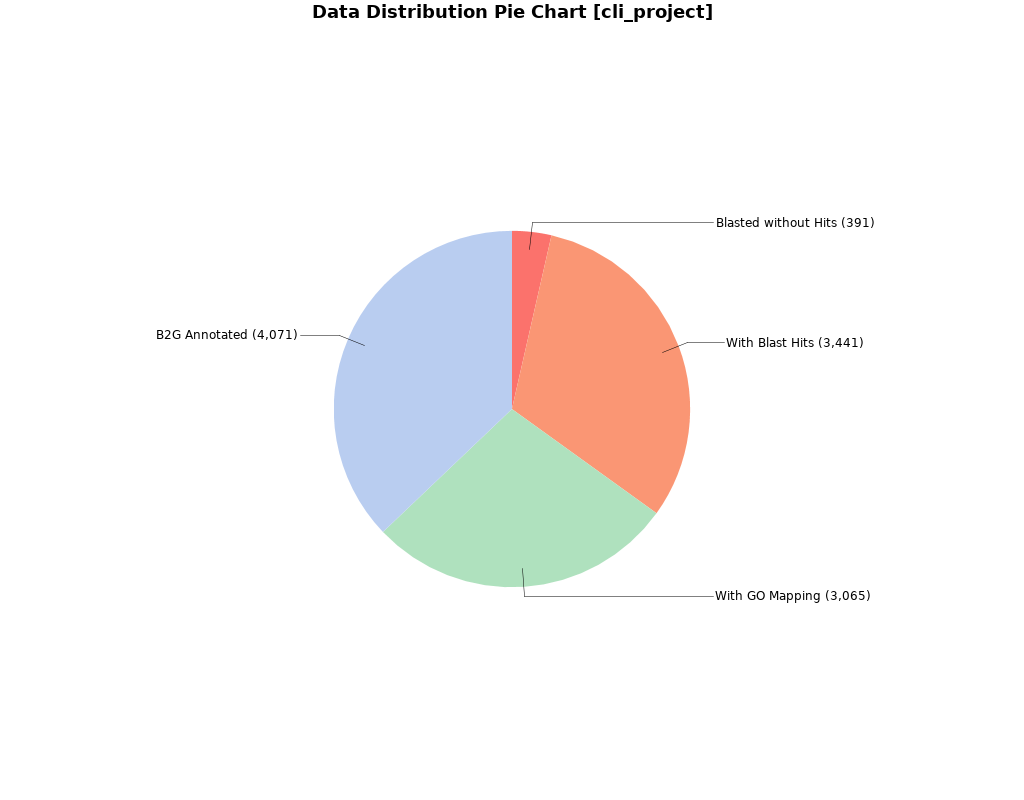

**Fig. S23. Blast2GO Annotation Results of *Philopotamus ludiferatus*.** Pie charts showing the percentage of proteins with functional Blast2GO annotations, verified by BLAST and mapped to GO terms.

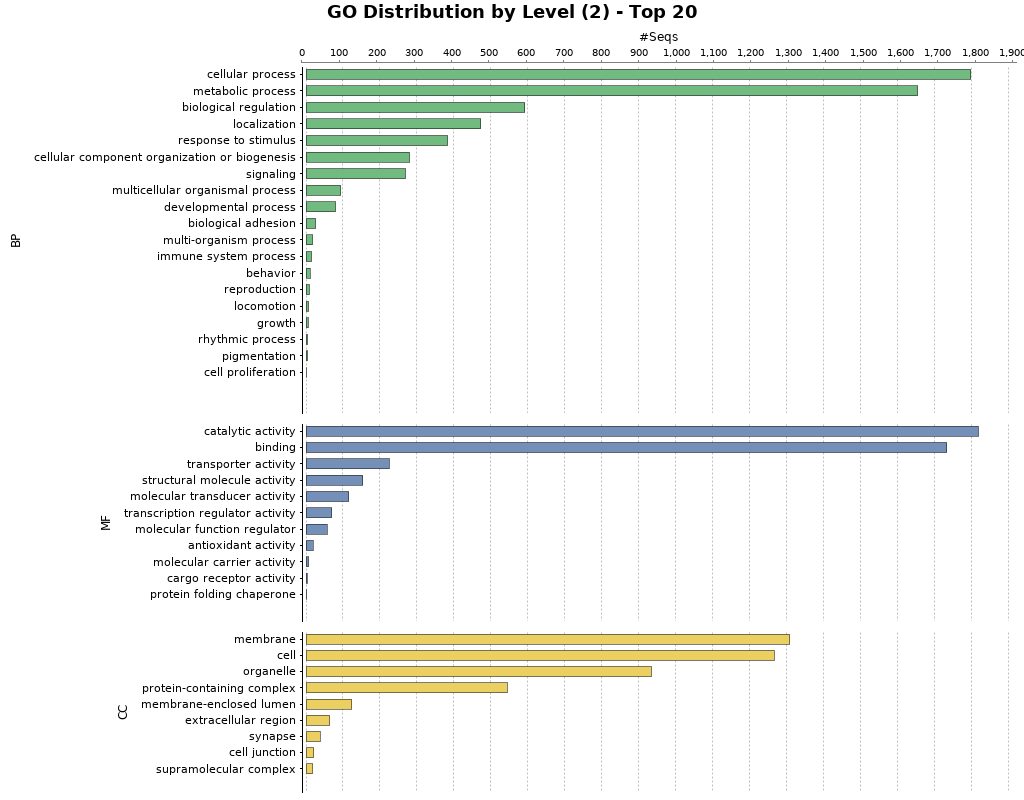

**Fig. S24. Blast2GO Functional Annotation for *Philopotamus ludiferatus.*** Barplot showing GO terms characterized by biological process, molecular function, and cellular component. Barplots are grouped by biological process (BP), molecular function (MF), and cellular component (CC).

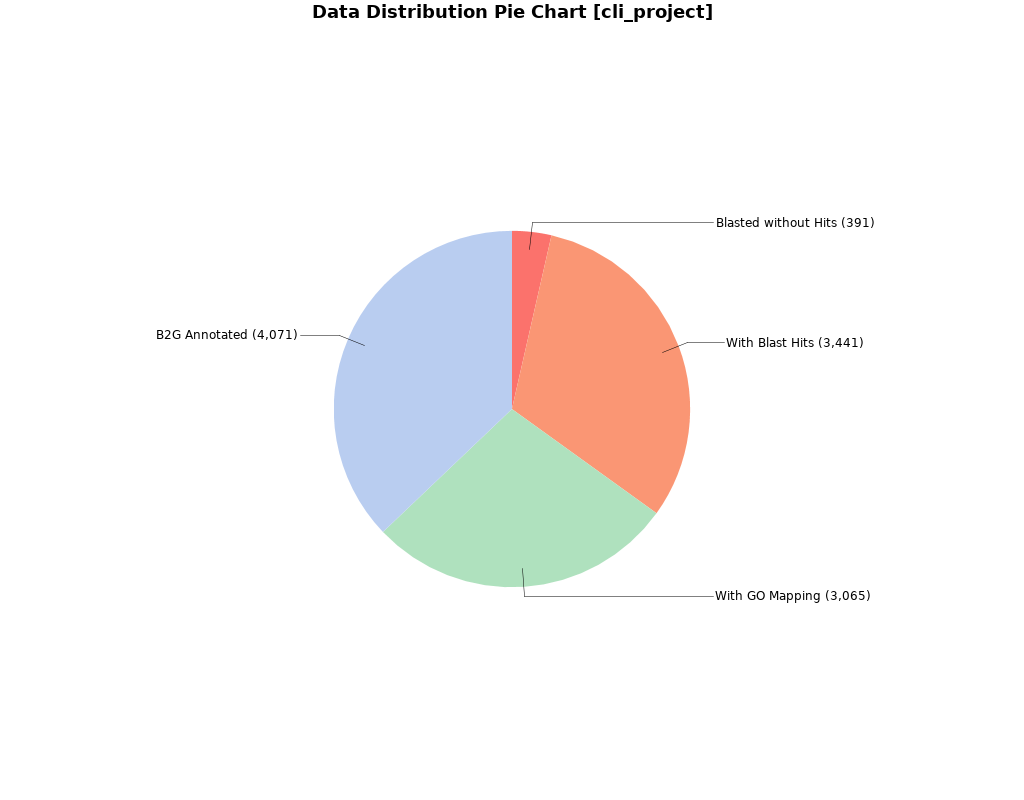

**Fig. S25. Blast2GO Annotation Results of *Rhyacophila brunneae*.** Pie charts showing the percentage of proteins with functional Blast2GO annotations, verified by BLAST and mapped to GO terms.

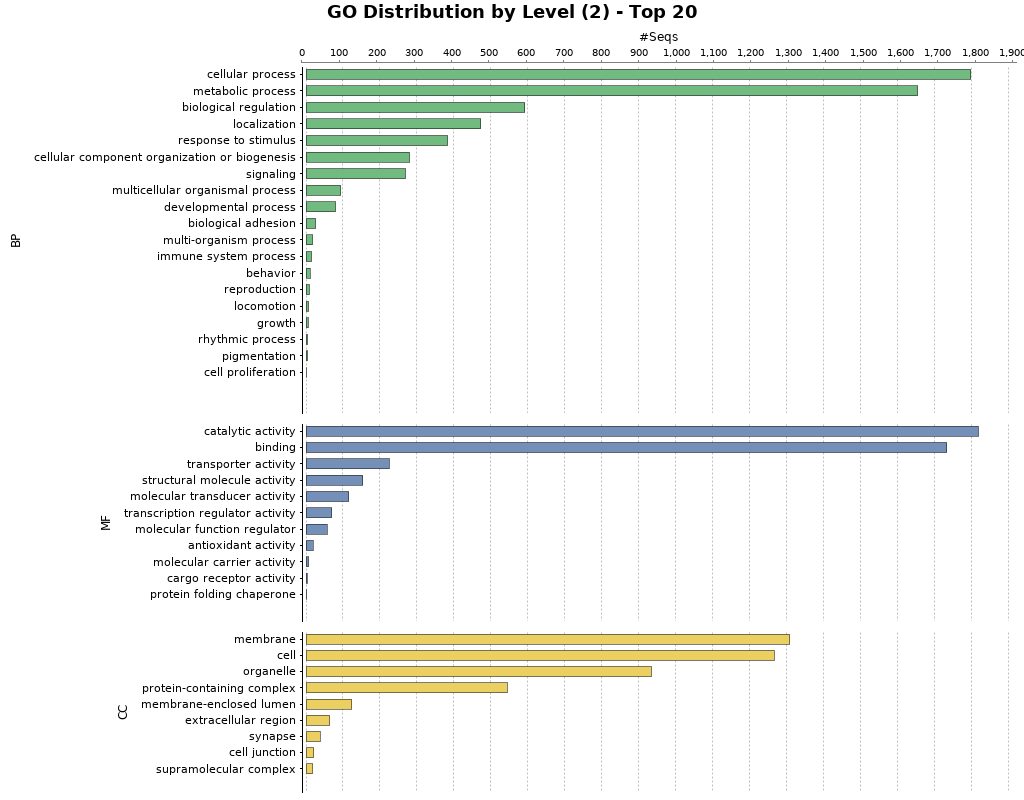

**Fig. S26. Blast2GO Functional Annotation for *Rhyacophila brunneae.*** Barplot showing GO terms characterized by biological process, molecular function, and cellular component. Barplots are grouped by biological process (BP), molecular function (MF), and cellular component (CC).

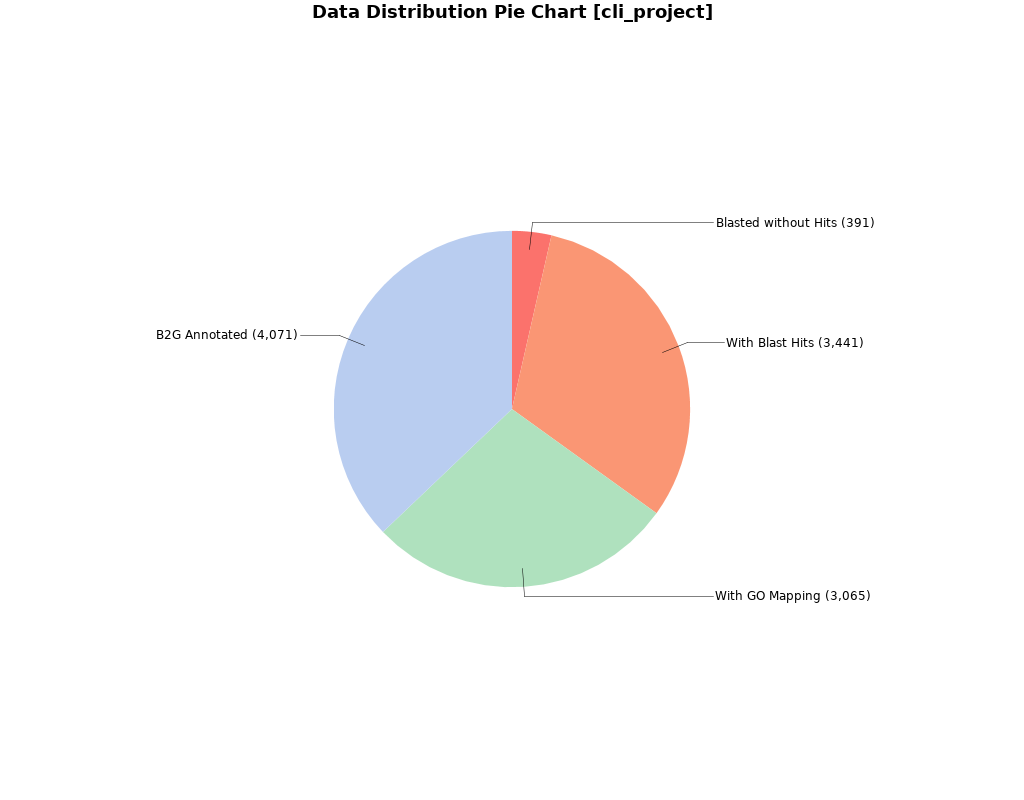

**Fig. S27. Blast2GO Annotation Results of *Rhyacophila evoluta HR1*.** Pie charts showing the percentage of proteins with functional Blast2GO annotations, verified by BLAST and mapped to GO terms.

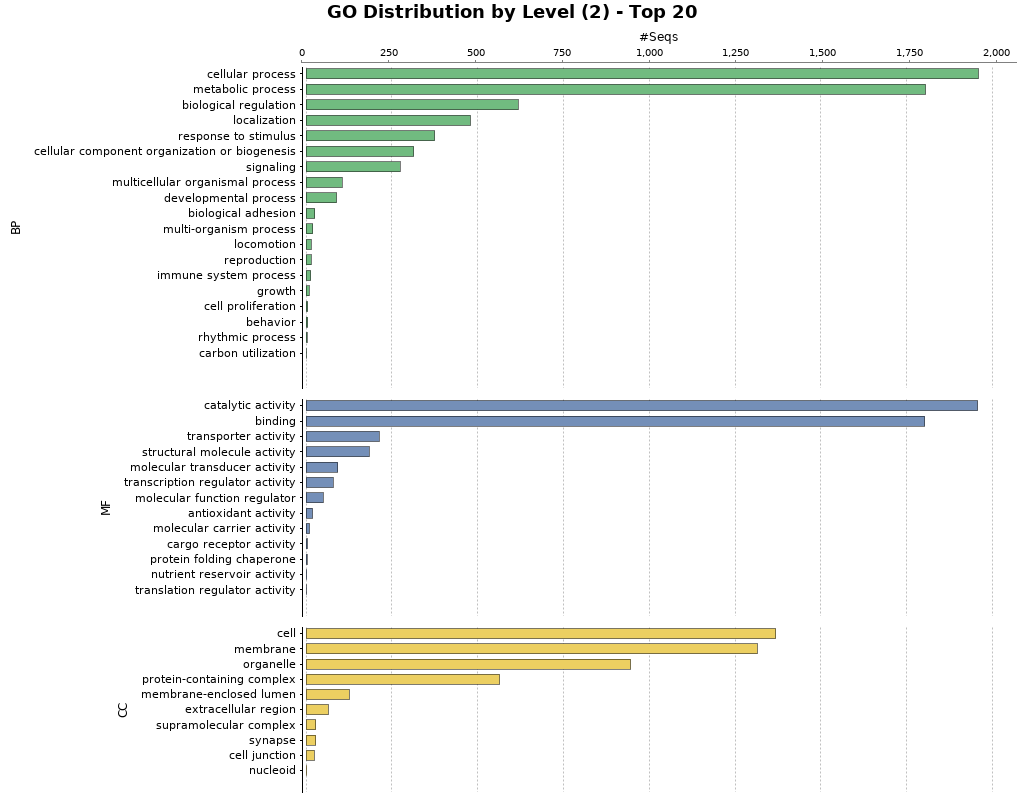

**Fig. S28. Blast2GO Functional Annotation for *Rhyacophila evoluta HR1.*** Barplot showing GO terms characterized by biological process, molecular function, and cellular component. Barplots are grouped by biological process (BP), molecular function (MF), and cellular component (CC).

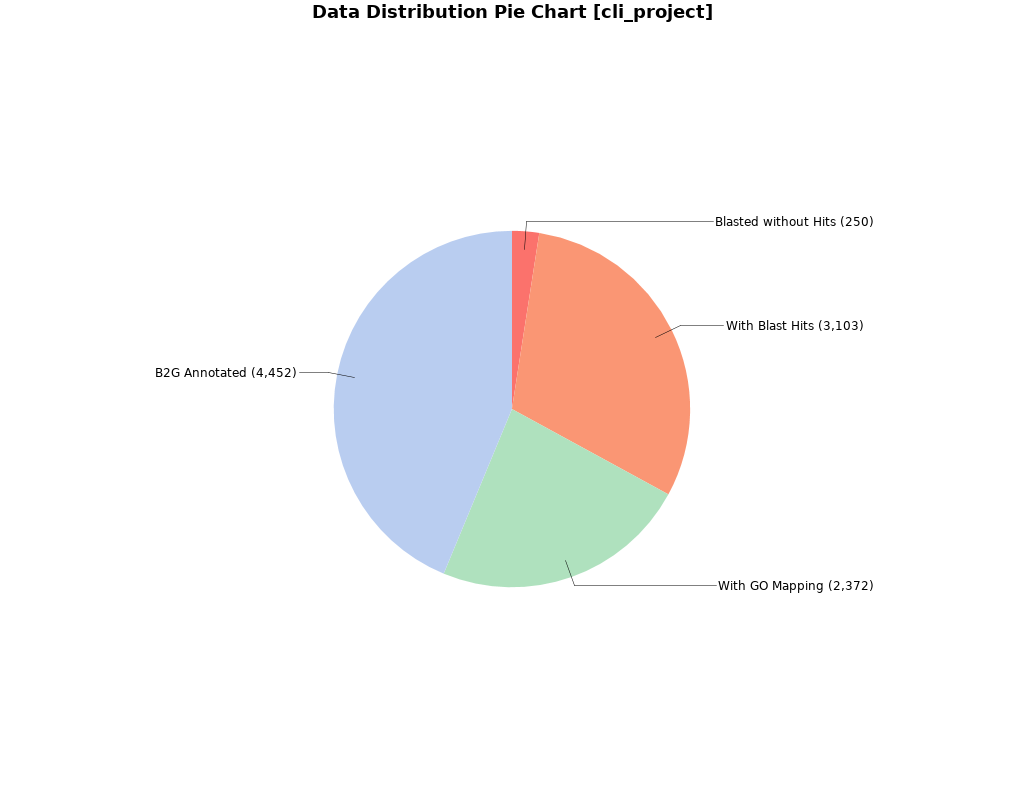

**Fig. S29. Blast2GO Annotation Results of *Rhyacophila evoluta RSS1*.** Pie charts showing the percentage of proteins with functional Blast2GO annotations, verified by BLAST and mapped to GO terms.

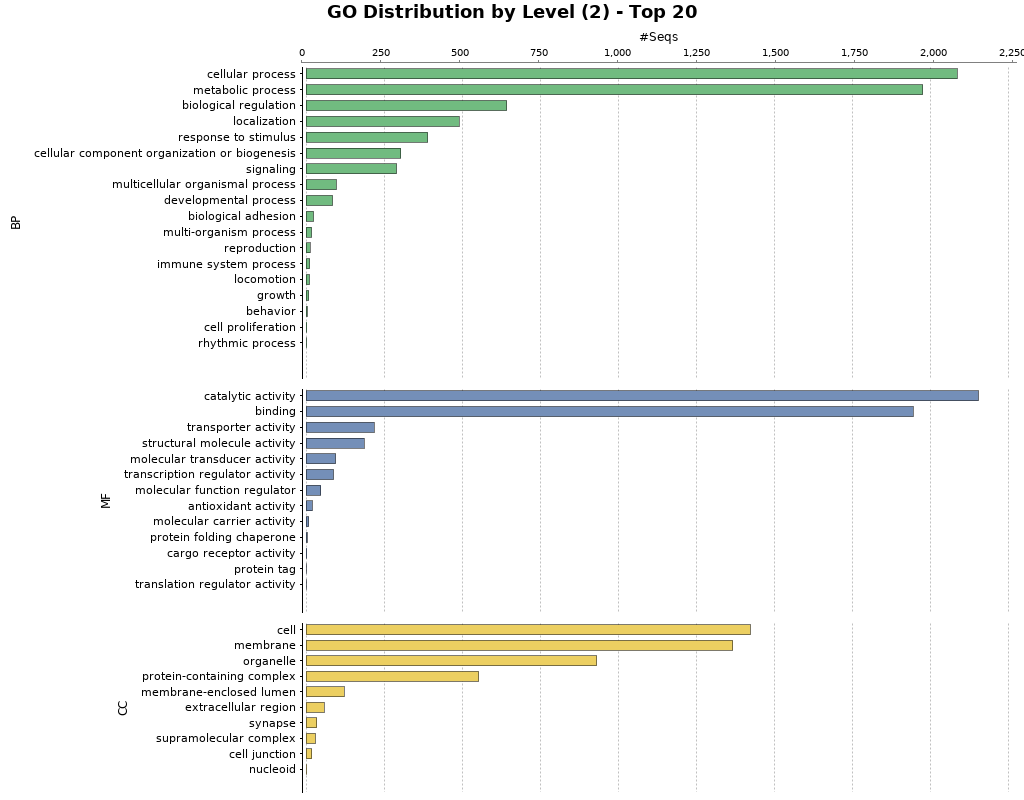

**Fig. S30. Blast2GO Functional Annotation for *Rhyacophila evoluta RSS1.*** Barplot showing GO terms characterized by biological process, molecular function, and cellular component.

**Supplementary note 5: Contamination filtering**

The final genome assemblies were screened and filtered for potential contaminations with taxon-annotated GC-coverage (TAGC) plots using BlobTools v1.0 [18]. For this purpose, all preprocessed Illumina reads were mapped against the final genome assemblies using BWA-MEM v0.7.17-r1188 [19]. Taxonomic assignment for BlobTools was done with blastn using the following parameters:

-*task megablast*, *max_target_seqs 1*, and -*max_hsps 1*. We only classified contigs with taxonomic assignment to phyla other than Arthropoda as contaminated when these also showed different GC content and coverage. The taxonomic assignment may be not as expected because a) some contigs are just too small to be recognized correctly and b) there might be conflict in the assignment and therefore blobtools just declares it as no-hit (see “taxrule” in bloptools manual). A valid reason for not excluding any contigs of assemblies shown in figures S31, S36, S38, S44, S45, S46, S47 is that in every "fraction" of the blobs there are contigs assigned to arthopods. However, contaminations were detected in some assemblies (S32-S34, S35, S37, S39, S40-S43). To filter out contigs based on the Blobplots, a table containing the information on coverage, GC content and taxonomic assignment was produced with the option *view* of BlobTools. We filtered contigs with potential contaminations from the assemblies using (xargs samtools faidx *>contaminated assembly> <contaminationfree_contigs.txt> > <filtered_assembly>*) and annotations (using *python filter_gffs.py <contaminated_contigs.txt> <contaminated.gff>*) after criteria outlined in legends of Figs. S31 to S47. Moreover, during uploading the genomes to NCBI, the NCBI genome assembly processing pipeline served as a second quality control. It showed that our contamination filtering was not conservative enough in few cases since it detected additional contaminations in some assemblies. These were further filtered out manually or by NCBI directly as follows: . When the contamination was not the whole contig but only parts of it, the contingency was broken where the contamination sequences were. We filtered out the contigs which contained contaminations from the MAKER annotations as described above, re-annotated the contamination filtered contigs and combined the results with the previous MAKER runs.

*Micrasema longulum ML1:*

ctg163846, ctg19699, ctg248737, ctg325643, ctg333503, ctg53726, ctg54533, ctg10643, ctg125325, ctg14195, ctg180425, ctg21916, ctg340516, ctg34404, ctg60979

*Micropterna sequax AB08:*

scf7180000362978: 811..1199, scf7180000455877: 11989..12080

*Philopotamus ludificatus Ph2:* contig25005

*Lepidostoma basale LB1:* scf1713:1059388..1059813

*Halesus radiates L2*: scf3403:246974..252200

*Rhyacophila brunneae Rhy_bru*:

scf1041: 163140..163728

scf1110:551504..551909

scf1128:33014..33597

scf1184:1536502..1536986

scf1270:1559810..1559871,1560071..1560361

scf1305:1032669..1033098,1034246..1034303

scf138:267360..267733

scf1417:192623..193022

scf1512:1368136..1368538

scf1588:293425..293830

scf1905:2898967..2899621

scf2084:985175..985580

scf418:215063..215545

scf541:247440..247517,251355..251805

scf651:112771..113167

scf713:77993..78081,78612..78966

scf79:2533756..2533988,2534216..2534291

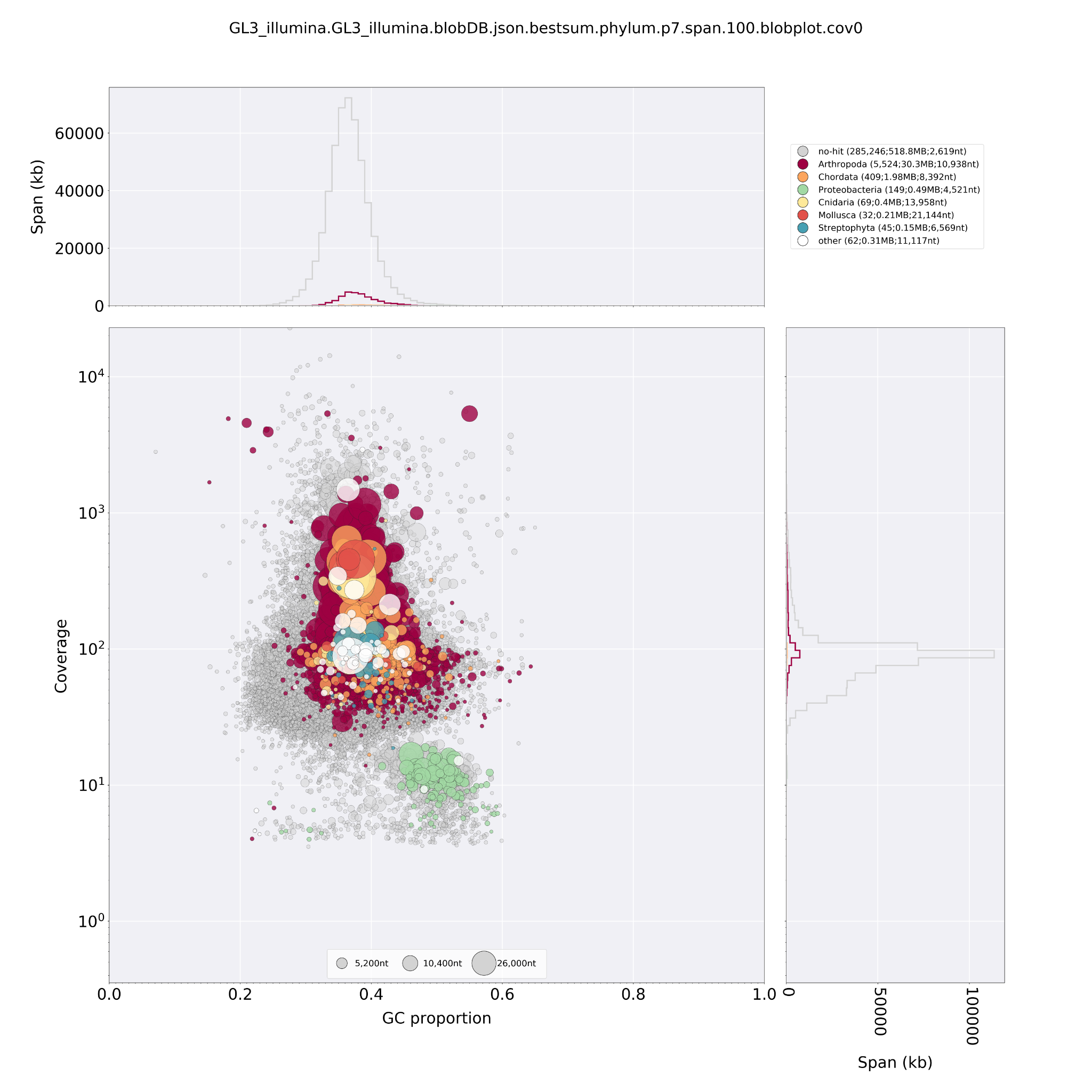

**Fig. S31. Taxon-annotated GC-coverage (TAGC) plots of *Agapetus fuscipens GL3* genome assembly*.*** Circles indicate contigs and the color indicates the best match to taxon annotation. The upper and right hand panel show the total span of contigs (kb) given GC proportion.

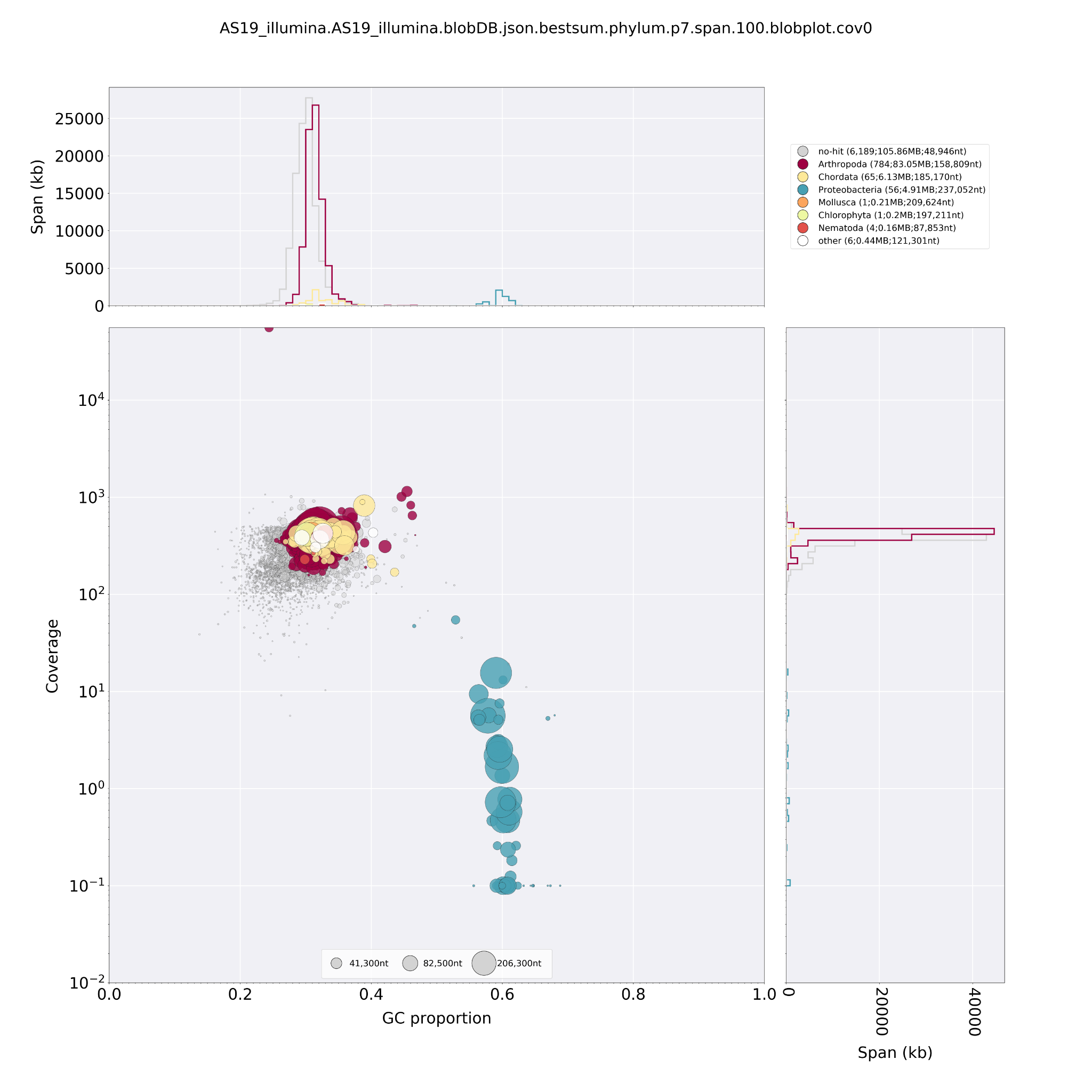

**Fig. S32. Taxon-annotated GC-coverage (TAGC) plots of *Agraylea sexmaculata AS19* genome assembly*.*** Circles indicate contigs and the color indicates the best match to taxon annotation. The upper and right hand panel show the total span of contigs (kb) given GC proportion. We removed all contigs assigned to Proteobacteria.

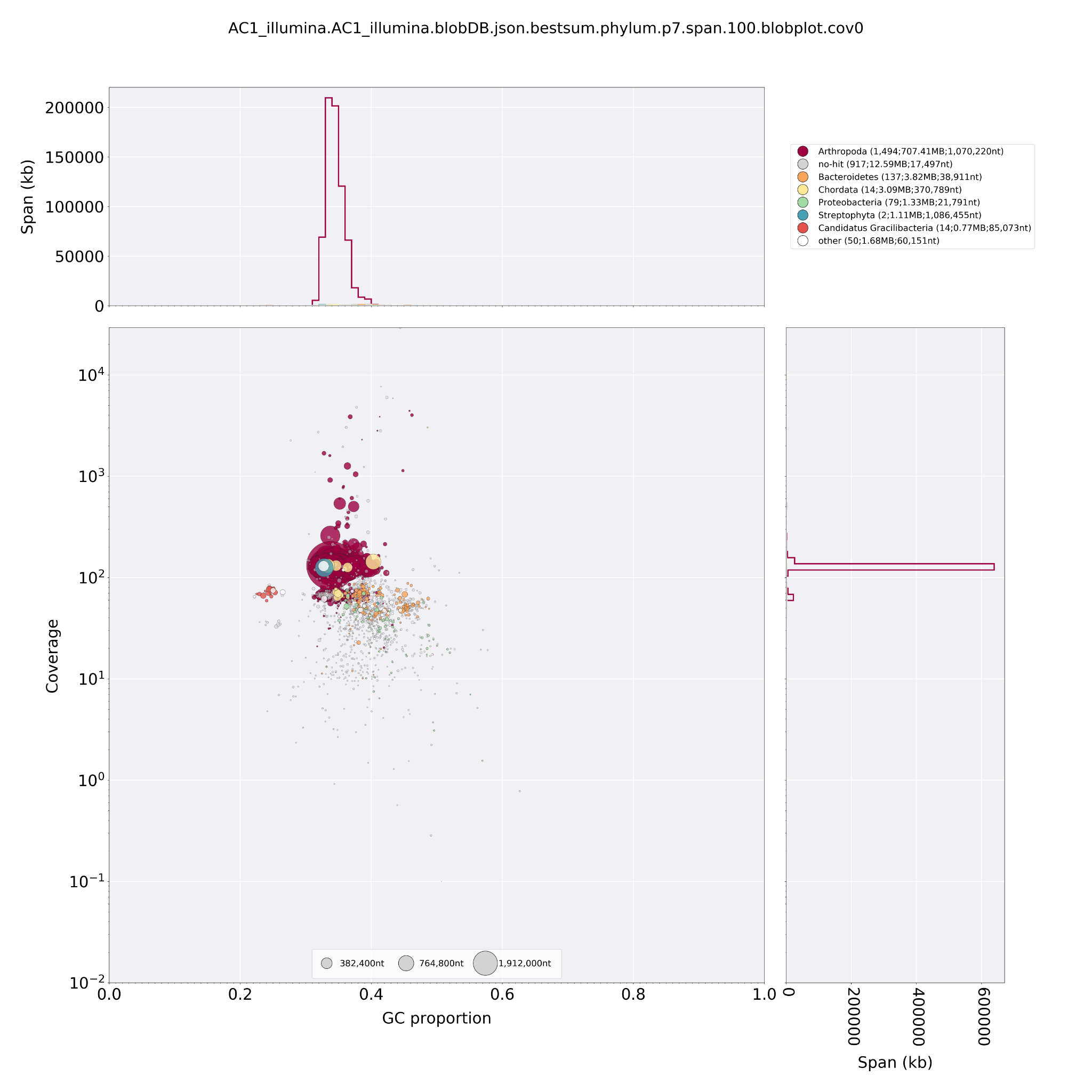

**Fig. S32. Taxon-annotated GC-coverage (TAGC) plots of *Drusus annulatus AC1* genome assembly*.*** Circles indicate contigs and the color indicates the best match to taxon annotation. The upper and right hand panel show the total span of contigs (kb) given GC proportion. We removed contigs with a coverage ≤ the lowest coverage in “Arthropod” contigs (20.363) and contigs which were out of the GC content range of “Arthropod” contigs (0.3042 - 0.4621).

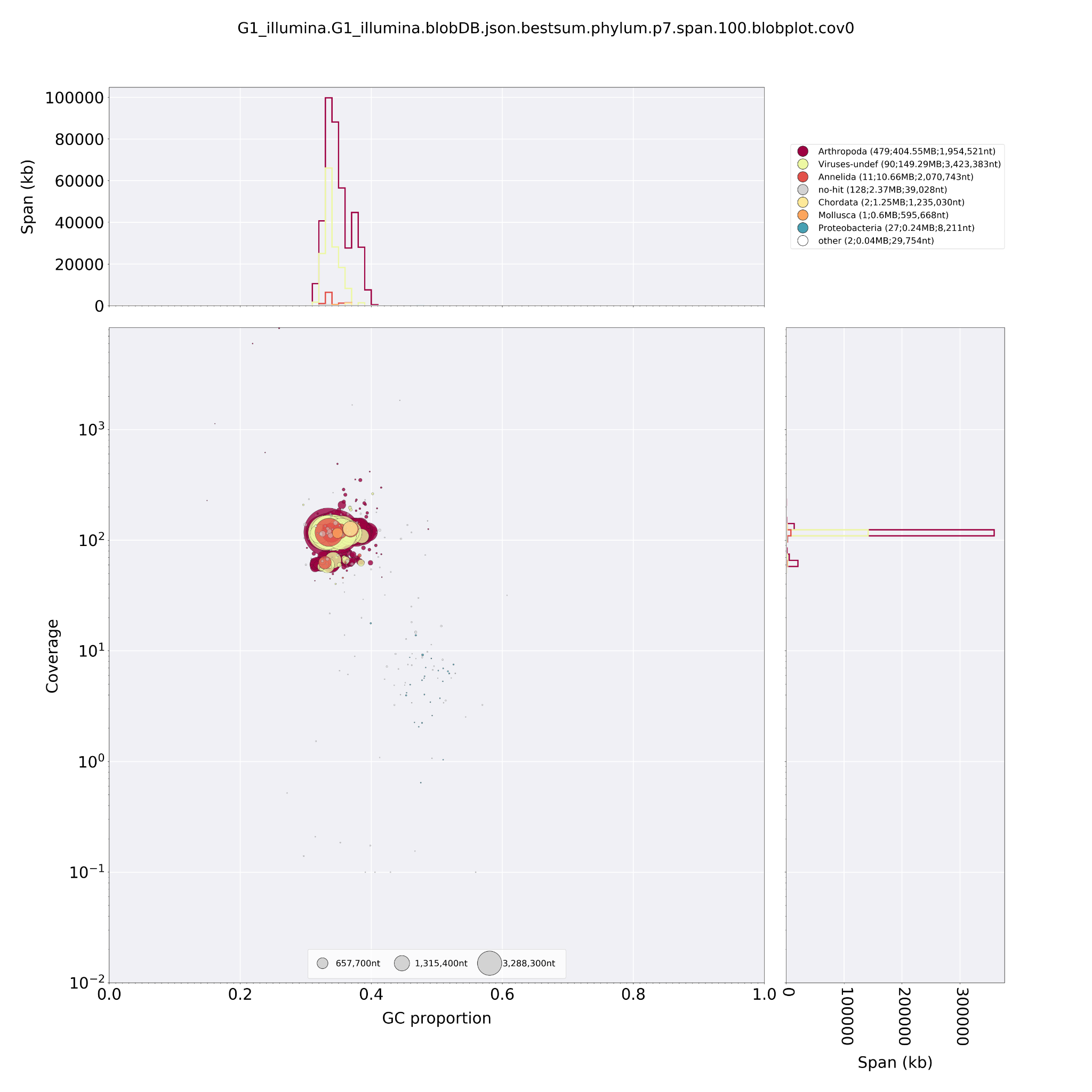

**Fig. S34. Taxon-annotated GC-coverage (TAGC) plots of *Glossosoma conforme G1****.* Circles indicate contigs and the color indicates the best match to taxon annotation. The upper and right hand panel show the total span of contigs (kb) given GC proportion. We removed contigs with coverage ≤ the lowest coverage in “Arthropod” contigs (43.031) and contigs with GC content ≥ highest GC content in “Arthropod” contigs (0.487).

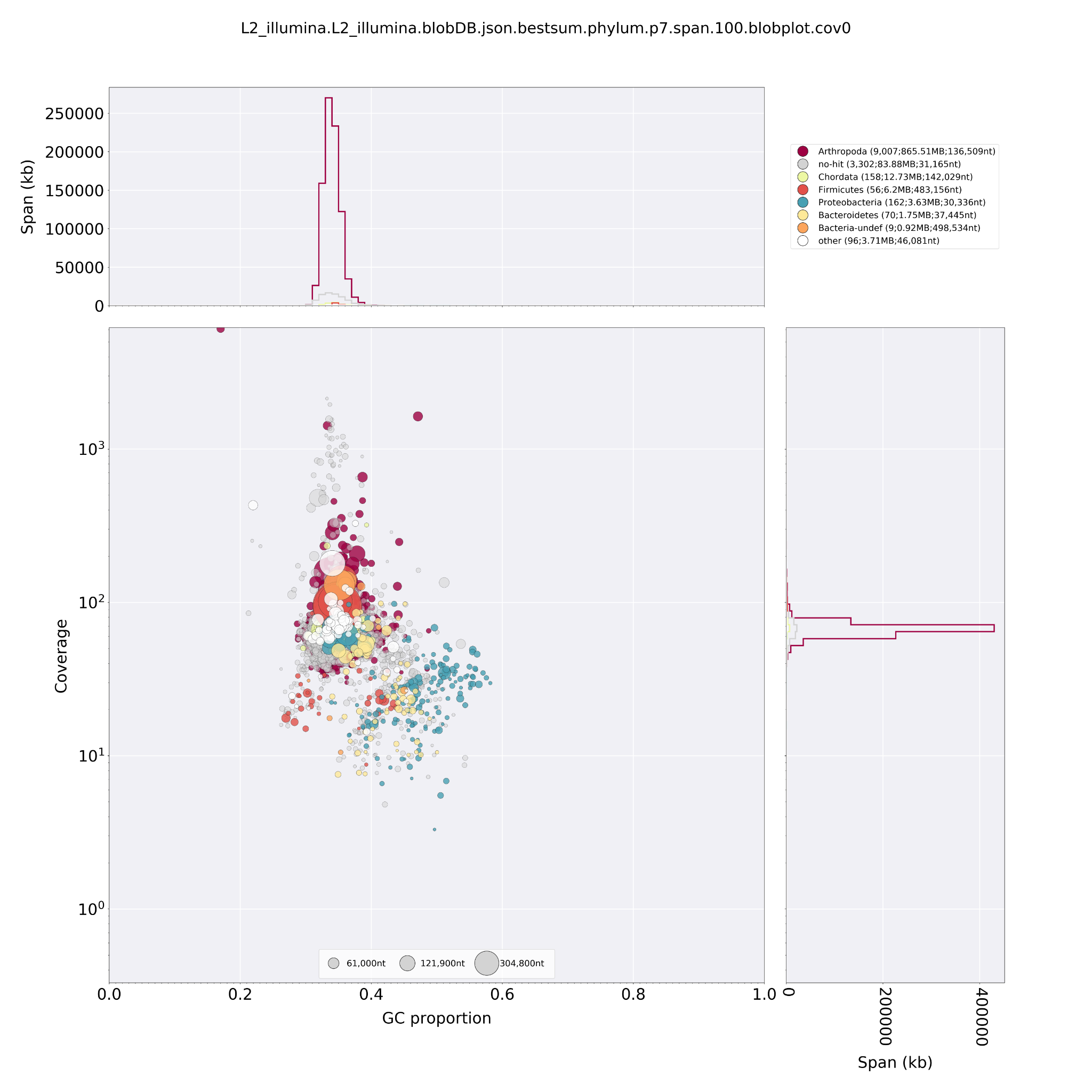

**Fig. S35. Taxon-annotated GC-coverage (TAGC) plots of *Halesus radiatus.*** Circles indicate contigs and the color indicates the best match to taxon annotation. The upper and right hand panel show the total span of contigs (kb) given GC proportion. We removed contigs with coverage ≤ the lowest coverage in “Arthropod” contigs (30.06) and contigs with GC content ≥ highest GC content in “Arthropod” contigs (0.4876).

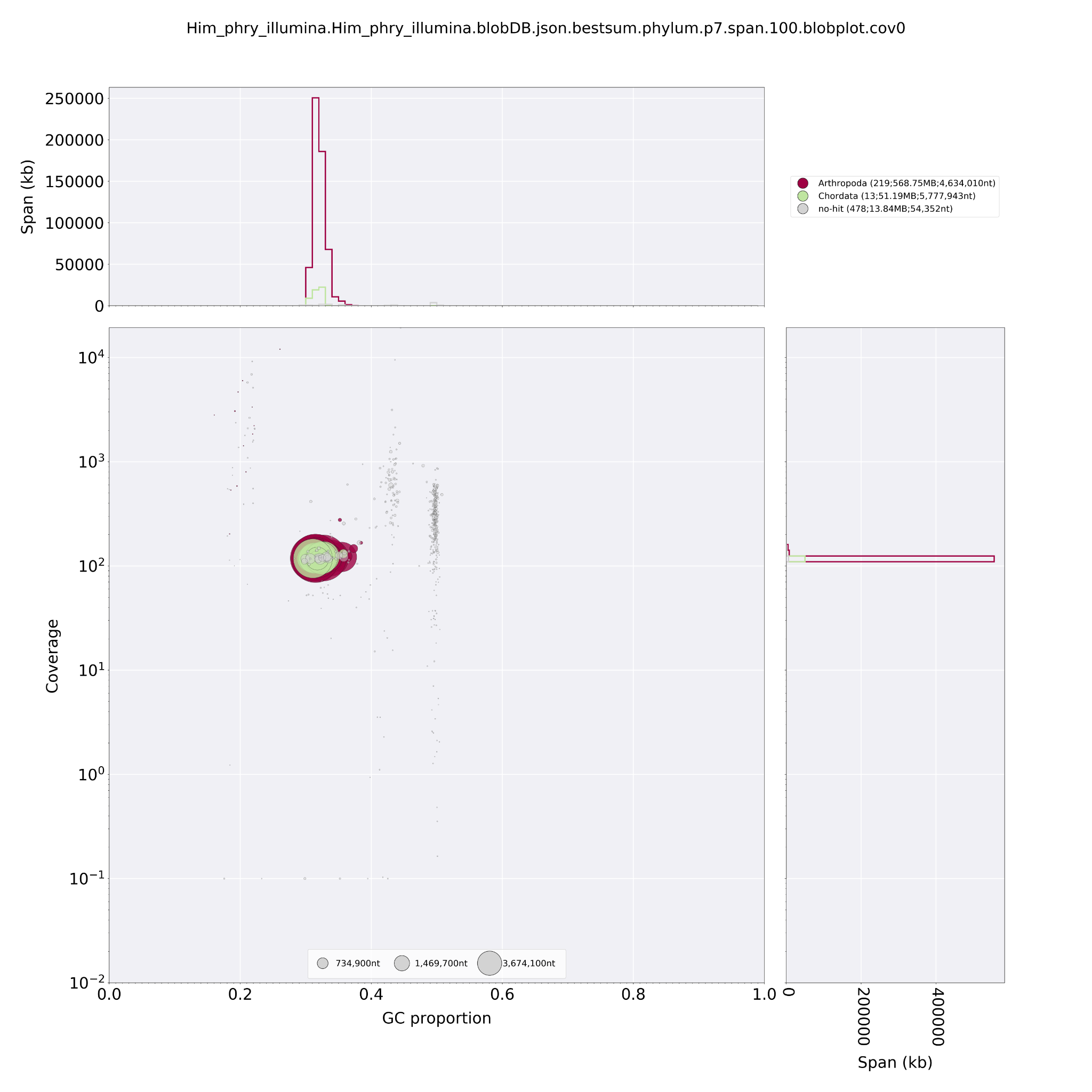

**Fig. S36. Taxon-annotated GC-coverage (TAGC) plots of *Himalopsyche phryganeae.*** Circles indicate contigs and the color indicates the best match to taxon annotation. The upper and right hand panel show the total span of contigs (kb) given GC proportion***.***

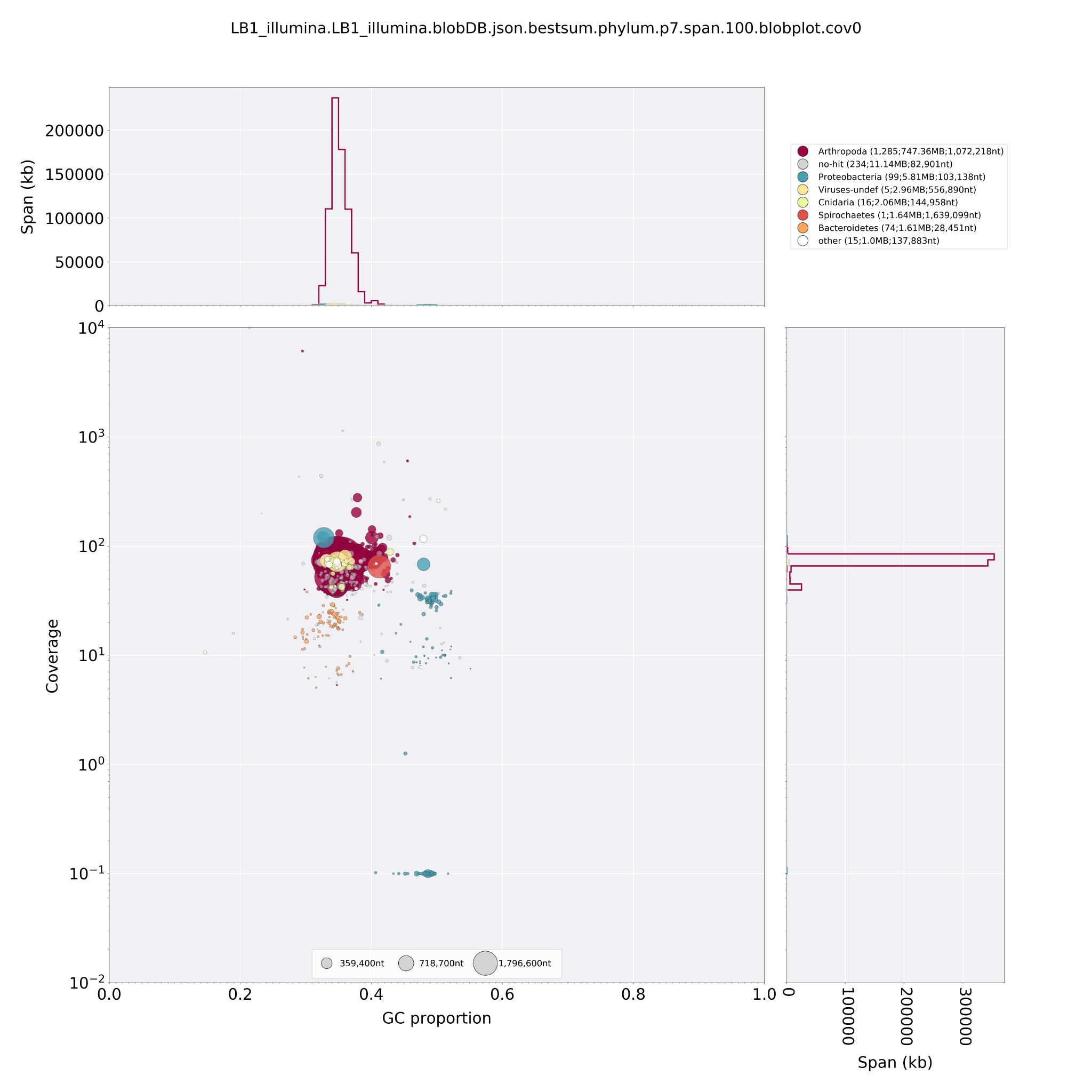

**Fig. S37. Taxon-annotated GC-coverage (TAGC) plots of *Lepidostoma basale.*** Circles indicate contigs and the color indicates the best match to taxon annotation. The upper and right hand panel show the total span of contigs (kb) given GC proportion***.*** We removed contigs with a coverage ≤ the lowest coverage in “Arthropod” contigs (5.336) and contigs which were out of the GC content range of “Arthropod” contigs (0.295 - 0.4657).

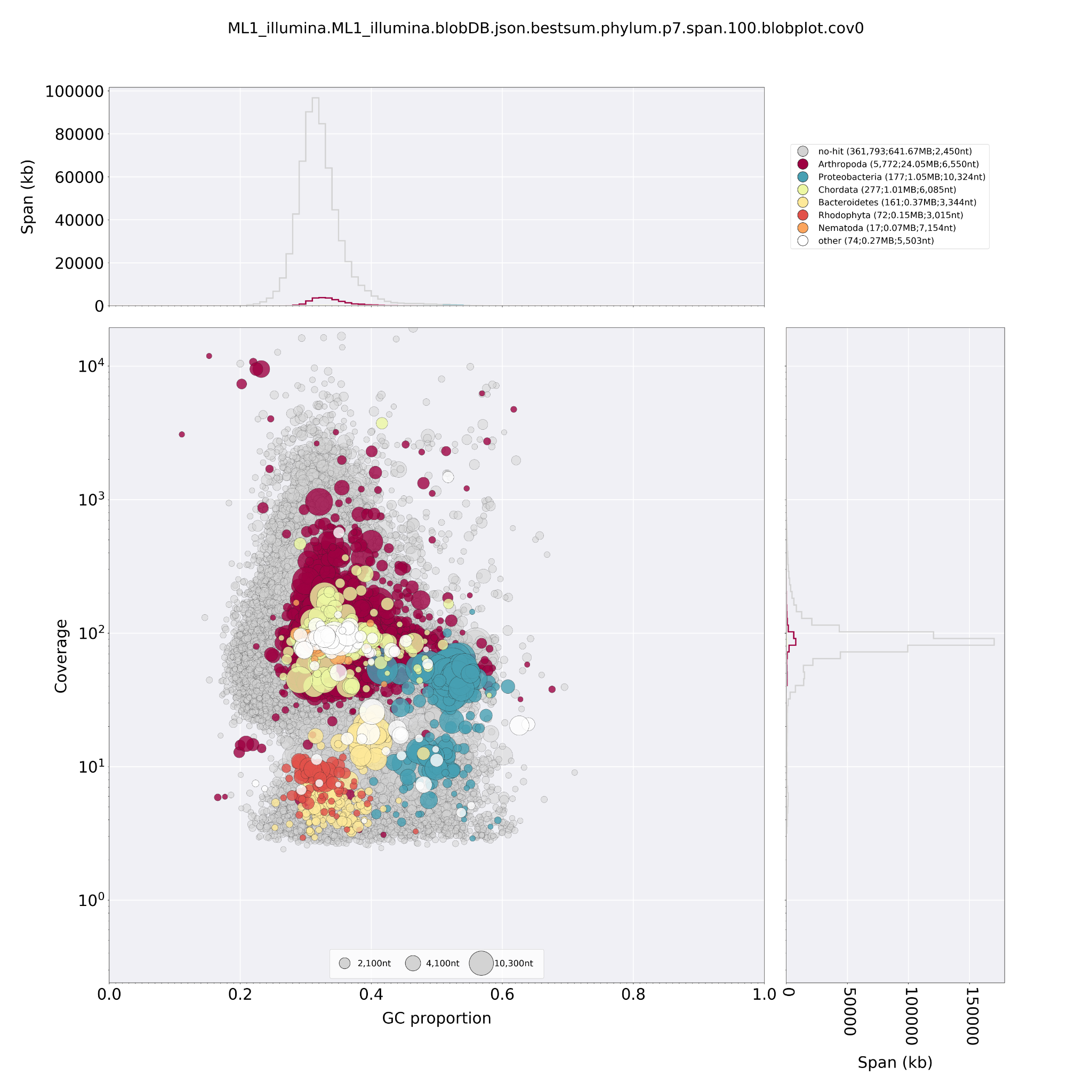

**Fig. S38. Taxon-annotated GC-coverage (TAGC) plots of *Micrasema longulum ML1.*** Circles indicate contigs and the color indicates the best match to taxon annotation. The upper and right hand panel show the total span of contigs (kb) given GC proportion***.***

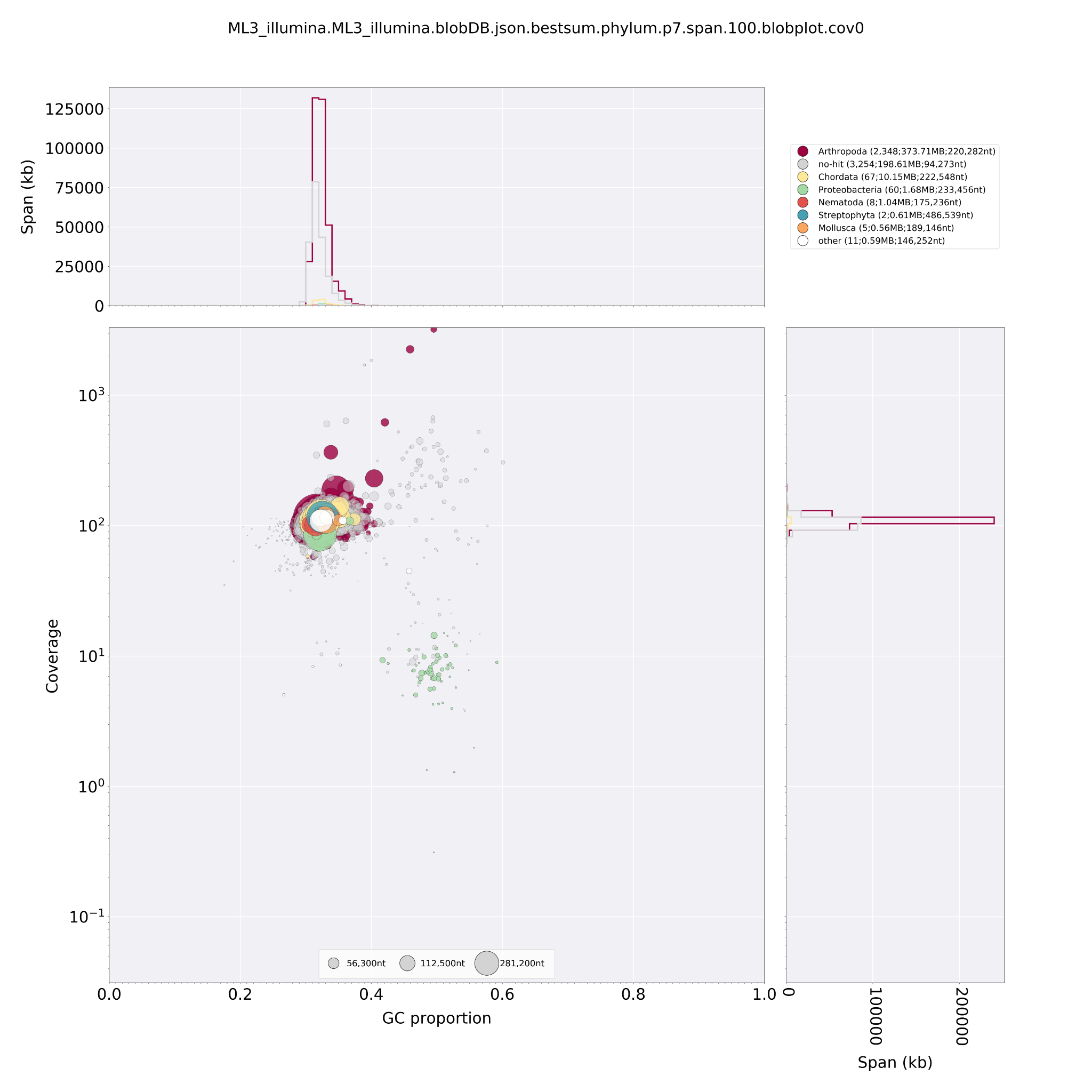

**Fig. S39.Taxon-annotated GC-coverage (TAGC) plots of *Micrasema longulum ML3.*** Circles indicate contigs and the color indicates the best match to taxon annotation. The upper and right hand panel show the total span of contigs (kb) given GC proportion***.*** We removed contigs with a coverage ≤ the lowest coverage in “Arthropod” contigs (56.642) and contigs which were out of the GC content range of “Arthropod” contigs (0.2962 - 0.4954).

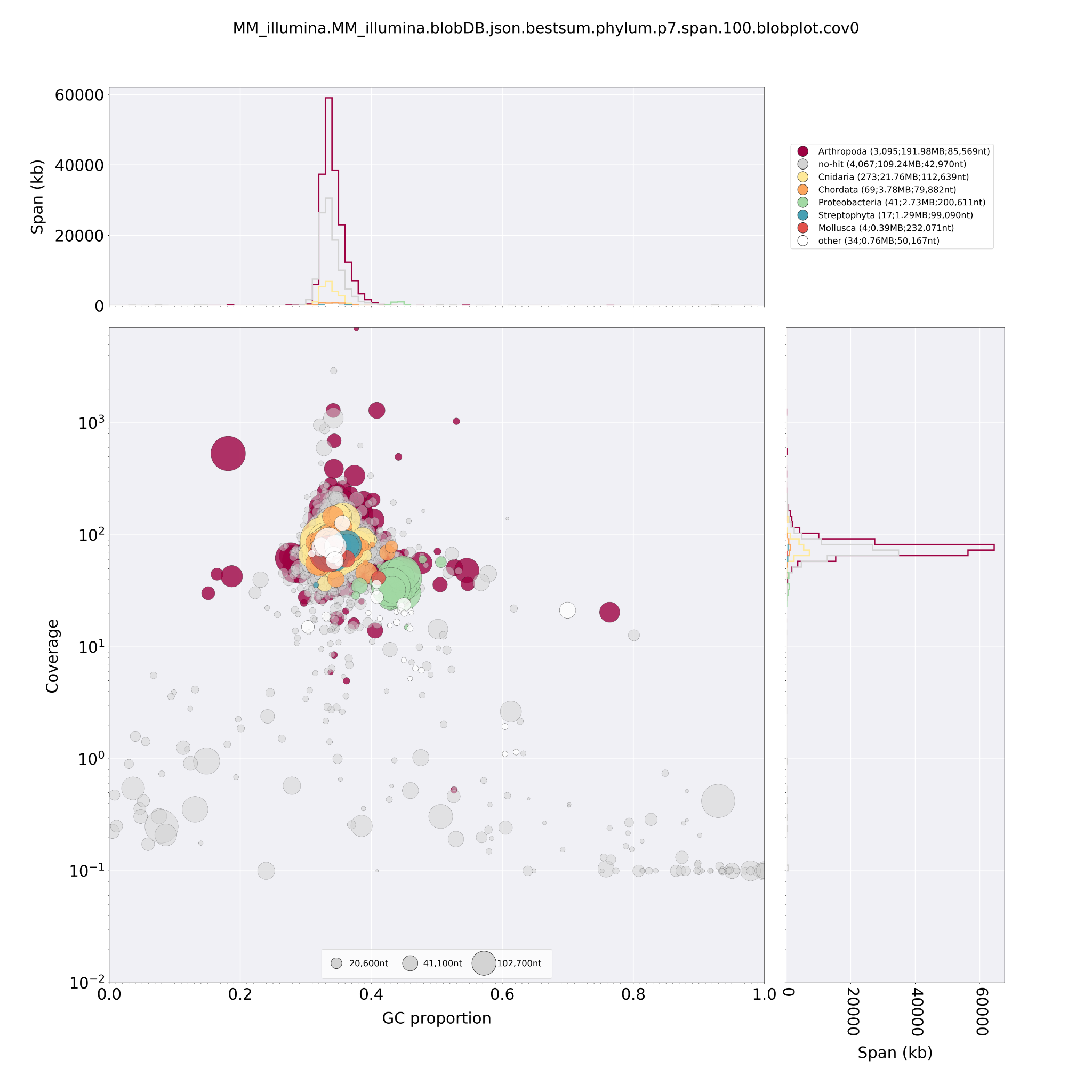

**Fig. S40. Taxon-annotated GC-coverage (TAGC) plots of *Micrasema minimum.*** Circles indicate contigs and the color indicates the best match to taxon annotation. The upper and right hand panel show the total span of contigs (kb) given GC proportion***.*** We removed contigs to Proteobacteria with a GC content of ≥ 0.4.

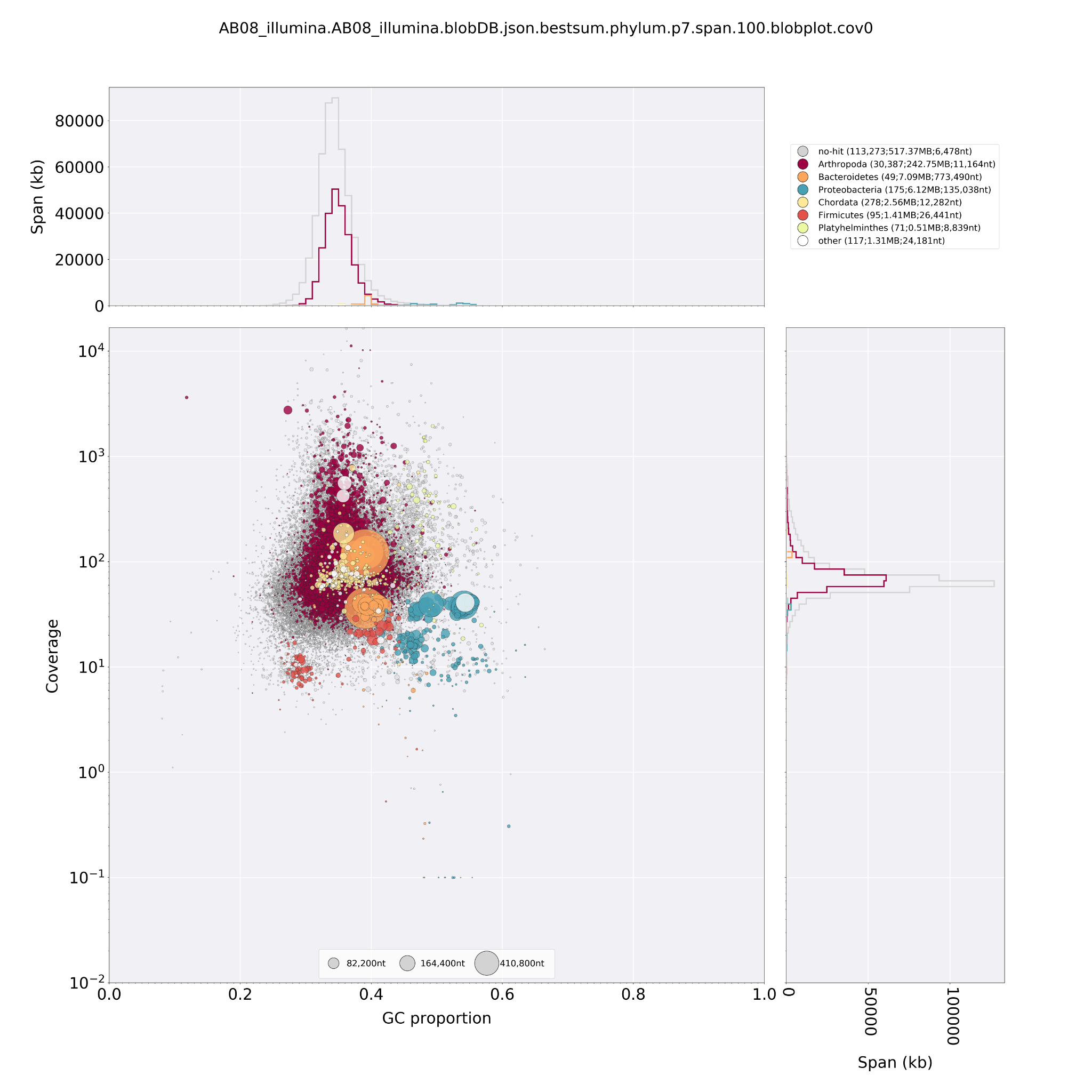

**Fig. S41. Taxon-annotated GC-coverage (TAGC) plots of *Micropterna sequax.*** Circles indicate contigs and the color indicates the best match to taxon annotation. The upper and right hand panel show the total span of contigs (kb) given GC proportion***.*** We removed contigs with a coverage ≤ the lowest coverage in “Arthropod” contigs (5.665) and contigs which were out of the GC content range of “Arthropod” contigs (0.1182 - 0.5602).

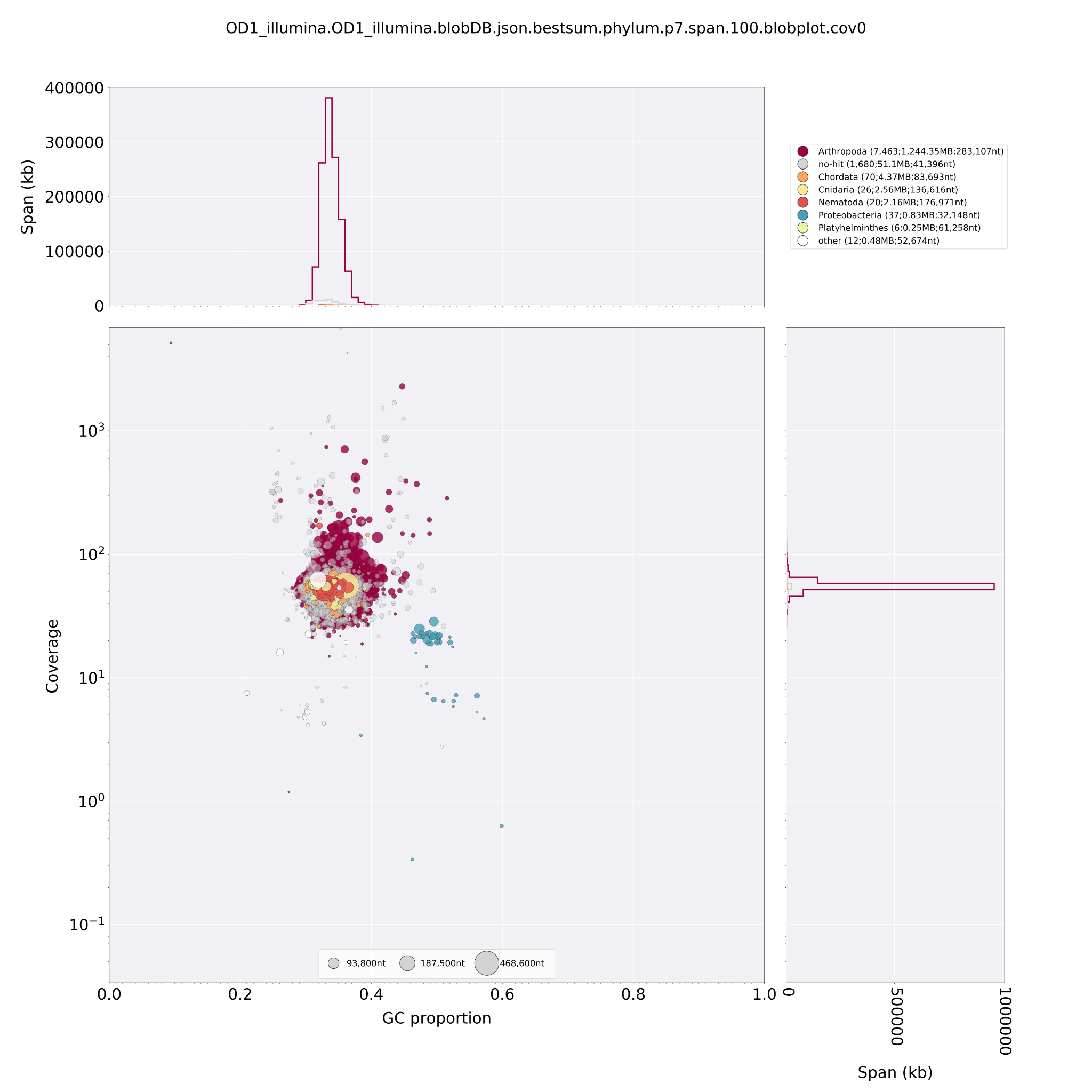

**Fig. S42. Taxon-annotated GC-coverage (TAGC) plots of *Odontocerum albicorne.*** Circles indicate contigs and the color indicates the best match to taxon annotation. The upper and right hand panel show the total span of contigs (kb) given GC proportion***.*** We removed contigs with coverage ≤ 1 and contigs with GC content ≥ highest GC content in “Arthropod” contigs (0.5157).

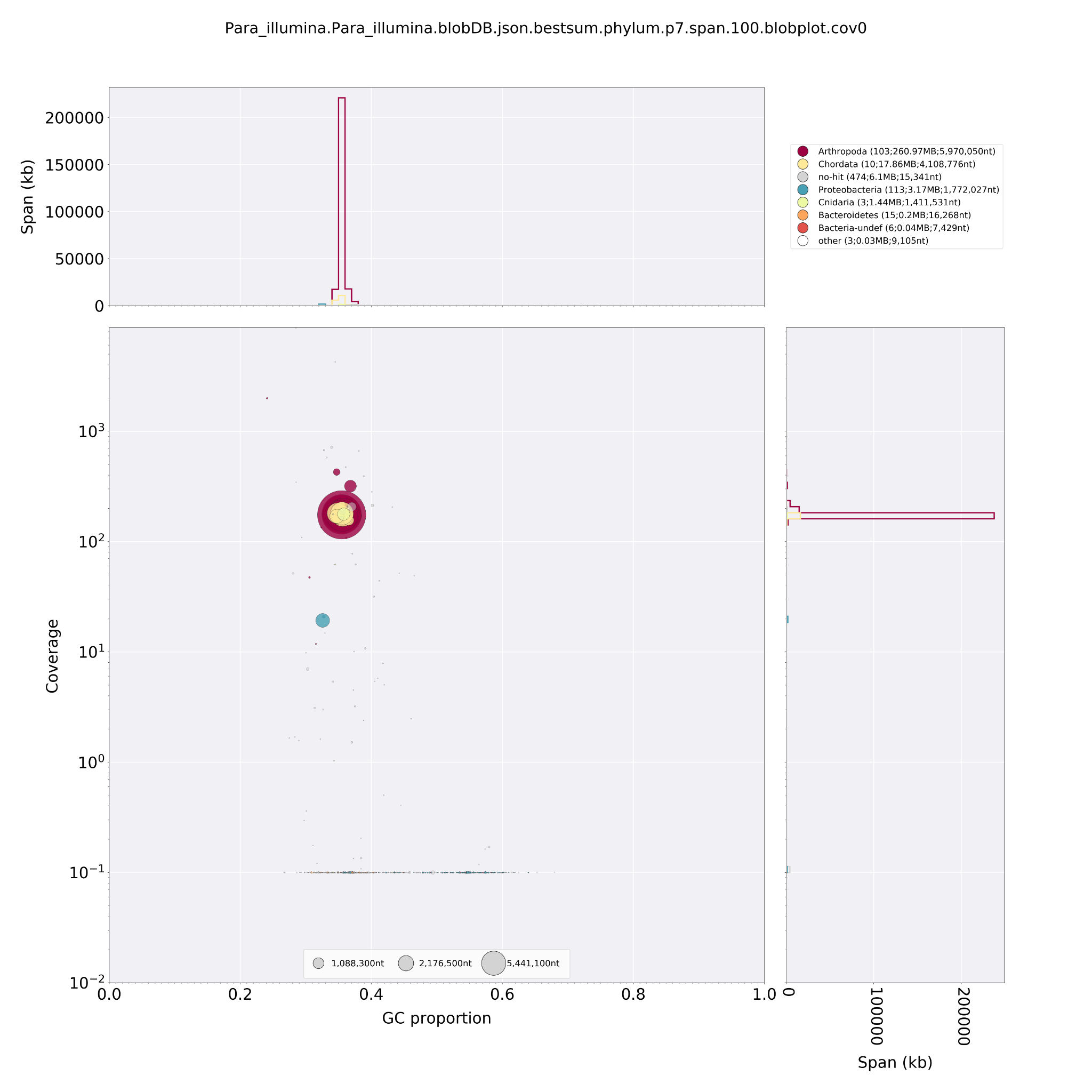

**Fig. S43. Taxon-annotated GC-coverage (TAGC) plots of *Parapsyche elsis.*** Circles indicate contigs and the color indicates the best match to taxon annotation. The upper and right hand panel show the total span of contigs (kb) given GC proportion***.*** We removed all contigs assigned to Proteobacteria and contigs with coverage ≤ 1.

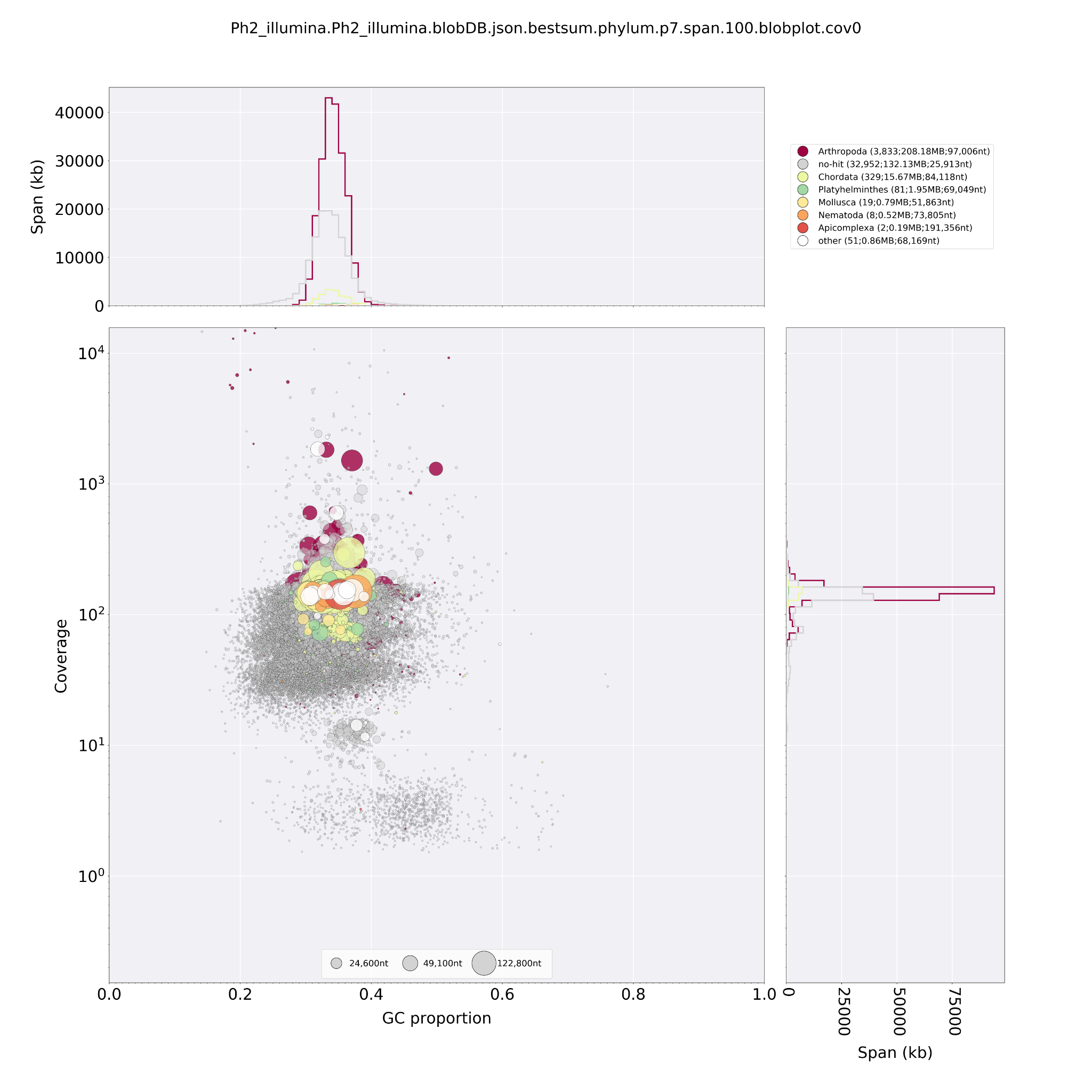

**Fig. S44.Taxon-annotated GC-coverage (TAGC) plots of *Philopotamus ludiferatus.*** Circles indicate contigs and the color indicates the best match to taxon annotation. The upper and right hand panel show the total span of contigs (kb) given GC proportion***.***

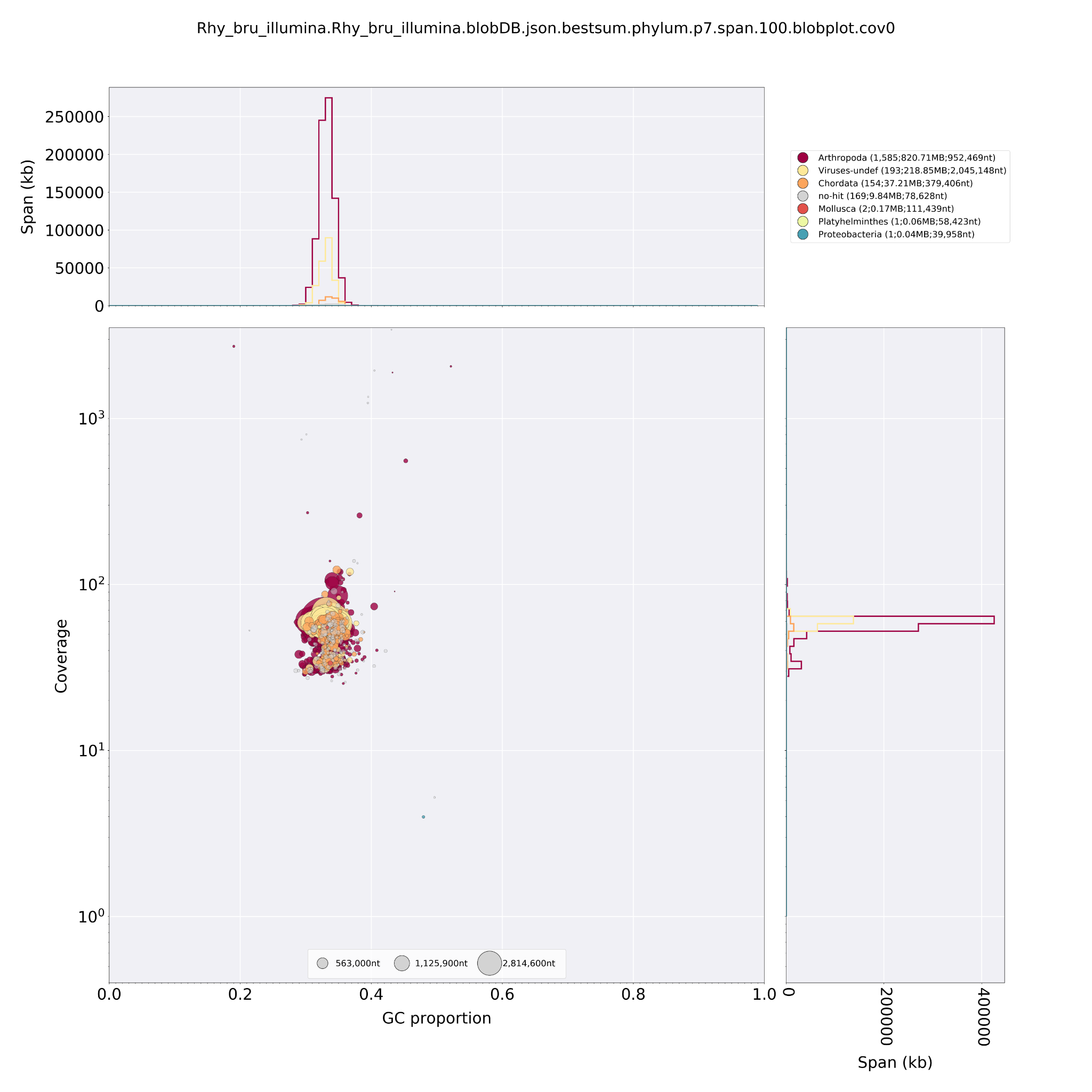

**Fig. S45. Taxon-annotated GC-coverage (TAGC) plots of *Rhyacophila brunneae.*** Circles indicate contigs and the color indicates the best match to taxon annotation. The upper and right hand panel show the total span of contigs (kb) given GC proportion***.***

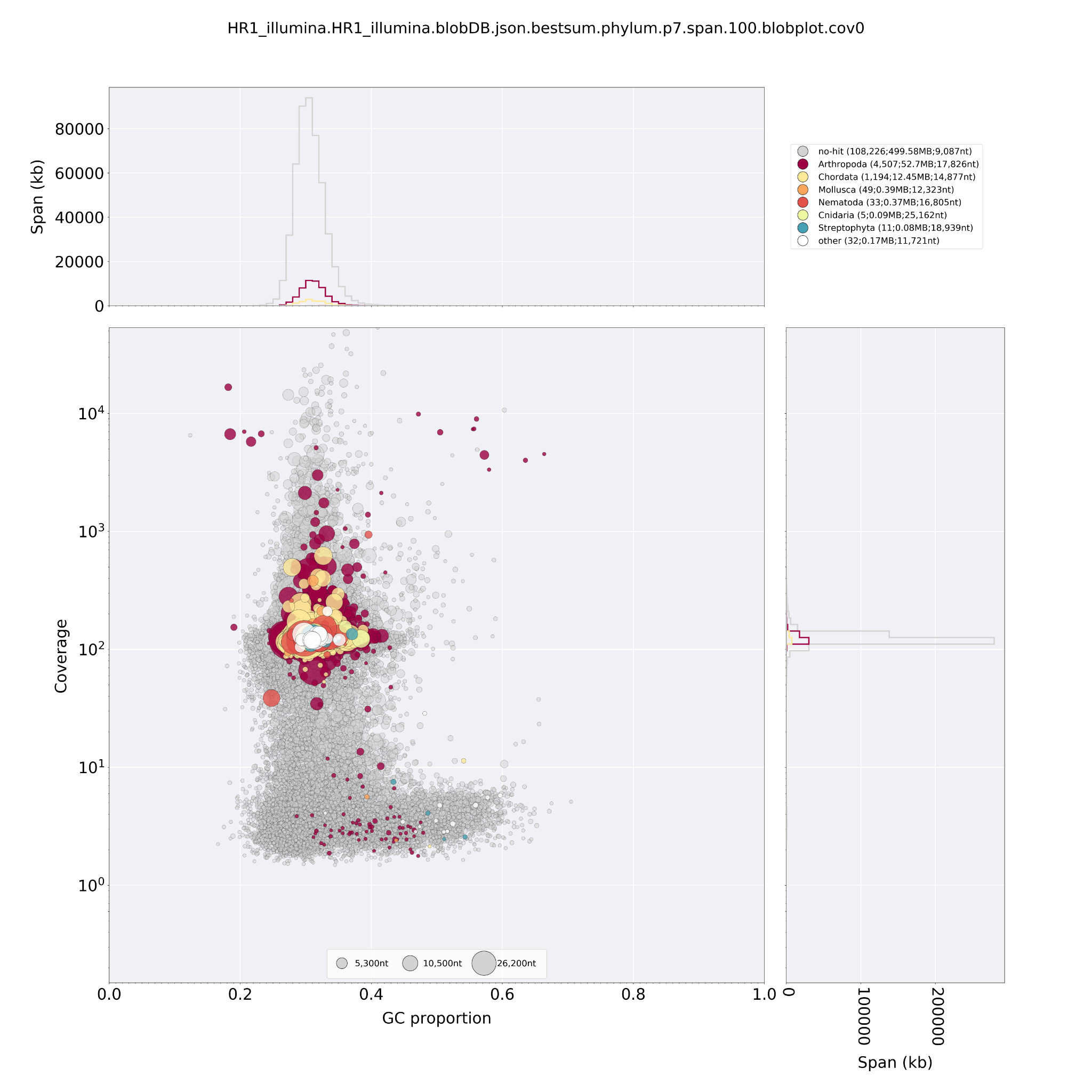

**Fig. S46. Taxon-annotated GC-coverage (TAGC) plots of *Rhyacophila evoluta HR1****.* Circles indicate contigs and the color indicates the best match to taxon annotation. The upper and right hand panel show the total span of contigs (kb) given GC proportion***.***

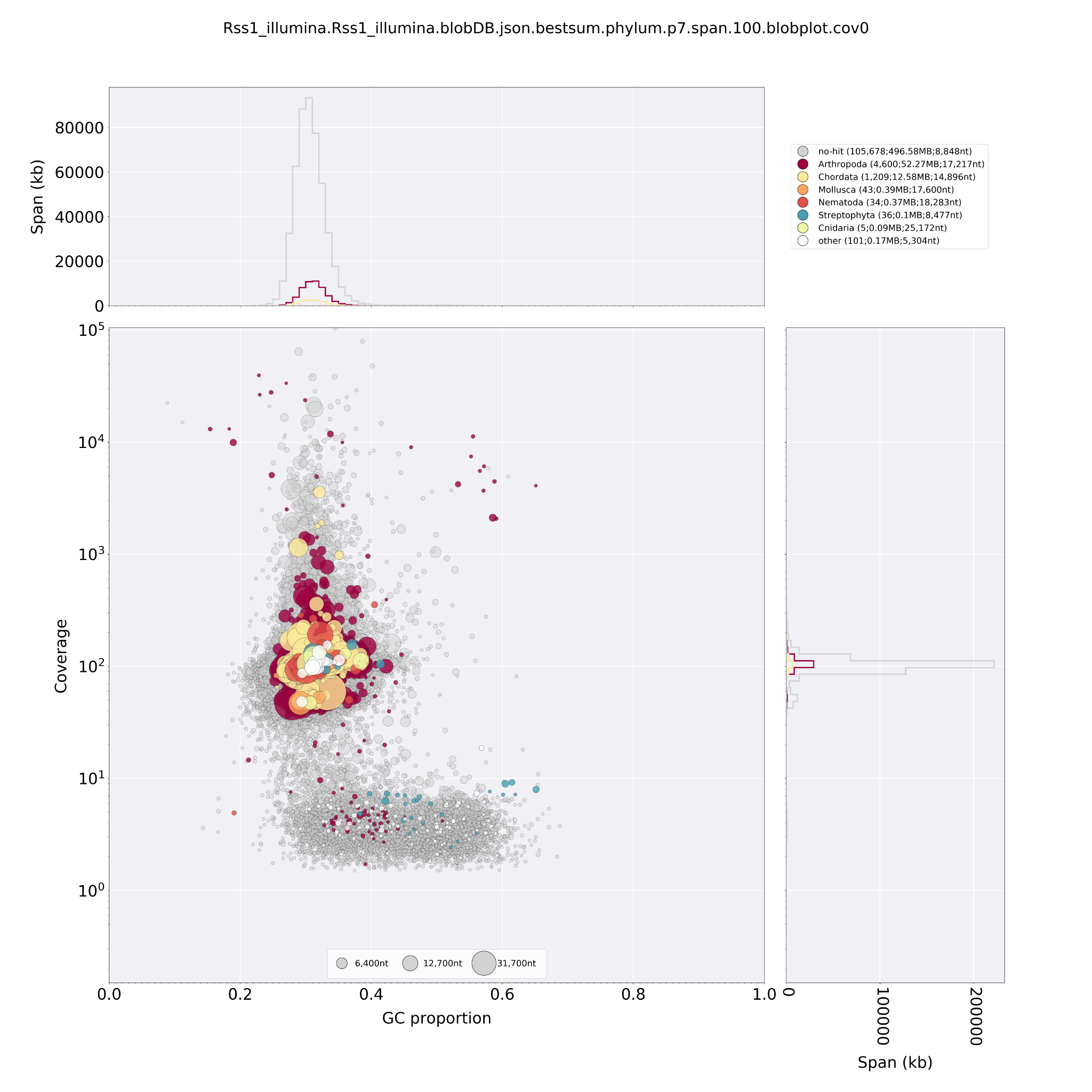

**Fig. S47. Taxon-annotated GC-coverage (TAGC) plots of *Rhyacophila evoluta RSS1****.* Circles indicate contigs and the color indicates the best match to taxon annotation. The upper and right-hand panel show the total span of contigs (kb) given GC proportion***.***

Supplementary note 6: Caddisfly silk usage

Trichoptera use silk produced in their salivary glands to build diverse underwater structures. While Annulipalpia is represented by fixed retreat- and net-making species, Integripalpia consists of ‘tube case’- and ’cocoon’-making species. Larvae of most families in suborder Integripalpia create portable, tubular cases made from diverse materials encountered in their habitats, such as small stones or plant material or made purely from silk. These larvae then pupate in their final case, by encapsulating themselves within the portable structure and fixing this to the substrate. Basal integripalpian families which show diverse case-making behaviors (‘free-living’, ‘tortoise case’-making, ‘purse case’-making), are referred to as cocoon-makers because they pupate in a silken pupal cocoon with an internal osmotic environment [20]. Previously, cocoon‐making caddisflies were recognized as their own suborder ‘Spicipalpia’ [21]. However, recent studies [22] rejected ‘Spicipalpia’ as a third suborder and defined Integripalpia to include both the tube‐case (Phryganides) and cocoon‐makers (basal Integripapia) as previously suggested by [23]. Our results support this classification (Fig. 1, main text, supplementary Fig. 48), though more data is needed to assess the phylogenetic position of *Agraylea sexmaculata* (Hydroptilidae, see polytomy in supplementary Fig. S48) as the only ‘purse-case’ maker in this study.

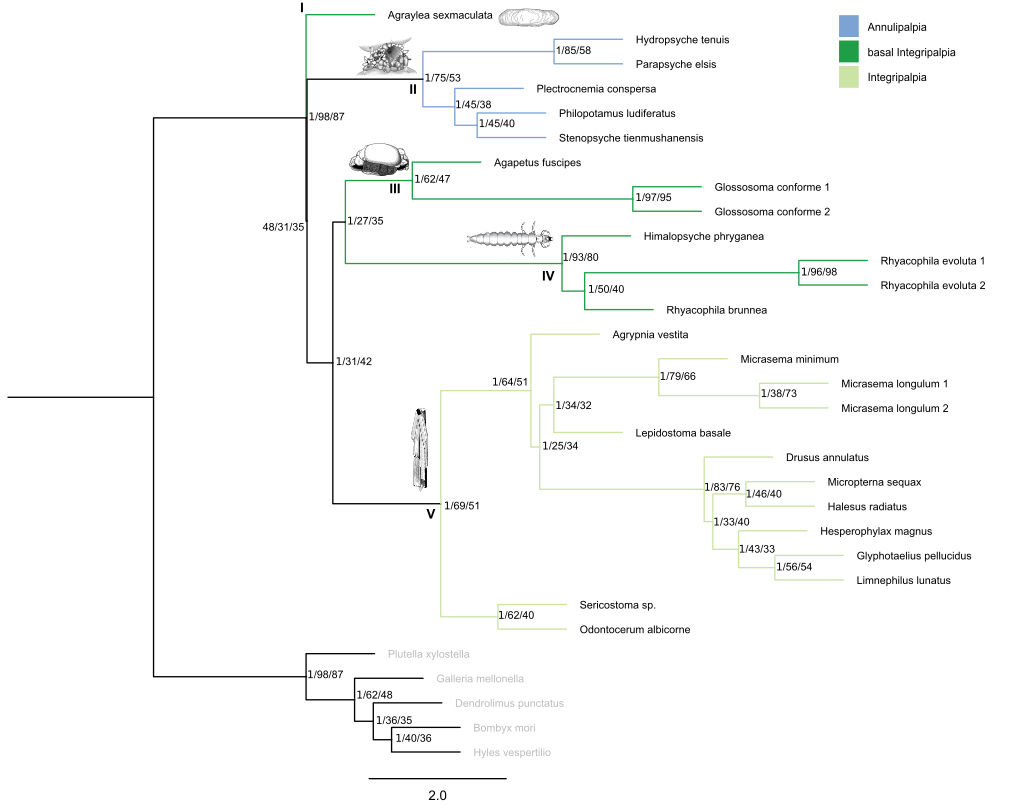
**Fig. S48.** Phylogenetic relationships derived from ASTRAL-III analyses using single BUSCO genes. Support values, site concordance and gene concordance factors are given for each node. Trichoptera are divided into two suborders: Annulipalpia (II: blue) which consists of ‘fixed retreat- and net-building’ species and Intergripalpia (I, III, IV, V: green) which includes ‘cocoon-making’ (basal Integripalpia, clades I, III, IV, dark green) and ‘case-building’ (V: light green) caddisflies. ‘Cocoon-makers’ are divided into ‘purse case’- (I), ‘tortoise case-making’ (III) and ‘free-living’ (IV) families. Illustrations represent an example of silk use in the respective clade. Taxa in grey (Lepidoptera) were used as outgroups.

**Supplementary note 7: Genome size estimations and genome profiling**

*k-mer distribution-based method*

We conducted genome profiling (estimation of major genome characteristics such as size, heterozygosity, and repetitiveness) on the trimmed, contamination filtered short-read sequence data with GenomeScope 2.0 [24]. Before running GenomeScope 2.0, we counted *k-mers* with JELLYFISH v2.2.10 [25]. using jellyfish count -C -s 25556999998 -F 3 and a *k-mer* length of 21 (-m 21) as recommended for most genomes by the authors of GenomeScope2. A histogram of *k-mer* frequencies was produced with jellyfish histo. GenomeScope 2.0 was run with the exported *k-mer* count histogram within the online web tool (http://qb.cshl.edu/genomescope/genomescope2.0/) using the following parameters: Kmer length = 21, Read length = 150, max *k-mer* coverage = 10000. For some species we ran the command-line version of Genomescope2 with the option -l to set the location of the first peak after correspondence with the Genomescope2 developer. For GenomeScope profiles see Data S1: 4-Genomescope 2 Results. For three specimens the online interface version of Genomescope2 had difficulties with finding the first peak with the default settings. After correspondence with the Genomescope2 developer, we ran the command-line version of Genomescope2 with the optional parameter -l lambda to set the initial guess for the average *k-mer* coverage of the sequencing based on the results of the Genomescope2 profile from the online version.

*Backmapping based approaches*

We mapped all trimmed and contamination filtered Illumina reads against the final genome assemblies using BWA-MEM v0.7.17-r1188 with the options -a and -c 10000. We used view -l -b and sort -l 9 functions of SAMtools v1.9 to print alignments to standard output in BAM format and sort them by leftmost coordinates. We conducted genome size estimations using backmap.pl v0.3 [26] with the options -nq and the sorted .bam files resulting from the re-mapping step. The script uses BEDTools v2.27.1 [27] to generate a coverage histogram and R v3.5.1 (R Core Team, 2017; https://www.R-project.org/) to plot the coverage distribution. Assuming even sequencing coverage throughout the genome, backmap.pl estimates the genome size by dividing the number of total nucleotides which were mapped to the assembly by the maximum of the per-position coverage frequency distribution. For coverage distribution per position see Supplementary Figs. 49-72. Genome size estimates are given in Data S1: 5-Backmap.pl Results.

*Flow cytometry*

We estimated genome sizes (2C-values,[28]) for 29 of Trichoptera species (19 genera) by flow cytometry (FCM) using the Partec CyFlow Space (Partec, Münster, Germany) equipped with a green solid-state laser (Partec, 532 nm, 30 mW). For sample preparation, we followed the two-step Otto protocol [29], with an internal standard *Lycopersicon esculentum* cv. Stupické polní tyčkové rané (2C = 1.96 pg; [30]). We mixed the whole or part of the caddisfly body with ca. 1 cm^2^ leaf of an internal reference standard and homogenized it with a razor blade in a Petri dish containing 1 ml of ice-cold Otto I buffer (0.1 M citric acid, 0.5% Tween 20; [29]). We filtered the suspension through a 42-μm nylon mesh and incubated it for approximately 15 min at room temperature. The staining solution consisted of 1 ml of Otto II buffer (0.4 M Na_2_HPO_4_·12 H_2_O), β-mercaptoethanol (final concentration of 2 μl/ml), intercalating fluorochrome propidium iodide (PI) and RNase IIA (both at final concentrations of 50 μg/ml). We recorded fluorescence intensities of 10,000-20,000 particles (nuclei) for three to eleven replicates (different individuals). We calculated sample/standard ratios from the means of the sample and standard fluorescence histograms, and considered only histograms with coefficients of variation <3.5% for the G0/G1 sample peak. For genome size calculation, we multiplied the sample/standard ratios with the genome size of the internal standard. For unit conversion, we used 1 pg DNA = 978 Mbp [30]. Genome size estimates are given in Data S1: 6-FCM Results.

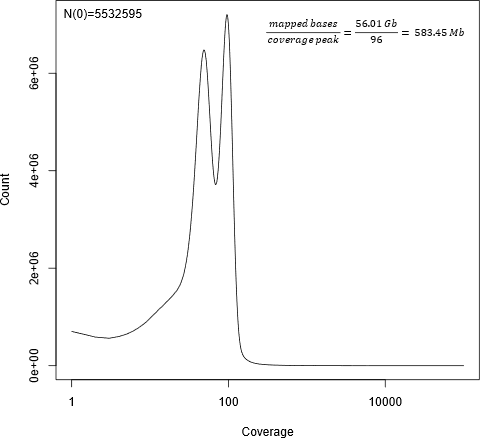

**Fig. S49. *Agapetus fuscipens*: Coverage distribution per position and genome size estimate from backmap.pl.** The x-axis is given in log-scale. For details, see Supplementary Note 7.

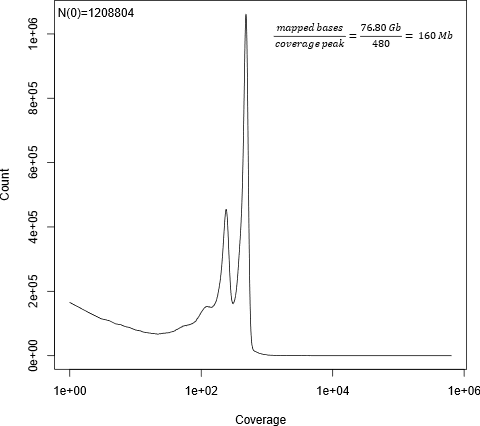

**Fig. S50: *Agraylea sexmaculata*: Coverage distribution per position and genome size estimate from backmap.pl.** The x-axis is given in log-scale. For details, see Supplementary Note 7.

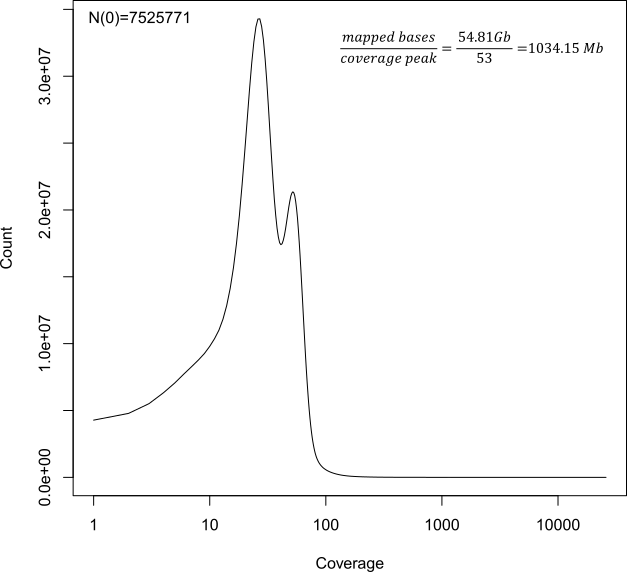

**Fig. S51. *Agrypnia vestita.* Coverage distribution per position and genome size estimate from backmap.pl.** The x-axis is given in log-scale. For details, see Supplementary Note 7.

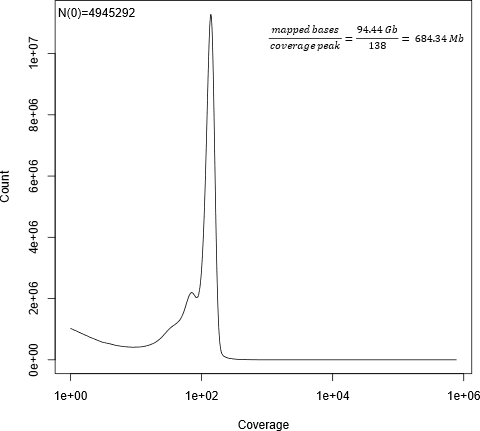

**Fig. S52. *Drusus annulatus***: **Coverage distribution per position and genome size estimate from backmap.pl.** The x-axis is given in log-scale. For details, see Supplementary Note 7.

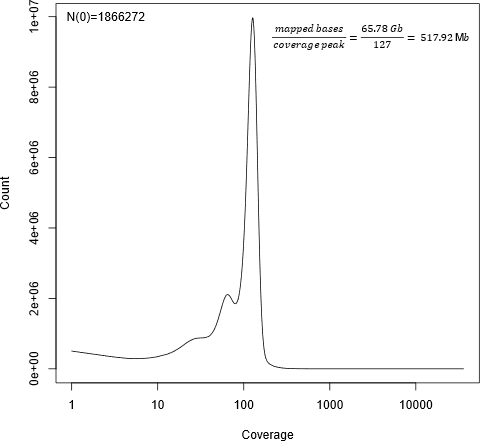

**Fig. S53. *Glossosoma conforme* G1: Coverage distribution per position and genome size estimate from backmap.pl.** The x-axis is given in log-scale. For details, see Supplementary Note 7.

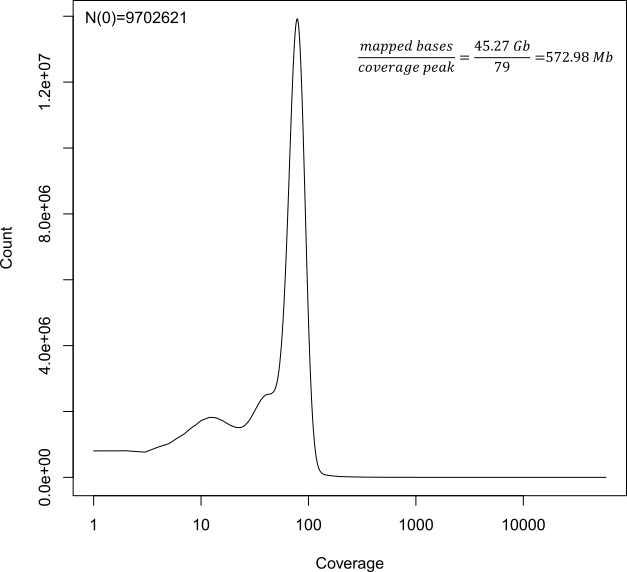

**Fig. S54. *Glossosoma conforme* Glo: Coverage distribution per position and genome size estimate from backmap.pl.** The x-axis is given in log-scale. For details, see Supplementary Note 7.

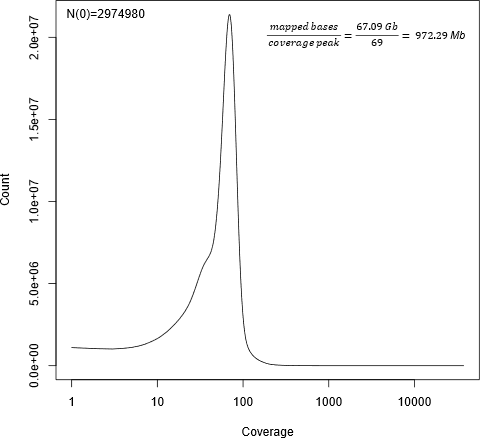

**Fig. S56. *Halesus radiatus*. Coverage distribution per position and genome size estimate from backmap.pl.** The x-axis is given in log-scale. For details, see Supplementary Note 7.

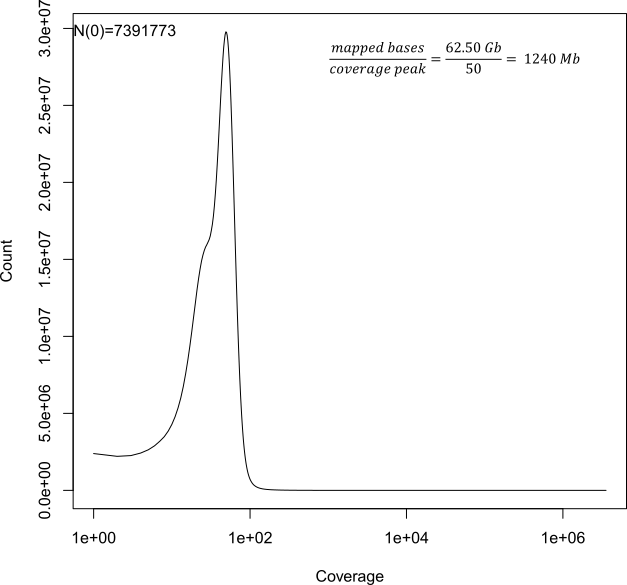

**Fig. S57. *Hesperophylax magnus*: Coverage distribution per position and genome size estimate from backmap.pl.** The x-axis is given in log-scale. For details, see Supplementary Note 7.

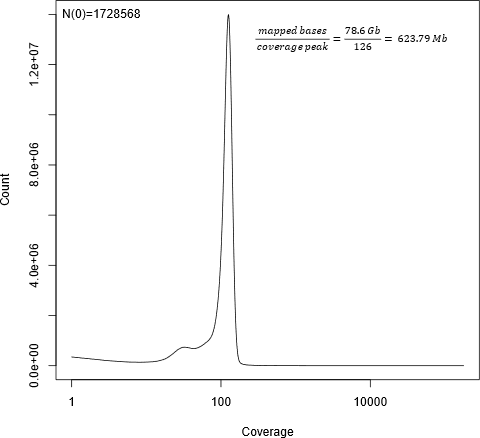

**Fig. S58. *Himalopsyche phryganeae*: Coverage distribution per position and genome size estimate from backmap.pl.** The x-axis is given in log-scale. For details, see Supplementary Note 7.

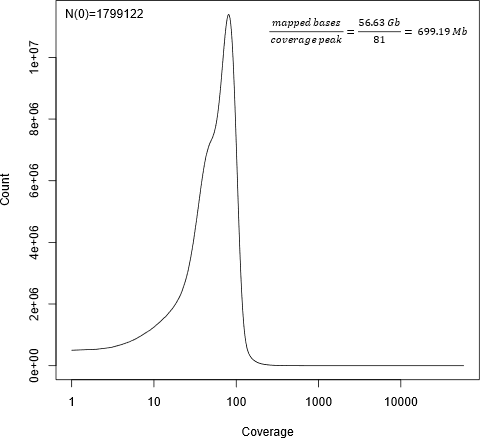

**Fig. S59. *Lepidostoma basale*: Coverage distribution per position and genome size estimate from backmap.pl.** The x-axis is given in log-scale. For details, see Supplementary Note 7.

.

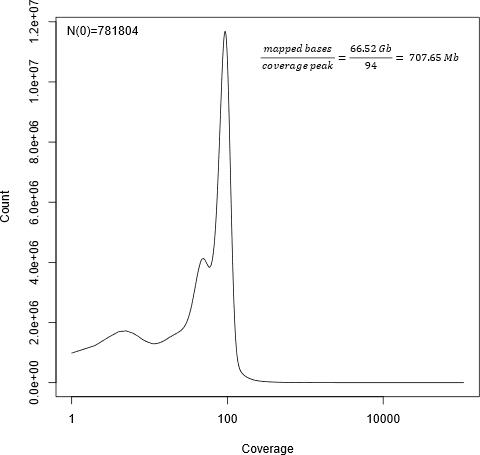

**Fig. S61. *Micrasema longulum ML1*: Coverage distribution per position and genome size estimate from backmap.pl.** The x-axis is given in log-scale. For details, see Supplementary Note 7.

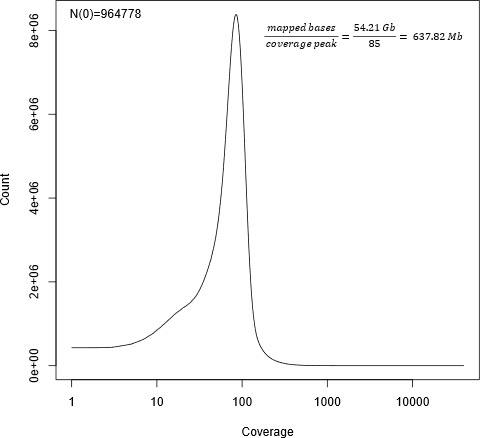

**Fig. S62. *Micrasema longulum ML3*: Coverage distribution per position and genome size estimate from backmap.pl.** The x-axis is given in log-scale. For details, see Supplementary Note 7.

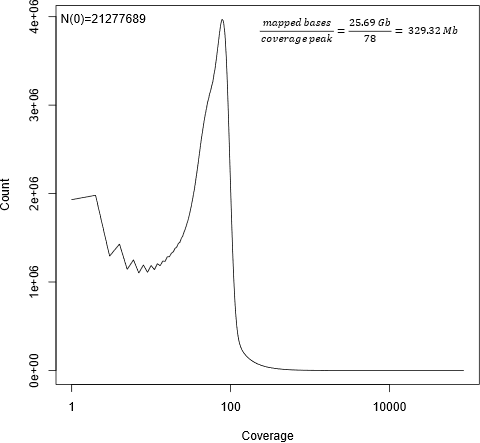

**Fig. S63. *Micrasema minimum*: Coverage distribution per position and genome size estimate from backmap.pl.** The x-axis is given in log-scale. For details, see Supplementary Note 7.

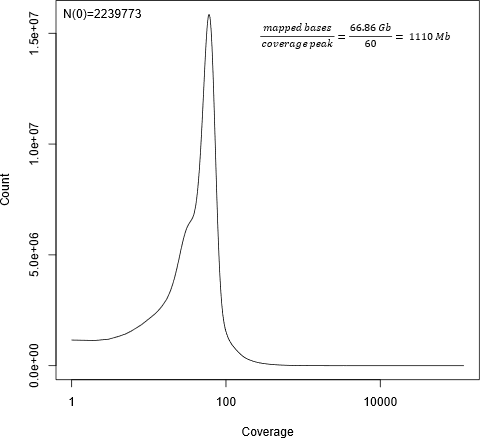

**Fig. S64. *Micropterna sequax*: Coverage distribution per position and genome size estimate from backmap.pl.** The x-axis is given in log-scale. For details, see Supplementary Note 7.

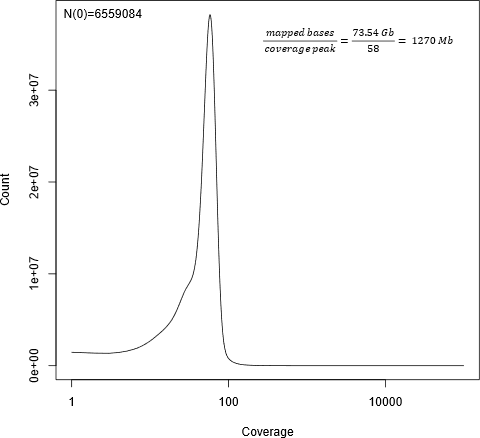

**Fig. S65. *Odontocerum albicorne*: Coverage distribution per position and genome size estimate from backmap.pl.** The x-axis is given in log-scale. For details, see Supplementary Note 7.

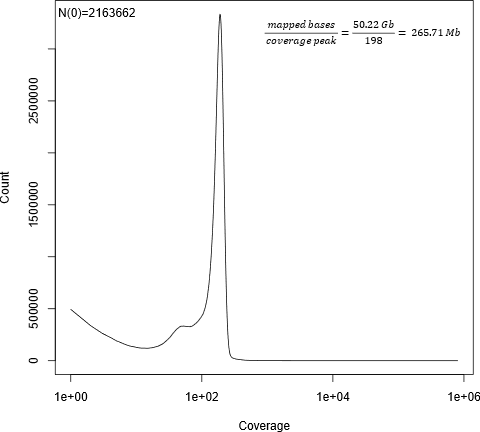

**Fig. S66. *Parapsyche elsis*: Coverage distribution per position and genome size estimate from backmap.pl.** The x-axis is given in log-scale. For details, see Supplementary Note 7.

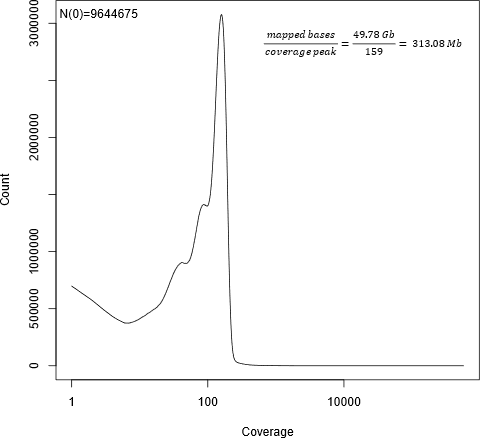

**Fig. S67. *Philopotamus ludificatus*: Coverage distribution per position and genome size estimate from backmap.pl.** The x-axis is given in log-scale. For details, see Supplementary Note 7.

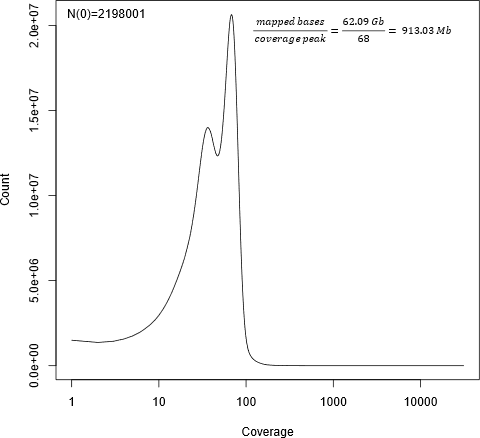

**Fig. S68. *Rhyacophila brunnea*: Coverage distribution per position and genome size estimate from backmap.pl.** The x-axis is given in log-scale. For details, see Supplementary Note 7.

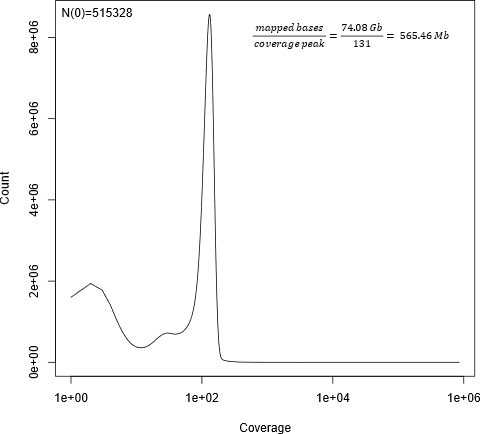

**Fig. S69. *Rhyacophila evoluta* HR1: Coverage distribution per position and genome size estimate from backmap.pl.** The x-axis is given in log-scale. For details, see Supplementary Note 7.

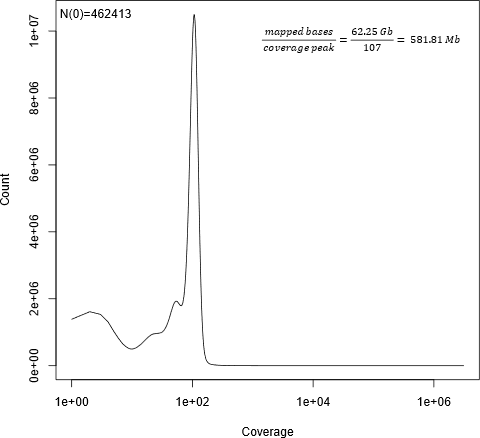

**Fig. S70. *Rhyacophila evoluta* Rss1: Coverage distribution per position and genome size estimate from backmap.pl.** The x-axis is given in log-scale. For details, see Supplementary Note 7.

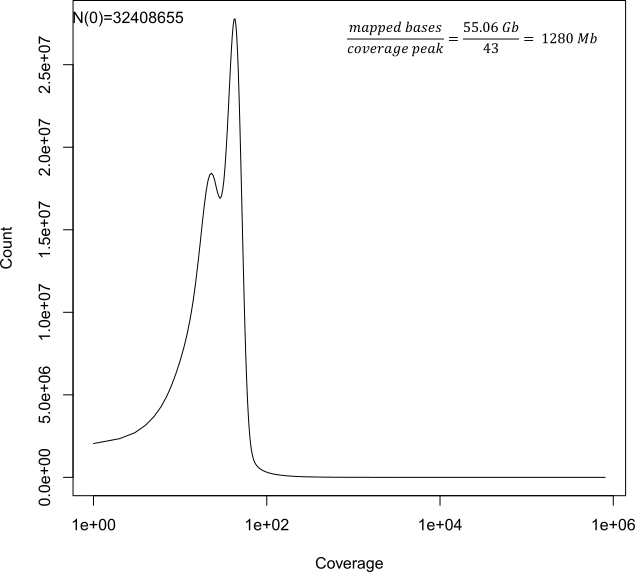

**Fig. S71. *Sericostoma sp*.: Coverage distribution per position and genome size estimate from backmap.pl.** The x-axis is given in log-scale. For details, see Supplementary Note 7.

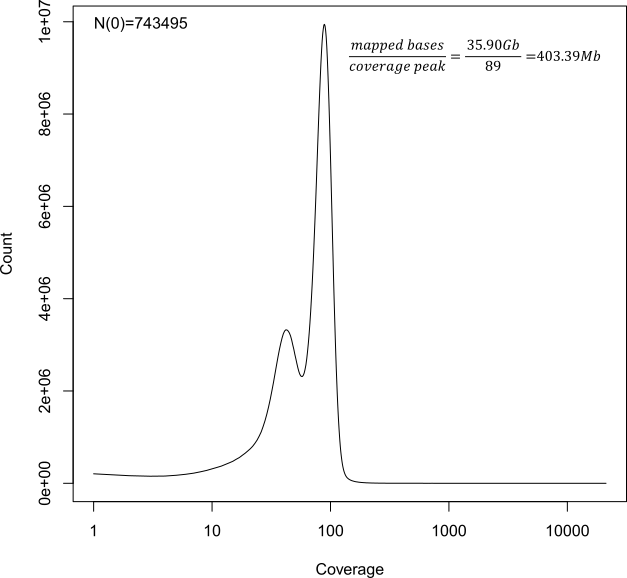

**Fig. S72. *Stenopsyche tienhuanesis*: Coverage distribution per position and genome size estimate from backmap.pl.** The x-axis is given in log-scale. For details, see Supplementary Note 7.

**Supplementary note 8: Bland-Altman-Plots**

We assessed the comparability of agreement between the three quantitative methods of genome size measurement (Genomescope2, Backmap.pl and FCM; supplementary note 7) by conducting Bland-Altman-Plots using the function BlandAltmanLeh::bland.altman.plot in ggplot2 [31] in RStudio (RStudio Team (2020). RStudio: Integrated Development for R. RStudio, PBC, Boston, MA URL http://www.rstudio.com/;) as follows:

#Bland-Altman-Plot: Genomescope2 vs. FCM

Genomescope2 <- c(222.82, 316.26, 463.24, 1103.37, 592.32, 918.70, 981.67, 642.96, 333.84, 516.62, 1134.92)

FCM <- c(260.60, 455.20, 721.79, 1616.01, 840.20, 1212.35, 1434.71, 663.60, 588.81, 651.30, 1895.64)

library(ggplot2)

BlandAltmanLeh::bland.altman.plot(Genomescope2, FCM, conf.int=.95, pch=19, main="A Bland-Altman-Plot: Genomescope2 vs. FCM")

#Bland-Altman-Plot: Genomescope2 vs. Backmap.pl

Genomescope2<- c(232.92, 222.82, 316.26, 284.67, 389.48, 463.24, 483.82, 153.52, 1103.37, 592.32, 918.70, 1060.79, 981.67, 642.96, 333.84, 542.26, 931.69, 568.37, 770.82, 516.62, 1134.92)

Backmap.pl <- c(265.71, 228.56, 364.91, 313.08, 403.98, 583.45, 545.45, 160.00, 1270.00, 684.34, 972.29, 1240.00, 1100.00, 672.74, 329.32, 699.19, 1034.15, 623.79, 913.03, 573.64, 1280.00)

library(ggplot2)

BlandAltmanLeh::bland.altman.plot(Genomescope2, Backmap.pl, conf.int=.95, pch=19, main="B Bland-Altman-Plot: Genomescope2 vs. Backmap.pl")

#Bland-Altman-Plot: Backmap.pl vs. FCM

Backmap.pl<- c(228.56, 364.91, 583.45, 1270, 684.34, 972.29, 1100, 672.74, 329.32, 573.635, 1280.00)

FCM <- c(260.60, 455.20, 721.79, 1616.01, 840.20, 1212.35, 1434.71, 663.60, 588.81, 651.30, 1895.64)

library(ggplot2)

BlandAltmanLeh::bland.altman.plot(Backmap.pl, FCM, conf.int=.95, pch=19, main="C Bland-Altman-Plot: Backmap.pl vs. FCM")

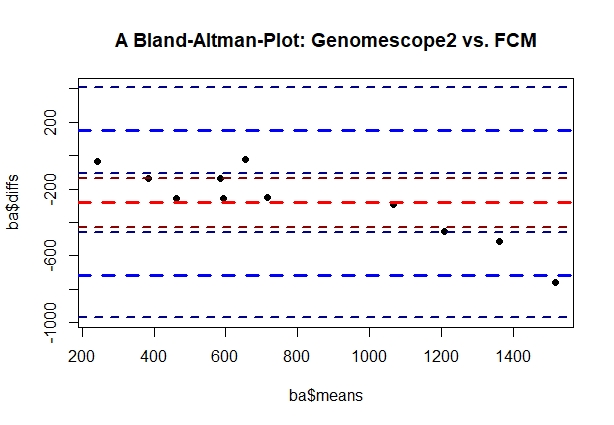

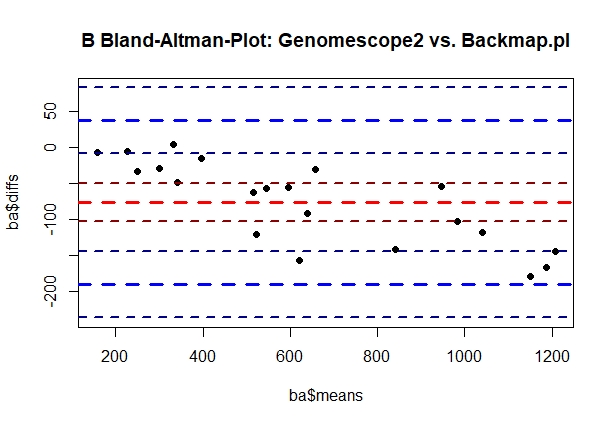

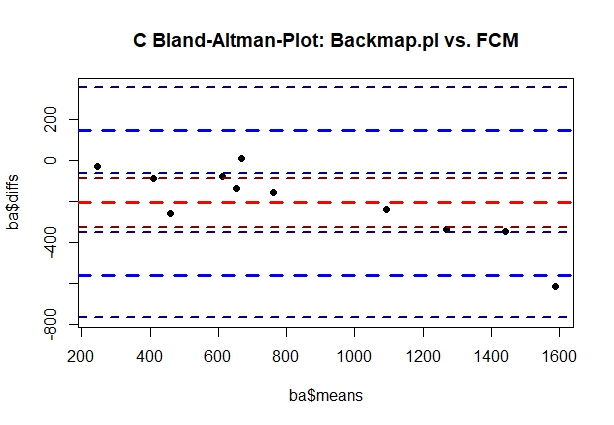

**Fig. S73. Bland-Altman-Plots** **to test the comparability of agreement between the three quantitative methods of genome size measurement (Genomescope2, Backmap.pl and FCM; supplementary note 7).** A: Genome size estimates obtained from Genomescope2 were compared to the ones obtained from Backmap.pl; B:Genome size estimates obtained from Genomescope2 were compared to the ones obtained from FCM; C: Genome size estimates obtained from Backmap.pl were compared to the ones obtained from FCM. All three methods showed agreement**.**

**Supplementary Note 9:** **Visualization of genome structure to estimate ploidy using smudgeplots**

We report that our gene-age distribution analyses do not support the hypothesis of a WGD, but given previous suggestions that WGD play a role in insects, we used smudgeplot to visualize the genome structure and estimated ploidy to rule out this possibility. We used smudgeplot with the default all algorithm. For this, we first extracted genomic kmers using reasonable coverage thresholds. These were estimated from the kmer histograms generated in step XXX with the internal smudgeplot script smudgeplot.py as follows:

L=$(smudgeplot.py cutoff kmer_*_k21.hist L)

U=$(smudgeplot.py cutoff kmer_*_k21.hist U)

echo $L $U

Then, kmers in the coverage range from L to U were extracted using jellyfish dump -c -L $L -U $U *_kmer_counts.jf -o *_jfkmers. The script smudgeplot.py hetkmers *_jfkmers -o *_kmer_pairs was used to compute the set of kmer pairs. After generating the list of kmer pair coverages, the smudgeplot was generated using the coverages of the kmer pairs (*_kmer_pairs_coverages.tsv file) using smudgeplot.py plot *_kmer_pairs_coverages.tsv -o * (where * is the abbreviation of the respective specimen we are looking at). The haploid kmer coverage was estimated directly from the data and compare to the estimation reported by Genomescope2.

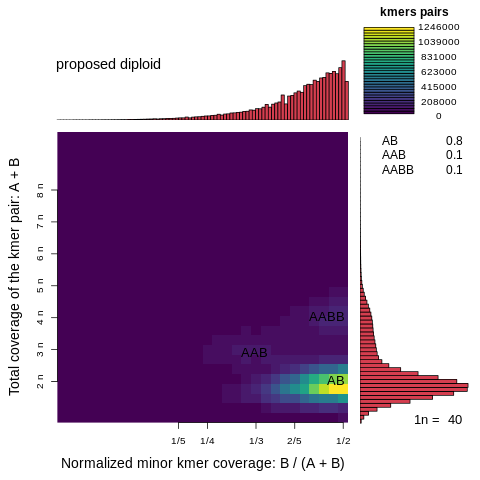

**Supplementary Figure 74: Smudgeplot for *Agapetus fuscipens* GL3 on the linear scale**

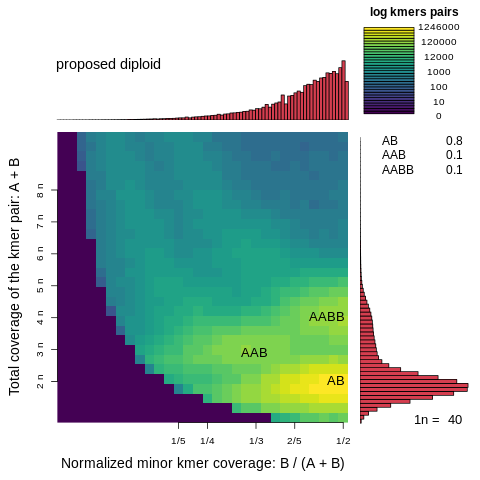

**Supplementary Figure 75: Smudgeplot for *Agapetus fuscipens* GL3 on the log scale**

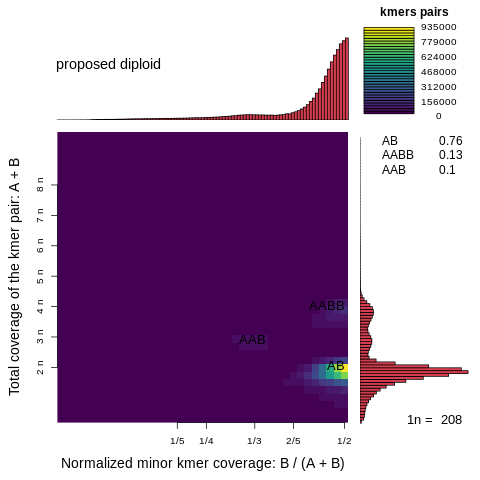

**Supplementary Figure 76: Smudgeplot for *Agraylea sexmaculata* AS19 on the linear scale**

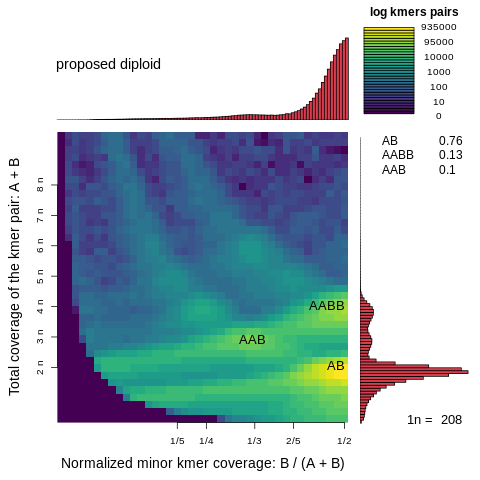

**Supplementary Figure 77: Smudgeplot for *Agraylea sexmaculata* AS19 on the log scale**

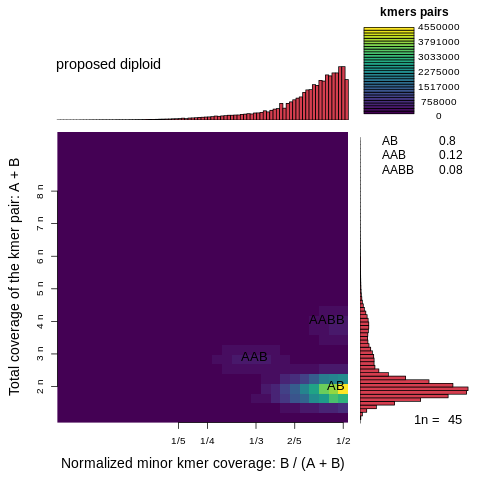

**Supplementary Figure 78: Smudgeplot for *Agrypnia vestiva* on the linear scale**

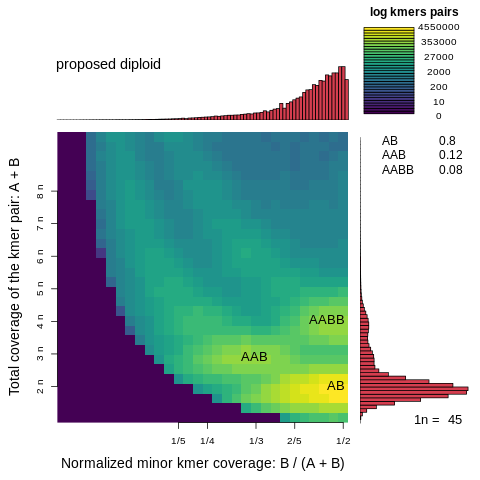

**Supplementary Figure 79: Smudgeplot for *Agrypnia vestiva* on the log scale**

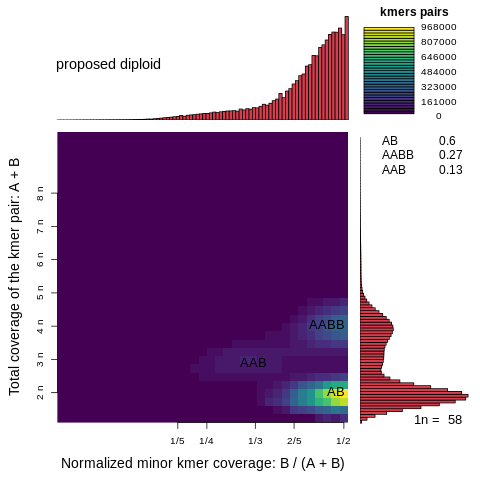

**Supplementary Figure 80: Smudgeplot for *Drusus annulatus* on the linear scale**

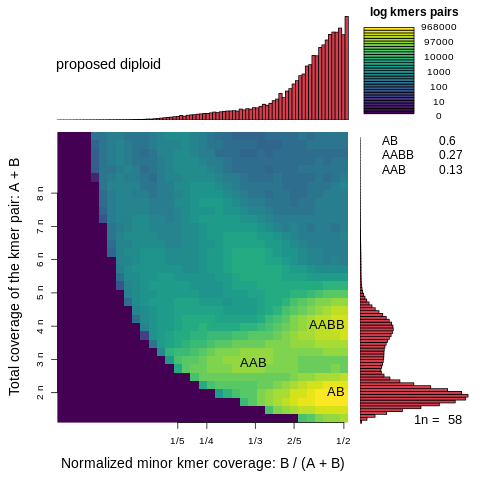

**Supplementary Figure 81: Smudgeplot for *Drusus annulatus* AC1 on the log scale**

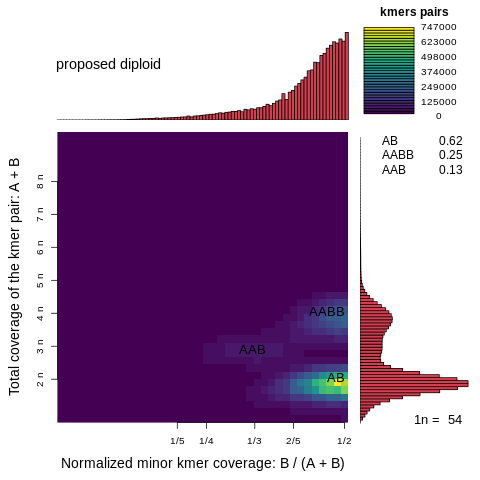

**Supplementary Figure 82: Smudgeplot for *Glossosma conforme* G1 on the linear scale**

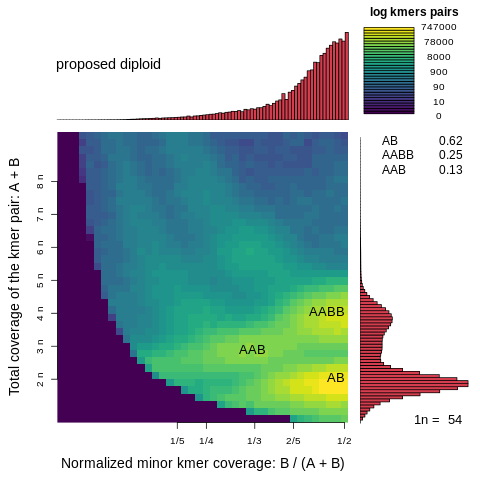

**Supplementary Figure 83: Smudgeplot for *Glossosma conforme* G1 on the log scale**

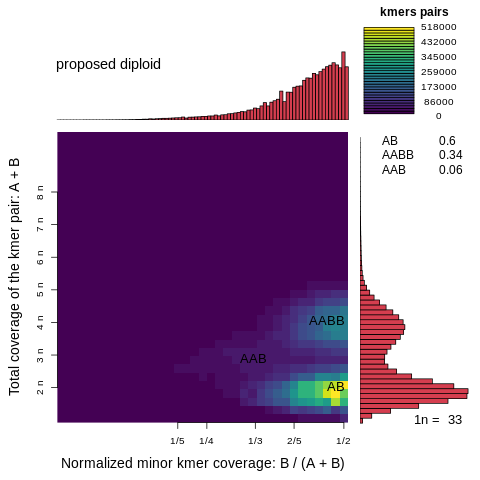

**Supplementary Figure 84: Smudgeplot for *Glossosma conforme* Glo on the linear scale**

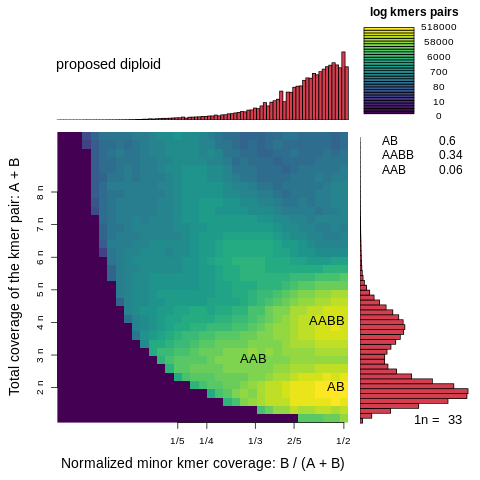

**Supplementary Figure 85: Smudgeplot for *Glossosma conforme* Glo on the log scale**

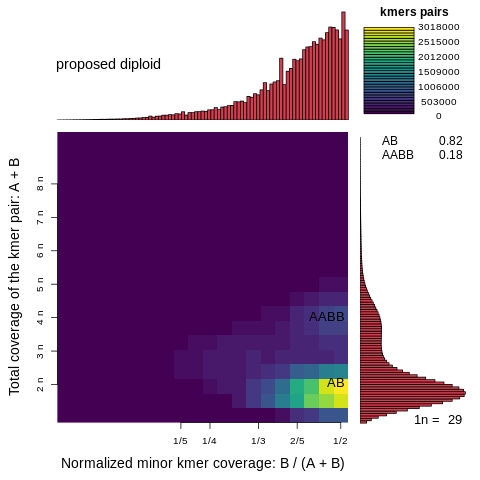

**Supplementary Figure 86: Smudgeplot for *Halesus radiatus* L2 on the linear scale**

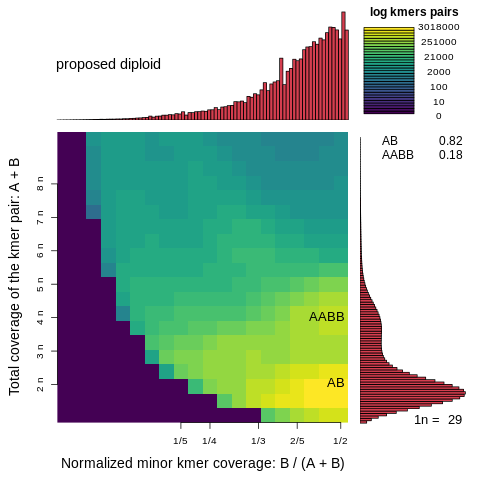

**Supplementary Figure 87: Smudgeplot for *Halesus radiatus* L2 on the log scale**

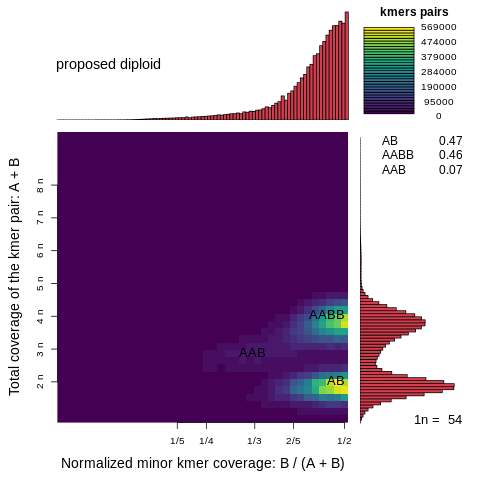

**Supplementary Figure 88: Smudgeplot for *Himalopsyche phryganeae* on the linear scale**

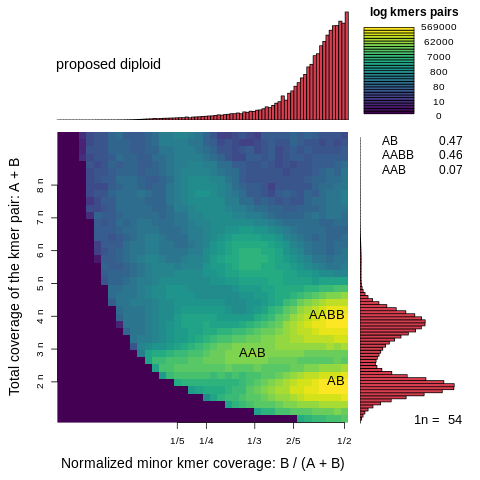

**Supplementary Figure 89: Smudgeplot for *Himalopsyche phryganeae* on the log scale**

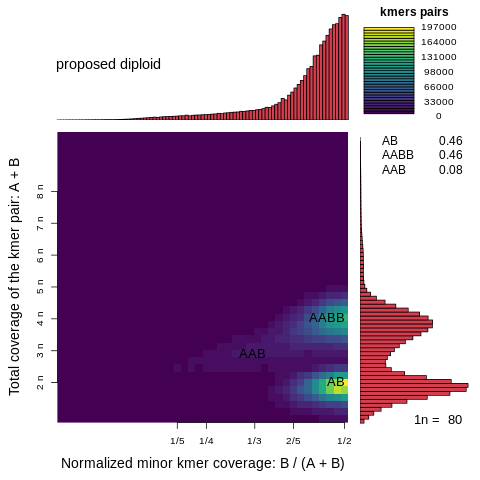

**Supplementary Figure 90: Smudgeplot for *Hesperophylax magnus* on the linear scale**

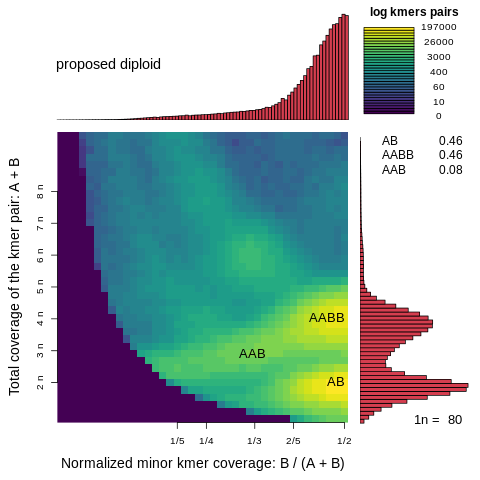

**Supplementary Figure 91: Smudgeplot for *Hesperophylax magnus* on the log scale**

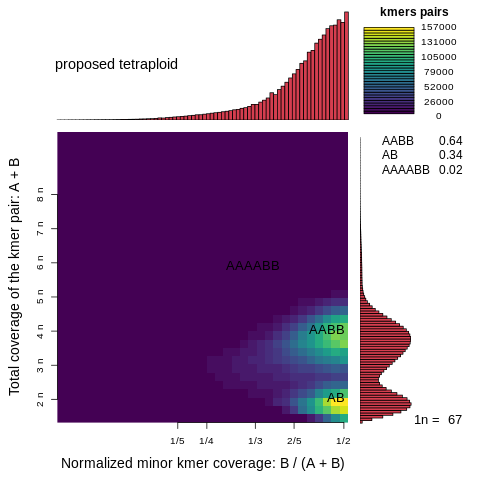

**Supplementary Figure 92: Smudgeplot for *Hydropsyche tenuis* on the linear scale**

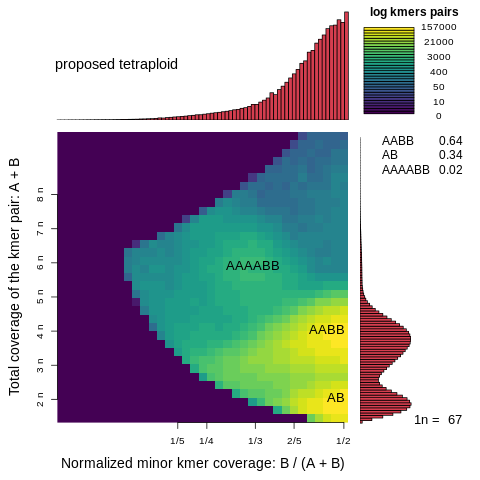

**Supplementary Figure 93: Smudgeplot for *Hydropsyche tenuis* on the log scale**

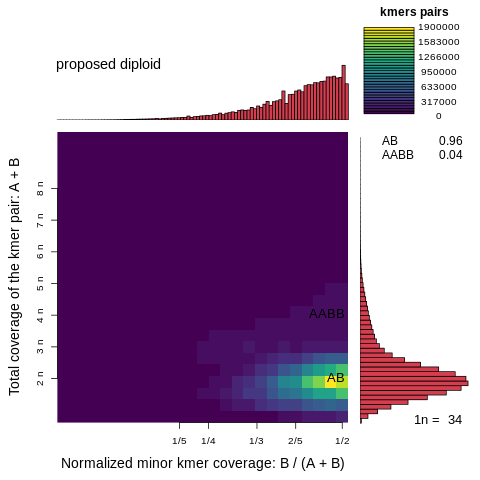

**Supplementary Figure 94: Smudgeplot for *Lepidostoma basale LB1* on the linear scale**

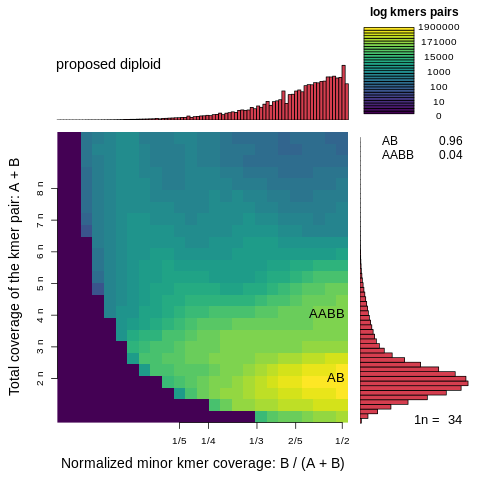

**Supplementary Figure 95: Smudgeplot for *Lepidostoma basale LB1* on the log scale**

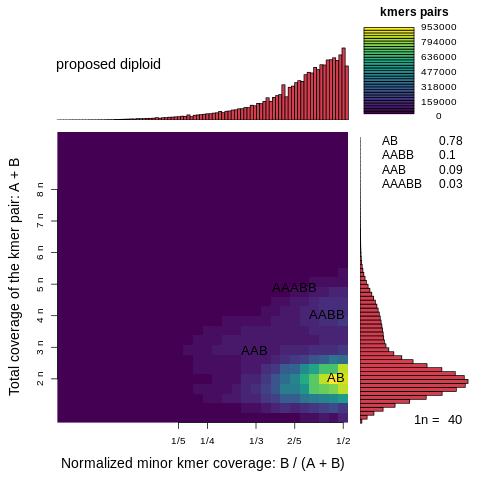

**Supplementary Figure 96: Smudgeplot for *Micrasema longulum ML3* on the linear scale**

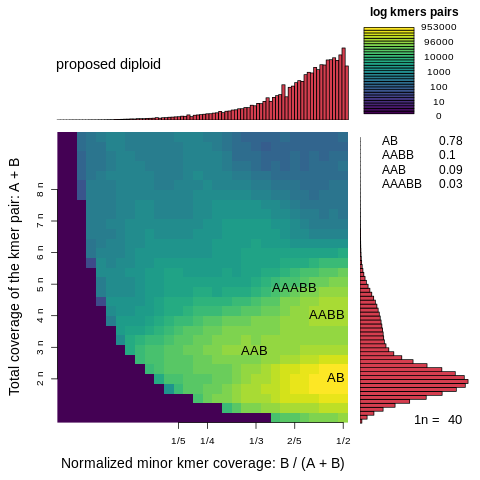

**Supplementary Figure 97: Smudgeplot for *Micrasema longulum ML3* on the log scale**

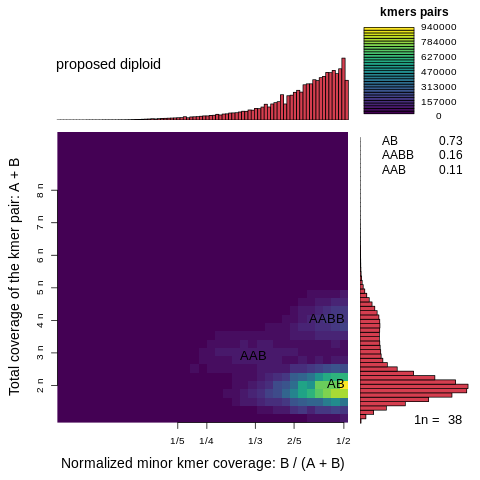

**Supplementary Figure 98: Smudgeplot for *Micrasema longulum ML1* on the linear scale**

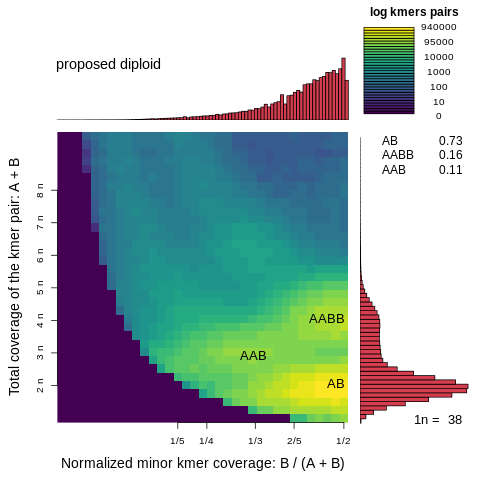

**Supplementary Figure 99: Smudgeplot for *Micrasema longulum ML1* on the log scale**

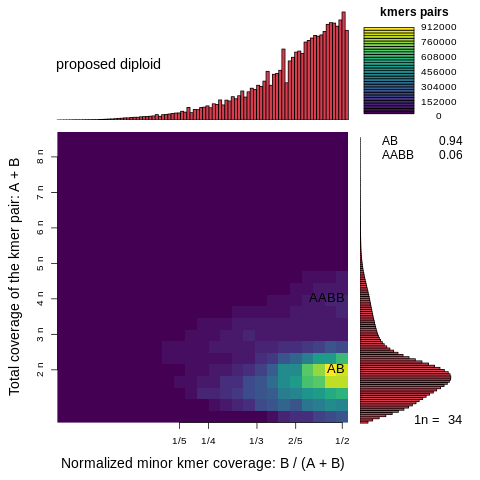

**Supplementary Figure 100: Smudgeplot for *Mcirasema minimum K05* on the linear scale**

**Supplementary Figure 101: Smudgeplot for *Micrasema minimum K05* on the log scale**

**Supplementary Figure 102: Smudgeplot for *Micropterna sequax AB8* on the linear scale**

**Supplementary Figure 103: Smudgeplot for *Micropterna sequax AB8* on the log scale**

**Supplementary Figure 104: Smudgeplot for *Odontocerum albicorne* OD1 on the linear scale**

**Supplementary Figure 105: Smudgeplot for *Odontocerum albicorne OD1* on the linear scale**

**Supplementary Figure 106: Smudgeplot for *Parapsyche elsis* on the linear scale**

**Supplementary Figure 107: Smudgeplot for *Parapsyche elsis* on the linear scale**

**Supplementary Figure 108: Smudgeplot for *Philopotamus ludificatus Ph2* on the linear scale**

**Supplementary Figure 109: Smudgeplot for *Philopotamus ludificatus Ph2* on the log scale**

**Supplementary Figure 110: Smudgeplot for *Plectrocnemia conspersa* on the linear scale**

**Supplementary Figure 111: Smudgeplot for *Plectrocnemia conspersa* on the log scale**

**Supplementary Figure 112: Smudgeplot for *Rhyacophila brunneae* on the linear scale**

**Supplementary Figure 113: Smudgeplot for *Rhyacophila brunneae* on the log scale**

**Supplementary Figure 114: Smudgeplot for *Rhyacophila evoluta HR1* on the linear scale**

**Supplementary Figure 115: Smudgeplot for *Rhyacophila evoluta HR1* on the log scale**

**Supplementary Figure 116: Smudgeplot for *Rhyacophila evoluta RSS1* on the linear scale**

**Supplementary Figure 117: Smudgeplot for *Rhyacophila evoluta RSS1* on the log scale**

**Supplementary Figure 118: Smudgeplot for *Sericostoma sp.* on the linear scale**

**Supplementary Figure 119: Smudgeplot for *Sericostoma sp.* on the log scale**

**Supplementary Figure 120: Smudgeplot for *Stenopsyche* on the linear scale**

**Supplementary Figure 121: Smudgeplot for *Stenopsyche* on the log scale**

**Supplementary Note 10:** **Repeat abundance and classification based on reference-free analyses**

Our reference-free analysis of RE abundance based on RepeatExplorer2 and dnaPipeTE (supplementary Figs. 122 & 123) showed results overall very consistent with our assembly-based approach, especially related to our major findings of LINE abundance. . This analysis also confirmed previous findings (Olsen et al. 2020) that satellite DNAs are not an abundant repeat class in trichopterans (average genome proportion = 1.56%, SD = 1.45%). Four species [(*Hydropsyche tenuis* (HT), *Hesperopylax magnus* (HM), *Limnephilus lunatus* (LL), and *Sericostoma sp.* (SS)] had genomic proportions ranging from 3.0–5.1% (whereas all trichopterans had <1% in assembly-based estimates), but overall we do not find evidence that underestimates of satellite DNAs are a substantial source of error in our assembly-based repeat estimates. However, dnaPipeTE revealed better annotations of DNA transposons (which were very uncommon in the Repeatmasker output), simple repeats, Helitrons, and some other categories. This results in a smaller proportion of unclassified repeats compared to Repeatmasker. This additional comparison suggests that dnaPipeTE may have a better internal repeat library.

**Fig. S122. Repeat abundance summary from Repeat-Explorer2**

**Assembly Percent**

**Taxon**

**Fig S123. Repeat abundance summary from dnaPipeTE**

**Fig. S124. Transposable element age distribution landscapes.** The y-axis shows TE abundance as a proportion of the genome (e.g., 1.0 = 1% of the genome). The x-axis shows sequence divergence relative to TE consensus sequences for major TE classes. TE classes with abundance skewed toward the left (i.e., low sequence divergence) are inferred to have a recent history of diversification relative to TE classes with right-skewed abundance. Plots were generated in dnaPipeTE [32]. For tip labels of the phylogenetic tree see Fig. 2.

**Supplementary Note 11:** *TE sequence association with protein-coding genes*

During the early exploration of our sequence data we had mapped reads back to BUSCO genes and noted that coverage depth profiles of some BUSCO genes showed regions of unexpected high coverage depth. The frequency of BUSCOs showing this unexpected pattern varyied widely across species. In the end, we discovered that these high coverage depth regions are TE sequences that are inserted within or adjacent to BUSCO genes, and incorrectly identified as BUSCO gene annotations by the BUSCO algorithm. Thus, we refer to these putative gene fragments discovered by BUSCO as ‘TE-associated BUSCOs’. Prior to arriving at this conclusion, we used the following methods to understand the pattern of high copy number regions we initially observed after mapping reads to BUSCO sequences. We first analyzed repeat dynamics of all BUSCO genes for all species to quantify the abundance of TE-associated BUSCOs across samples and test two alternative hypotheses that could account for the observed patterns: (1.) inflated copy-number of BUSCO gene fragments occur at sequences of repetitive elements (e.g., TEs) inserted into genes; (2.) inflated regions are BUSCO gene fragments that have proliferated throughout the genome (e.g., by hitch-hiking with TEs. This analysis also allowed us to quantify shifts in associations between TEs and genic regions across Trichoptera lineages with varying repeat abundance.

We tested for a general correlation between BUSCO repeat abundance and the abundance and major RE categories using Pearson’s correlation in R v4.0.2. To gain more detailed insight, we generated copy number profiles by mapping short reads back to all Endopterygota BUSCO genes (OrthoDB v.9) for all study species using RepeatProfiler [33]. We quantified the abundance of BUSCOs with highly covered regions by identifying BUSCO genes with coverage peak that exceeded 20X average coverage for all BUSCO genes in that species. Average coverage was calculated by averaging coverage depth of all BUSCO genes after excluding the top and bottom 15% when sorted by max coverage which eliminated 0-coverage and unexpectedly high-coverage BUSCOs from the calculation. After producing a BLAST database from each genome assembly using the application makeblastdb in ncbi-blast 2.10.0 [34] applying the parameters -dbtype nucl, -parse_seqids_blastdb, -blastdb_version5, we quantify the genomic abundance of bases in TE-associated BUSCOs by using each TE-associated BUSCO gene as a query in a BLAST (blastn) search against the genome assembly with the following settings: outfmt 6, -max_target_seqs 50000. We allowed 50K hits based on the maximum coverage of TE-associated BUSCOs observed in profiles. We used the perl script rmOutToGFF3.pl from RepeatMasker to convert the Repeatmasker OUT files from each Repeatmasker run to version 3 to gff files containing the RepeatMasker hints. We reclassified the repeats in the gff3 files with custom scripts, concatenated the two RepeatMasker runs of each species and sorted the resulting file using bedtools.

We used custom scripts to parse BLAST output, collapse hits with overlapping coordinates and extract coordinates of filtered hits. For each unique BLAST hit, we then asked whether it mapped to masked or unmasked coordinates in the assembly by cross-referencing the coordinates of BLAST hits against a gff file containing RepeatMasker annotations using the ‘intersect’ function in BEDTools. We also used custom scripts to calculate the total number of bases in filtered BLAST after subtracting the number of bases at the locus belonging to all ‘complete’ BUSCO genes, and to categorize BUSCO repeats based on their annotation status and repeat classification. We plotted the number of highly covered regions in BUSCO genes belonging to repetitive element categories (i.e., classes and subclasses) alongside plots of the relative genomic abundance of each respective category.

To test whether high coverage depth fragments are due to TE insertions or to proliferation of true BUSCO gene fragments we examined patterns of BUSCO gene structure in pairwise alignments of species in which one species has a highly covered regions (i.e., the “inflated” species) and its counterpart does not (i.e., the “reference” species). To reduce the likelihood that indels due to sequence divergence (i.e., not due to REs) would complicate the analysis we conducted this test with three pairs of closely related species: *Micrasema longulum* and *Micrasema minimum*; *Rhyacophila brunnea* and *Rhyacophila evoluta*; and *Hesperophylax magnus* and *Halesus radiatus*. We predicted that if high coverage regions were due to RE sequence insertion (i.e., Hyp1 above) that inflated species would show indels at coordinates of high coverage regions relative to reference species. If high coverage regions were due to proliferation of true BUSCO sequence (i.g., Hyp2 above) we predicted that alignments would be contiguous over the these region. Following this test, we observed that 73 of 75 randomly sampled alignments were consistently missing or non-contiguous in inflated regions in the reference sequence (Fig. 4B), suggesting that high copy number fragments consists of non-conserved sequences as would be expected from TE insertions. We further tested this prediction by taking the set of BUSCOs that only show high coverage depth in the inflated species and contrasted results of two BLAST searches followed by an intersect analysis. First, we used BLAST to compare the TE-associated BUSCO s taken from the inflated species against the genome assembly of the inflated species and quantified the number of bases in hits that mapped to annotated repeats in the assembly. We then used BLAST to compare the same set of BUSCOs taken from the reference species against the inflated species and quantified the number of bases in hits that mapped to annotated repeats in the assembly. We predicted that if TE-associated BUSCO s were due to TE insertions, that our first BLAST search would show a disproportionate number of hits that mapped to regions annotated as RE, and that when we repeated the search using orthologous BUSCOs taken from the reference sequences as queries, the number of hits belonging to annotated REs would drop substantially. We used the bedtools “intersect” function to determine the number of bases in BLAST hits that overlapped with RE annotations after merging duplicate coordinates in the BLAST output. Following this analysis, we observed that the number of BLAST hits in the genome assembly of an “inflated species” when using the BUSCO sequence from the “reference species” as a query was significantly fewer than when using the BUSCO sequence from the “inflated species” as a query (Fig, 4C, Data S2). Our intersect analysis showed that large fractions of BLAST hits for TE-associated BUSCOs (when BLASTED to their own assembly) are in fact annotated TE sequences, especially LINES and DNA transposons (Fig. 4D). We further validated the insertion of TES by mapping Nanopore reads to a subset of TE-associated BUSCOs, thus ensuring that junctions between TE-associated BUSCO regions and non-TE-associated regions were embedded in full reads and do not occur disproportionately at ends of reads as would be expected in chimeric. Taken together, these findings provide strong evidence that the TE-associated BUSCO fragments we observe in BUSCO genes are derived from TE sequences.

**Fig. S125.** **Presence and absence of** TE-associated BUSCOs**.** We conducted a binary tree search using the presence/absence matrix of TE-associated BUSCOs for each specimen, in which each of the ~2000 Endopterygota BUSCO genes were coded as ‘inflated’ or ‘non-inflated’ for 22 of the 26 study species (MS, ML2, GP, and were excluded due missing data) and analyzed as phylogenetic characters (Appendix 2). The overall lack of phylogenetic congruence with known phylogenetic relationships, especially in deep nodes, suggests high levels of homoplasy in the TE-associated BUSCO data set. Some expected relationships are recovered with high support (e.g. both specimens of *Rhyacophila evoluta* and *Glossosoma conforme* are in well-supported clades, though the latter is interrupted by the placement of *Limnephilus lunatus* which occurs on a notably long branch). Several ‘case-making’ Integripalpia, including five of the six species with the most abundant TE-associated BUSCOs (Fig. 4), also form a well-supported clade that generally mirrors phylogenetic relationships except for the placement of one basal Integripalpia [*Rhyacophila brunnea*, the basal Integripalpia with by far the most TE-associated BUSCOs (Fig. 4)] on a long branch. Finding high levels of homoplasy in the TE-associated BUSCO data set suggests the rate of evolution among TE-associated BUSCOs is fast, a conclusion which is consistent with the results of TE landscape plots (Fig. S 4).

**Fig. S126. Correlations between bases in** TE-associated **BUSCO BLAST hits and genomic abundance of repeat categories.** We investigated interactions between TEs and genic sequences by quantifying the abundance of highly covered sequences found within or adjacent to BUSCO genes. We found major expansion of TE-associated BUSCO fragments that are significantly correlated with total repeat abundance, as well as the genomic proportion of LINEs and DNA transposons.

Supplementary Note 12:

Because genome assemblies varied in contiguity, we conducted a variety of additional analyses to ensure that assembly quality is unlikely to cause artefacts during the analyses that would impact our conclusions. To test the robustness of our conclusions related to classification and abundance of transposable elements (TEs), we analyzed TE composition using both an assembly-based approach with RepeatMasker, as well as a short-read-based reference-free approaches using RepeatExplorer2 and dnapipeTE. We observe congruent patterns in the abundance of major repeat categories and unclassified repeat abundance with both methods (see Fig. S3 vs. Fig. 2 in main text). Another comparison that strengthens confidence in our TE-based analyses comes from the two species for which we included two replicates of conspecific individuals. Despite different assembly quality we observe high congruence in repeat annotation and abundance across replicate individuals (see Fig. S4 and Fig. 2 in main text: *G. conforme* 1 vs. *G.conforme* 2; *M. longulum* 1 vs. *M. longulum* 2).

Another area for which we considered the potential impact of assembly quality on our conclusion was the TE-gene association study. We quantified highly covered sequences that are incorrectly classified as BUSCOs, which we refer to as “TE-associated BUSCOs”, since theseare frequently associated with the presence of TE sequences occurring adjacent to BUSCO. One line of evidence that suggests assembly artefacts are not driving the pattern of increasing TE-associated BUSCOs with increasing assembly size is the fact that in less contiguous assemblies we found notably fewer TE-associated BUSCOs (including in replicate conspecifics with varying assembly quality), such that we could not justify including the four samples of lowest assembly quality in the full analysis. If assembly artefacts in low-quality assemblies were driving the abundance of TE-associatedBUSCOs we would have expected to find an increase of TE-associated BUSCOs with decreasing assembly quality. To further test whether assembly artefacts might be driving the abundance of TE-associated BUSCOs in more subtle ways, we mapped Nanopore reads back to an exemplary subset of TE-associatedBUSCOs to test whether junctions between TE-associated BUSCOs and non-TE-assocaited regions are embedded in full reads. For this purpose, we looked at TE-associated BUSCOs from *Hesperophylax magnus.* More specifically, we extracted all Nanopore reads >= 10 kbp sequenced of this specimen and mapped them back to the assembly of *H. magnus*. We converted the resulting sam file to a bam file with samtools and retained only reads which mapped in the alignment using samtools “view” -b -F 4. After sorting and indexing the sam file with samtools “sort” and “index” functions, the resulting bam file was imported to IGV together with the genome assembly. In IGV, we zoomed into the regions of interest (TE-associatedBUSCOs and flanking regions) which are given in Table S1. Figs. S1-S10 show A) the highly covered regions in the BUSCO gene with the flanking region and B) highly covered regions in the BUSCO gene. Junctions between highly covered regions in BUSCO genes and non-highly covered regions are embedded in full reads and do not occur disproportionately at ends of reads. This test corroborates our conclusion that the TE sequences associated with BUSCOs do in fact represent valid TE-gene associations and not simply assembly artefacts.

| BUSCO | scaffold_extract | rep_scaff_start | rep_scaff_end |
| --- | --- | --- | --- |
| EOG090R0A7C | >scaffold_868:201955-232310 | 220705 | 220814 |
| EOG090R0A26 | >scaffold_89:2596962-2639655 | 2625525 | 2625615 |
| EOG090R0AIP | >scaffold_6928:157581-209457 | 198718 | 199773 |
| EOG090R0B6S | >scaffold_6701:1555973-1588111 | 1573592 | 1574053 |
| EOG090R0BAL | >scaffold_1750:91532-114236 | 107561 | 107609 |
| EOG090R0BV8 | >scaffold_5384:414318-454283 | 429002 | 429191 |
| EOG090R0D3M | >scaffold_264:638549-690392 | 679193 | 679608 |
| EOG090R0D5K | >scaffold_2906:96588-135291 | 125160 | 125249 |
| EOG090R0DJA | >scaffold_5579:100801-141491 | 111993 | 112328 |
| EOG090R0DQF | >scaffold_396:386030-431740 | 406547 | 406614 |

**Table S1:** Ten BUSCOs of *Hesperophylax magnus*, their location in the genome and the start and end of the highly covered region.

**B**

**A**

**Fig. S127:** BUSCO EOG090R0A7C in IGV.

A: scaffold_868:201955-232310; B: scaffold_868:220705-220814

**B**

**A**

**Fig. S128:** BUSCO EOG090R0A26.

A: scaffold_89:2596962-2639655; B: scaffold_89: 2625525-2625615

**B**

**A**

**Fig. S129:** BUSCO **EOG090R0AIP** in IGV.

A: **scaffold_6928:157581-209457**; B: scaffold_6928: **198718-199773**

**B**

**A**

**Fig. S129:** BUSCO **EOG090R0AIP** in IGV.

A: **scaffold_6928:157581-209457**; B: scaffold_6928: **198718-199773**

**B**

**A**

**Fig. S131:** BUSCO **EOG090R0BAL** in IGV.

A: **scaffold_1750:91532-114236**; B: scaffold_1750: **107561-107609**

**B**

**A**

**Fig. S132:** BUSCO **EOG090R0BV8** in IGV.

A: **scaffold_5384:414318-454283**; B: scaffold_5384: **429002-429191**

**B**

**A**

**Fig. S133:** BUSCO **EOG090R0D3M** in IGV.

A: **scaffold_264:638549-690392**; B: scaffold_264: **679193-679608**

**B**

**A**

**Fig. S134:** BUSCO **EOG090R0D5K** in IGV.

A: **scaffold_2906:96588-135291**; B: scaffold_2906: **125160-125249**

**B**

**A**

**Fig. S135:** BUSCO **EOG090R0DJA** in IGV.

A: **scaffold_5579:100801-141491**; B: scaffold_5579: **111993-112328**

Fig. S136: BUSCO EOG090R0DQF in IGV.

**B**

**A**

A: scaffold_396:386030-431740; B: scaffold_396: 406547-406614

**Fig. S137. Inference of WGDs from gene age distributions KS2**

**Fig. S138. Inference of WGDs from gene age distributions KS5**

Data S1. (separate file)

Excel sheet with sampling information, assembly statistics, genome size measurements, Ks test

Data S2. (separate file)

Excel file with counts from intersect (TE-gene interactions) analysis
